# rAAV-miniBEND: A targeted vector for brain endothelial cell gene delivery and cerebrovascular malformation modeling

**DOI:** 10.1101/2025.06.10.658979

**Authors:** Jun-Liszt Li, Zhanying Bi, Xing-jun Chen, Tianyue Ming, Baoshan Qiu, Fengzhi Li, Ziyan Feng, Daosheng Ai, Tingting Zhang, Jiayu Wang, Shuai Lin, Yiping Lu, Zhanjing Wang, Juan Huang, Fei Zhao, Hu Zhao, Yilong Wang, Wenzhi Sun, Woo-ping Ge

**Author notes:** To whom correspondence should be addressed: **Woo-ping Ge, Ph.D.** (Lead contact) Wenzhi Sun, Ph.D. These authors contributed equally to this work (Jun-Liszt Li, Zhanying Bi, Xing-jun Chen).

## Abstract

Defects in brain endothelial cells (brainECs) can cause severe cerebrovascular malformations, including arteriovenous malformation (AVM) and cerebral cavernous malformation (CCM). The lack of appropriate tools for cerebrovascular disease modeling and local genetic manipulation of the brain vasculature hinders research on cerebrovascular malformations. Here we develop a recombinant adeno-associated virus (rAAV) tool miniBEND (rAAV-based <u>mini</u>-system for <u>b</u>rain <u>end</u>othelial cells, rAAV-miniBEND), which combines a minimal promoter and an optimized cis-acting element (cis-element) isolated from the mouse gene *Tek*. This system achieves gene expression specifically in mouse and rat brainECs. Using rAAV-miniBEND, we achieve high-efficiency and high-specificity gene expression in brainECs through intracranial injection at various developmental stages and through intravenous administration at all postnatal stages in mice. Furthermore, we use rAAV-miniBEND to model sporadic CCMs mediated by *MAP3K3*^I441M^ and AVMs mediated by *Braf*^V600E^. We demonstrate that somatic expression of *Braf*^V600E^ in brainECs induces an AVM phenotype, and that brainEC proliferation are important for AVM development. Thus, our rAAV-miniBEND system provides a valuable and widely applicable tool for cerebrovascular disease modeling and local or global brainEC gene delivery.

## Main

The brain, despite being only 2% of body mass, consumes 20% of the glucose and 25% of the oxygen taken in by humans^1–3^. All essential nutrients, including metabolites and lipids, are delivered to the brain through its vascular system^4^. Defects in this system, including various vascular malformations, lead to neurological diseases such as ischemic stroke, seizures, and cognitive impairments^5^. Cerebrovascular research and treatment options are limited, however, as genetic manipulation of the brain vasculature is challenging. Recently, using capsid engineering techniques, researchers have developed multiple adeno-associated virus (AAV) capsid variants^6–10^. These variants have proven to be potent and invaluable tools, exhibiting diverse transduction efficiencies and specificities for brain endothelial cells (brainEC) labeling and gene delivery in adult rodents. Notable examples include AAV2-BR1^6^, AAV-PHP.V1^10^, AAV-BI30^11^, and AAV-X1^12^. These variants, however, lack high transduction efficiency and specificity when administered through venous routes at early developmental stages or when introduced intracranially for local brainEC transduction during either developmental or adult stages. Both of these strategies are crucial for mimicking cerebrovascular malformations for mechanistic studies.

Cerebrovascular malformations, including intracranial arteriovenous malformation (AVM) and cerebral cavernous malformation (CCM) which affect ∼0.015% and 0.2% of the worldwide population, respectively, pose severe threats to the brain vasculature^13–17^. Somatic mutations in brainECs underlie these diseases, especially intracranial AVMs, which represent high-risk, life-threatening conditions^18^. Abnormalities in endothelial cells elevate the risk of hemorrhagic stroke, seizures, and other neurological deficits^19, 20^. Presently, no drugs are available for treating these brain vascular malformations^21^, which makes it challenging to study their pathogenesis because of the absence of stable disease models mimicking the somatic mutations reported in human patients. Traditional mouse genetics approaches for studying AVMs or CCMs with *KRAS*^22^ or different *CCM1/2/3*^18, 23, 24^ mutations are time-consuming and inefficient. Recombinant viral vectors, particularly recombinant adeno-associated virus (rAAVs), offer an attractive alternative for locally manipulating brainECs and replicating sporadic and local brain lesions.

Creating rAAVs with modified capsids carrying general promoters such as CAG, CMV, or EF-1α presents a challenge in achieving high cell-type specificity. Although modified capsids have shown specificity in multiple contexts^11^, using rAAV vectors with minimal promoters and cis-regulatory elements offers advantages for targeting specific cell types and modularly carrying genes of interest^25^. Because of the DNA packaging limitation of AAV vectors (i.e., <4.7 kb), researchers have developed minimal promoters for cell type–restricted expression and to save space for the gene of interest ^26, 27^. A recent study designed mini-promoters based on *Cldn5*, confirming that Ple261 is the most effective at 2963 bp, close to 3 kb^28^. Although this mini-promoter can be used in rAAV vectors, the limited space for the exogenous gene still hampers disease modeling and gene therapy applications.

Here we describe a recombinant adeno-associated virus (rAAV) system, rAAV-miniBEND (rAAV-based <u>mini</u>-system for <u>b</u>rain <u>end</u>othelial cells), that we developed for gene delivery to brainECs, enabling the modeling of cerebrovascular malformations. The rAAV-miniBEND system was created by subcloning selected segments of the *Tek* promoter and cis-acting element (cis-element), which were optimized through truncation engineering and *in vivo* characterization. This system exhibits exceptional labeling efficiency in the developing mouse brain, demonstrating higher specificity than other AAV systems in both mouse and rat brains. Notably, it allows highly efficient gene delivery within the brain vasculature when administered intracranially, a capability unmatched by other AAV systems. Our system also saves 2.5 kb of space for the gene(s) of interest. Using the rAAV-miniBEND system, we successfully generated mouse models of both sporadic and focal CCM disease by expressing the *MAP3K3*^I441M^ somatic variant in brain vasculature. Additionally, we demonstrated that the somatic variant *Braf*^V600E^ is able to induce bAVMs (brain Arteriovenous Malformations). Our findings underscore the rAAV-miniBEND system’s value, as it serves as a crucial tool for modeling diverse cerebrovascular malformations and facilitating cerebrovascular gene delivery. This system holds potential for mimicking human bAVM disease symptoms at various levels and supporting in-depth mechanistic studies.

## Results

### Development and characterization of the rAAV-miniBEND system for restricted gene expression in brainECs

To develop an rAAV vector tailored to restrict gene expression specifically to brainECs, we leveraged the endogenous gene expression regulatory system of *Tek* to incorporate the minimal promoter and cis-acting element into the vector design. *Tek* is an endothelial cell–specific gene marker with conserved gene-specific expression in both humans and rodents^29, 30^, and it results in widespread expression across endothelial cells in various vascular segments (**Supplementary** Figure 1). We collected genomic sequences of *Tek* from mouse (*Mus musculus*), rat (*Rattus norvegicus*), pig (*Sus scrofa*), cow (*Bos taurus*), marmoset (*Callithrix jacchus*), macaque (*Macaca fascicularis*), and human (*Homo sapiens*) from the UCSC Genome Browser. After generating a multiple sequence alignment, we analyzed the conserved intron 1 region, 5’ UTR region and the 2.5-kb region upstream of the transcription start site (TSS) (**Fig. 1a and Supplementary** Figure 2). The identified conserved regions were then subcloned into the rAAV vector. To systematically assess the functional properties of different truncated versions of the promoter sequences and cis-regulatory elements, we designed a standardized rAAV expression vector (**Fig. 1a**).

**Fig. 1.**
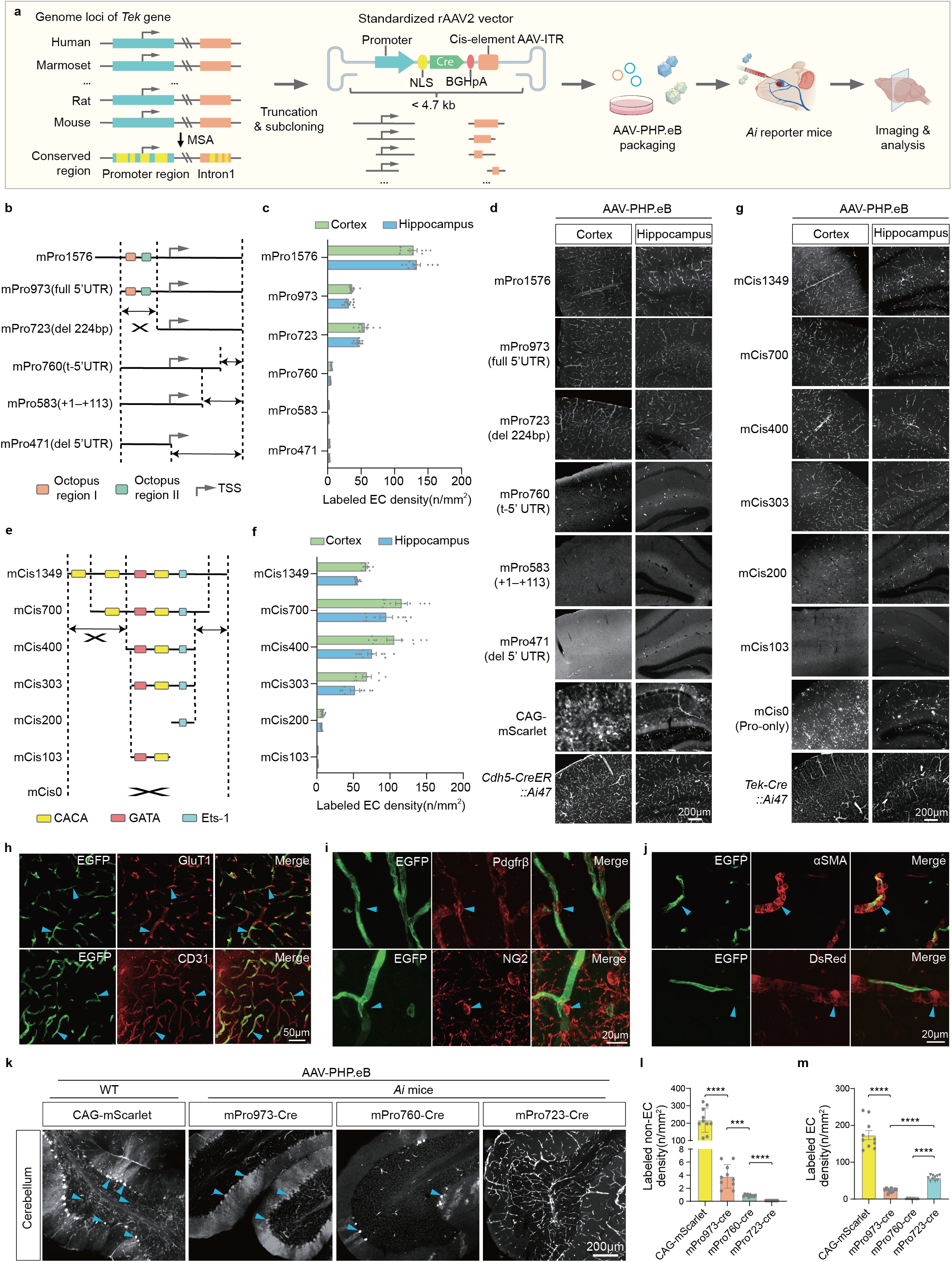
**Development of the rAAV-miniBEND system for brainEC-specific labeling.** **a**, Schematic diagram illustrating the development and characterization of the rAAV-miniBEND system. The yellow regions refer to the conserved region. The diagram was created using BioRender.com. **b,e**, Schematic diagram depicting the truncation strategy of the *Tek* promoter (b) and cis-element (e). The mCis700 was used in the recombinant AAV vectors in **b**. The mouse promoter mPro973 was used in the recombinant AAV vectors in **e**. **c,f**, Bar chart displaying the transduction efficiency mediated by various truncated versions of the promoter (c) and cis-element (f) in different brain regions. Light green bars represent the cortical region, and sky-blue bars represent the hippocampal region. Labeled EC density indicates the number of labeled endothelial cells per square millimeter per 10^11^ AAV particles, with multiple regions for cell counting and statistical analysis (n = 10 regions from 2 mice for each group in c; n = 8 regions from 2 mice for mCis303, mCis103 in f, n = 10 regions from 2 mice for mCis1349, mCis700, mCis400 in f, n =12 regions from 3 mice for mCis200 in f). The mouse promoter mPro973 was used in the recombinant AAV vectors in **f**. The mCis700 was used in the recombinant AAV vectors in **c**. **d,g**, Representative images of fluorescent protein expression (EGFP/tdTomato) mediated by various truncated versions of the promoter (d) and cis-element (g) contained in rAAV viruses shown in (b and e). **h**, Representative images of EGFP expression and co-labeling with antibodies against GluT1 and CD31. The virus rAAV-PHP.V1-mPro973-Cre-mCis1349 was administered to *Ai47* reporter mice via retro-orbital injection. Blue arrowheads, endothelial cells. **i**, Representative images of EGFP expression and co-labeling with antibodies against marker proteins of pericytes (Pdgfrβ and NG2) in brain sections from *Ai47* mice injected with the virus rAAV-PHP.eB-mPro973-Cre-mCis1349 via retro-orbital injection. Blue arrowheads, pericytes. **j**, Representative images of EGFP expression and labeling with antibodies against αSMA, a marker of vascular smooth muscle cells, in brain sections from *Ai47* or *NG2BacDsRedtg::Ai47* mice injected with AAV-PHP.eB-miniBEND-Cre. Red fluorescence indicates the DsRed signal from the *NG2BacDsRedtg* transgenic mice. Blue arrowheads, vascular smooth muscle cells. **k**, Representative images showing the transduction pattern with rAAV using AAV-PHP.eB serotype virus with different recombinant vectors in the cerebellum: CAG-mScarlet, mPro973-Cre, mPro760-Cre, and mPro723-Cre. The *Tek* cis-element used in the latter three groups was mCis700. These viruses were injected into wild-type mice or *Ai* transgenic reporter lines *Ai47* or *Ai14* as indicated. Blue arrowheads, neurons. **l,m**, The non-specific labeling ratio (l) and EC labeling efficiency (m) evaluated based on the labeling density in each transduced mouse, with each group having 10 regions selected for analysis, n = 2 mice for each group. unpaired two-sided Welch’s *t*-test, P < 0.0001 (CAG-mScarlet and mPro973-Cre); P = 0.0006 (mPro973-Cre and mPro760-Cre); P<0.0001 (mPro760-Cre and mPro723-Cre); P<0.0001 (mPro973-Cre and mPro723-Cre).Data indicate the mean ± s.e.m., \*\*\*\**p* < 0.0001, \*\*\**p* < 0.001. All image data shown are representative of two (d, k) or three (mPro723 in d, mCis200 and mCis0-Pro-only in g; h-j) independent experiments.

Using a recombinase Cre construct as the driver, which could be tested in *Ai47*^31^ or *Ai14*^32^ transgenic mice (referred to collectively as *Ai* mice) as a sensitive reporter system for Cre expression, we retro-orbitally injected *Ai* mice with the AAV virus (AAV-PHP.eB-miniBENDs-Cre) to monitor rAAV transduction and labeling specificity. *Ai14* or *Ai47* mice express robust tdTomato or GFP fluorescence following Cre-mediated recombination. The mice were housed for ∼2 weeks after virus injection and were subsequently examined for tdTomato or GFP expression (**Fig. 1a**).

To identify the crucial regulatory sequences within the promoter and cis-element, we conducted a series of truncation experiments for both sequences (**Supporting Materials**), which were subsequently subcloned into our rAAV-Cre backbone (**Fig. 1b–g**). Upon *in vivo* transduction and whole-brain examination of GFP expression, we observed that truncation of the 5’ proximal region of the promoter had no discernible effect on promoter activity. However, truncation in the 5’ untranslated region (UTR) significantly impaired the promoter activity (**Fig. 1c,d**). Notably, mPro723, a truncated version of mPro973 with a deletion of 224 bp from the 5’ proximal region of the promoter, including two crucial motifs for transcription factor binding (Octopus region I and Octopus region II)^33^, exhibited effective promoter activity, even displaying a slight improvement as compared to mPro973 (**Fig. 1c,d**).

To identify the crucial regulatory sequences within the cis-regulatory element, we constructed rAAV vectors containing six truncated versions of the cis-regulatory elements from the mouse genome (mCis): mCis1349, mCis700, mCis400, mCis303, mCis200, and mCis103 (**Fig. 1e, also see Supporting Materials** with sequences), and all of these rAAV vectors were packaged into AAV viral particles with the PHP.eB capsid for *in vivo* testing. We selected the AAV-PHP.eB capsid because of its high transduction efficiency in multiple cell types in the brain, including neurons, endothelial cells, and astrocytes. Additionally, the AAV-PHP.eB capsid can be delivered via intravenous injection and transduce cells across the entire brain, making it an attractive option for gene delivery throughout the brain. Among these versions, mCis1349 encompassed the most regulatory elements, including CACA repeat motifs, GATA motifs, and binding motifs for the Ets-1 transcription factor^34–36^. mCis700, mCis400, and mCis303 effectively preserved the overall transcriptional activity when compared to mCis1349.

However, mCis200 and mCis103 exhibited robust suppression of transcriptional activity (**Fig. 1f,g**), suggesting the loss of essential regulatory sequences necessary for normal transcriptional initiation in the promoter region. Specifically, mCis200 partially retained transcriptional activity, indicating the partial inhibitory effect of the Ets-1 motif on transcription in the promoter region. In contrast, mCis103 almost entirely lost all transcriptional activity, highlighting the stronger inhibitory effects of the GATA and CACA repeat motifs on transcriptional initiation in the promoter region. Notably, the expression of the Cre gene in the mCis0 group (Pro-only, i.e., only the promoter sequence was present without the cis-regulatory element) was not restricted to brainECs but was broadly expressed in neurons and astrocytes (**Fig. 1e–g**). Therefore, we conclude that the cis-acting elements within the intron 1 region of *Tek* result in transcriptional inhibition in non-EC cells: they restrict the expression of exogenous genes exclusively to brainECs.

In addition to testing the mouse *Tek* promoter (mPro) and cis-element (mCis), we also conducted experiments involving the truncated version of the human *TEK* promoter and cis-element (hPro and hCis, respectively, see **Supporting Materials** for all sequences) and evaluated their activity. hPro1612 and hPro762 also exhibited high specificity when labeling mouse brainECs (**Supplementary** Figure 3). To explore the potential of using cis-regulatory elements from *Tek* genes of other species in combination with hPro to achieve specific gene expression in brainECs, we synthesized and cloned cis-regulatory elements corresponding to the intron 1 region of *Tek* from various species, including rats, marmosets, and pigs. Each of these elements was individually integrated into our designed recombinant vector and packaged into the AAV-PHP.eB virus for *in vivo* experiments in mice. The results unequivocally demonstrated that cis-regulatory elements from different species could function effectively with the human promoter, enabling specific gene expression in mouse brainECs (**Supplementary** Figure 3).

To evaluate the cell-type specificity of labeling by the rAAV-miniBEND system, we conducted staining using antibodies against GluT1 and CD31 for endothelial cell labeling. The cells expressing GFP exhibited vessel-like morphology, demonstrating very high specificity and co-localization with the brain endothelial cell markers GluT1 (96.38%, n = 266/276 cells) and CD31 (100%, n = 286/286 cells; **Fig. 1h**). This result suggests that the rAAV-miniBEND system specifically enable the expression of GFP in brainECs. Moreover, we used antibodies against Pdgfrβ and NG2 to stain for pericytes and anti-αSMA to stain for smooth-muscle cells. These two mural cell types showed no co-localization signals with GFP^+^ cells (**Fig. 1i-j**). We further stained the brain section using antibodies against NeuN, Iba-1, and GFAP for the labeling of neurons (**Supplementary** Figure 4), astrocytes (**Supplementary** Figure 5), and microglia (**Supplementary** Figure 6), respectively. We did not detect co-localization of these cells with EGFP expression.

In addition, injection of AAV-miniBEND-Cre and AAV-miniBEND-EGFP viruses in the mouse strains *NG2DsRedBAC::Ai47* (Fig. 1j) and *Pdgfr*β*-Cre::Ai14* (**Supplementary** Figure 7), in which both smooth-muscle cells and pericytes express the red fluorescent proteins DsRed and tdTomato, respectively. There were no pericytes or smooth-muscle cells showing EGFP expression in these transgenic strains (**Fig. 1j, Supplementary** Figure 7), which further confirmed the specificity of the rAAV-miniBEND system.

To further examine the cell-type specificity of labeling by the rAAV-miniBEND system, we examined GFP and tdTomato fluorescence signals across all brain regions in *Ai14 or Ai47* mice after intravenous administration of rAAV-miniBEND-EGFP or rAAV-miniBEND-Cre viruses. The rAAV-miniBEND system in combination with the AAV-PHP.eB capsid resulted in a high level of brainEC labeling in nearly all brain regions except the cerebellum (Fig.1k, **Supplementary** Figure 8a**, b**). In comparison with the AAV-PHP.eB-CAG-mScarlet group, although the truncated version mPro973 resulted in a notable reduction in non-specific labeling with the AAV-PHP.eB capsid, it still labeled Purkinje cells within the cerebellar region to some extent (**Fig. 1k**). Notably, a reduction in labeling of Purkinje cells was observed in both the mPro760-Cre and mPro723-Cre groups, with mPro723-Cre demonstrating almost undetectable labeling in neurons and glial cells. Our results suggest that progressive truncation of the 5’ end proximal region of the mouse-derived *Tek* promoter effectively minimized non-specific labeling of neurons in the brain, with the optimized truncation version, mPro723, displaying the greatest cell-type specificity (Fig. 1l, m). Ple261 is a mini-promoter modified from the CLDN5 gene for integration into AAV for brain endothelial cell labeling^28, 37^ .We injected both AAV-PHP.eB-miniBEND-Cre and AAV-Ple261-iCre-pA into Ai reporter mouse lines. We found very specific labeling of brain endothelial cells with AAV-PhP.eB-miniBEND-Cre, but a lot of neurons and glial cells were labeled in the brain after infection with AAV-PHP.eB-Ple261-iCre (although we also found brain endothelial cell labeling, **Supplementary** Figure 9). Our labeling results were consistent with the previous report on AAV-PHP.eB-Ple261-iCre^10^. Through comparison with Ple261, our results strongly show the specificity of the miniBEND system for brain endothelial cell labeling.

To test the efficacy of the rAAV-miniBEND system in targeting retinal vascular endothelial cells, we performed intravenous injections of AAV-PHP.eB-mPro973-miniBEND-Cre viruses into *NG2DsRedBAC::Ai47* mice. EGFP expression mediated by AAV-miniBEND-Cre was observed in retinal endothelial cells and in the capillary, arterial, and venous segments of the retina (**Supplementary** Figure 10a), indicating that the rAAV-miniBEND system can be used to target retinal endothelial cells of all segments. It has been reported that AAV-BR1 minimally infects endothelial cells when injected intravitreally^38^. Given that this administration route is valuable for clinical treatment of retinal diseases, we performed intravitreal injections of AAV-PHP.eB-miniBEND-Cre and AAV-9P13-miniBEND-Cre viruses. The AAV-PHP.eB capsid, in combination with the rAAV-miniBEND system, enabled higher efficacy in the labeling of retinal endothelial cells as compared with the AAV-9P13 capsid when the viruses were injected into *Ai* reporter mice (**Supplementary** Figure 10b-c). Additionally, we conducted tests using AAV-BI30 to target retinal vascular endothelial cells in rats. Remarkably, the rAAV-miniBEND system, integrated in AAV-BI30, demonstrated higher transduction efficiency and specificity when compared to the Ef1α-EGFP-based AAV-BI30 group. The latter exhibited a significant proportion of non-specific neuronal labeling (**Supplementary** Figure 10d). Our findings suggest that the rAAV-miniBEND system effectively maintains gene delivery specificity to retinal endothelial cells in rodent species.

### rAAV-miniBEND enables gene delivery to brainECs of developing mice and rats

Achieving cell type–specific genetic manipulation in the developing brain is critical for mimicking certain cerebral malformations^18, 23, 24, 39^. To explore the capability of the rAAV-miniBEND system with respect to gene delivery to brainECs at different developmental stages, we tested its efficacy in embryonic (intracerebroventricular [i.c.v.] injection), early postnatal (lateral ventricle injection), and adolescent (retro-orbital and local injection) mice. The expression patterns of EGFP in *Ai47* mice following retro-orbital injection of AAV-BI30-Ef1α-Cre and AAV-BR1-Ef1α-Cre viruses were examined. We observed that AAV-BI30 and AAV-BR1 are valuable tools for labeling adult brain endothelial cells, as previously reported. Both viruses exhibit high transduction efficiency in endothelial cells of the adult brain (**Supplementary** Figure 11), but some neurons were also labeled in the AAV-BR1-Ef1α-Cre group (**Supplementary** Figure 11). Further, using AAV-BR1, we observed extensive non-brainEC labeling including astrocytes and neurons when the viruses were injected into adolescent mice (171.18 ±11.36 cells/mm^2^, **Fig. 2a–d**). In these experiments, we administered two types of AAV particles, AAV-BR1-Ef1 α-EGFP and AAV-BR1-miniBEND-Cre, to wild-type and Ai reporter line mice, respectively (**Fig. 2a–d**).

**Fig. 2.**
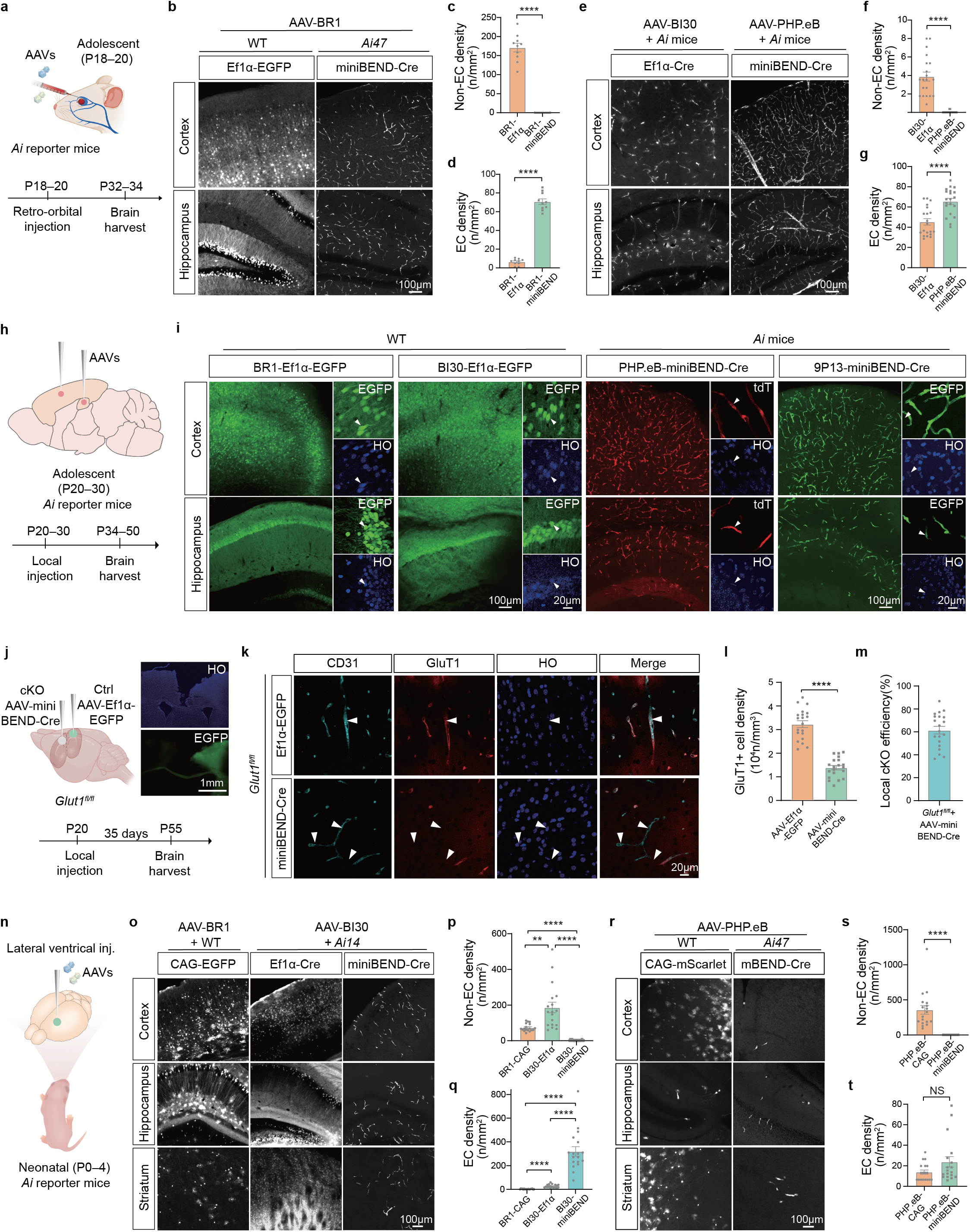
Characterization of the rAAV-miniBEND system for gene delivery to brainECs. **a**, Schematic diagram of the rAAV-miniBEND system injection strategy for adolescent mice. **b**, Representative images of EGFP expression illustrating the transduction efficiency of AAV-BR1 viruses carrying different recombinant vectors in various brain regions. Ef1α-EGFP, AAV-BR1-Ef1α-EGFP; miniBEND-Cre, AAV-BR1-miniBEND (mPro1576-mCis700)-Cre. **c,d**, Density of non-endothelial cells (non-ECs) and endothelial cells (ECs) labeled in each group as in (b), evaluated on the basis of labeling density (number of labeled non-EC or EC cells per square millimeter per 10^11^ AAV particles. i.e., The final cell density is presented per 10¹¹ AAV particles injected intravenously in mice), with each group having 10 regions selected for analysis, n = 2 mice for each group, unpaired two-sided Welch’s *t*-test, P < 0.0001, \*\*\*\**p* < 0.0001. **e**, Representative images displaying the transduction efficiency of AAV-BI30 virus carrying different recombinant vectors and AAV-PHP.eB virus in each brain region. Ef1α-Cre, AAV-BI30-Ef1α-Cre; miniBEND-Cre, AAV-PHP.eB-miniBEND (mPro723-mCis700)-Cre. **f,g**, Density of non-ECs and ECs labeled in each group as in (e), evaluated based on the labeling density as in (b), with each group having 10 regions selected for analysis (n = 10 regions from 2 mice), unpaired two-sided Welch’s *t*-test, P < 0.0001, \*\*\*\**p* < 0.0001. **h**, Schematic diagram showing local injection of the rAAV-miniBEND system into the brains of adolescent mice. **i**, Representative images showing the transduction efficiency of each group of viruses in the cerebral cortex versus hippocampus. BR1-Ef1α-EGFP, AAV-BR1-Ef1α-EGFP; BI30-Ef1α-EGFP, AAV-BI30-Ef1α-EGFP; PHP.eB-miniBEND-Cre, AAV-PHP.eB-mPro723-Cre-mCis700; 9P13-miniBEND-Cre, AAV-9P13-mPro723-Cre-mCis700. White arrowheads, neurons. **j**, Schematic of the experimental design for local injection of rAAV-miniBEND for conditional gene knockout (cKO) in *Glut1^fl/fl^* mice. Stereotaxic injection of the virus was at P20, and brain samples were collected 35 days later. EGFP fluorescence was showed in the right hemisphere as the control (insets). Blue, nucleus staining by Hoechst33342. **k**, Representative images displaying the labeling for GluT1 and CD31 in the *Glut1^fl/fl^* mice after local injection of AAV-PHP.eB-miniBEND (mPro723-mCis700)-Cre (cKO) and AAV-Ef1α-EGFP (Ctrl) in the left and right hemisphere, respectively. White arrowheads, brainECs. **l**, Cell density of GluT1^+^ brainECs in the cKO and Ctrl brain regions shown in (k). Cell density: the number of GluT1^+^ EC cells per mm² area, n = 20 regions from 4 mice for each group, unpaired two-sided Welch’s *t*-test, P < 0.0001, \*\*\*\**p* < 0.0001. **m**, Efficiency of rAAV-miniBEND (AAV-PHP.eB-mPro723-mCis700)-Cre for cKO (i.e., percentage of GluT1^−^ cells among CD31^+^ cells) after local injection of *Glut1^fl/fl^* transgenic mice (n = 20 regions from 4 mice). **n**, Schematic of the rAAV-miniBEND viruses injected into the lateral ventricle of neonatal mouse brains. **o**, Representative images showing the labeling of brainECs in different brain regions after injection of AAV-BR1-CAG-EGFP, AAV-BI30-Ef1α-Cre, and AAV-BI30-miniBEND (mPro723-mCis700)-Cre viruses into the lateral ventricle of early wild-type (WT) postnatal mice (P0–4) as indicated. AAV-BI30-miniBEND-Cre, AAV-BI30-miniBEND (mPro723-mCis700)-Cre. **p,q**, Density of labeled non-ECs (p) and brainECs (q) for each group in (o), evaluated based on the labeling density (the number of labeled non-EC or EC cells per square millimeter per 10^10^ AAV particles, from slice thickness, 50μm), with each group having 18 regions analyzed (n = 18 regions from 3 mice), unpaired two-sided Welch’s t-test, P = 0.0013 (in p, BR1-CAG and BI30-Ef1α), P < 0.0001 (in both p and q, BR1-CAG and BI30-miniBEND), P < 0.0001 (in both p and q, BI30-Ef1α and BI30-miniBEND), P < 0.0001 (in q, BR1-CAG and BI30-Ef1α), \*\**p* <0.001, \*\*\*\**p* < 0.0001. **r**, Representative images displaying the labeling of brain ECs in multiple brain regions after injection of AAV viruses into the lateral ventricle of WT or *Ai* neonatal mice as indicated. CAG-mScarlet, AAV-PHP.eB-CAG-mScarlet; mBEND-Cre, AAV-PHP.eB-miniBEND (mPro1576/mPro973-mCis700)-Cre. **s,t**, Density of labeled non-ECs (**s**) and ECs (**t**) for each group in (r). Labeling density was determined as in (p,q), with each group having 18 regions analyzed (n = 18 regions from 3 mice for each group). Data indicate the mean ± s.e.m. along with individual data points; unpaired two-sided Welch’s t-test, P < 0.0001 (in s, PHP.eB-CAG and PHP.eB-miniBEND), P = 0.0869 (in t, PHPeB-CAG and PHP.eB-miniBEND), \*\*\*\**p* < 0.0001; NS, not significant; HO, nuclear staining with Hoechst 33342. All image data shown are representative of two (i) or four (j) independent experiments. Diagrams in (a), (h), (j), (n) were created using BioRender.com.

We selected the Ai reporter line (*Ai14* or *Ai47*) for AAV-BR1-miniBEND-Cre administration to enhance sensitivity in detecting leakage of the miniBEND system, as EGFP expression may not be sufficiently bright to detect expression in non-endothelial cells. Although the AAV-BI30 serotype exhibited significantly improved performance in endothelial cell labeling, neuronal labeling was still present in adolescent mice (3.89 ± 0.48 cells/mm^2^, **Fig. 2e–f**). However, employing our rAAV-miniBEND system (AAV-PHP.eB-miniBEND) yielded a significant improvement in the specificity of endothelial cell labeling. When AAV-PHP.eB-miniBEND was compared with AAV-BI30, non-brainEC labeling was markedly reduced in the former, and we also observed a modest increase in the labeling efficiency of brainECs with AAV-PHP.eB-miniBEND (**Fig. 2e–g**).

To further examine the percentage of infected cells in developing brain (*Ai14* or *Ai47*, P5–10) after intravenous injection of AAV-PHP.eB-miniBEND-Cre and AAV-BI30-Cre. We then examined the percentages of CD31^+^ cells among all tdTomato (or EGFP) positive cells in the left hemisphere using flow cytometry. Our results showed that ∼95% of tdTomato^+^ cells in *Ai14* (or *Ai47*) brains infected with AAV-PHP.eB-miniBEND-Cre were CD31^+^, compared to ∼80% for AAV-BI30-Cre. Imaging from brain sections in the right hemispheres of the same mice confirmed that AAV-PHP.eB-miniBEND-Cre achieved nearly 100% CD31^+^ cell targeting (n=649 red cells), compared to 75.1% (n=1,268 red cells) with AAV-BI30-CAG-Cre (**Supplementary** Figure 12). This provides strong evidence of AAV-PHP.eB-miniBEND-Cre’s high specificity for endothelial cells.

To assess the transduction specificity of the rAAV**-**miniBEND system in brainECs, we integrated the miniBEND expression elements containing Cre with other rAAV vectors featuring conventional AAV capsid serotypes and administered AAV via retro-orbital injection in *Ai* reporter mice. We tested four capsid serotypes known for labeling non-brainEC cell types such as neurons: AAV-DJ/8^40^, AAV1-rh.10^41^, AAV-9P13^42^, and AAV-9P36^42^. Notably, The miniBEND promoter and regulatory element were compatible with multiple AAV capsid serotypes that exhibit tropism for brainECs. This integration significantly enhanced the specificity of the brainEC labeling outcomes (**Supplementary** Figure 13).

Acknowledging the significant challenge of off-target effects in clinical applications of AAV, particularly in the liver^43^, we examined off-target labeling in peripheral organs, specifically in the liver and lung, using the rAAV-miniBEND system. Using *Tek-Cre::Ai47* transgenic mice as a reference for peripheral vascular labeling, we observed the widespread distribution of fluorescence in peripheral vascular endothelial cells after intravenous injection of the AAV-PHP.eB-Ef1α-Cre or AAV-BI30-Ef1α-Cre in *Ai* reporter mice (**Supplementary** Figure 14 **and Supplementary** Figure 15). However, almost no fluorescence was detected in the liver, kidney, heart, intestine from mice injected with an AAV-PHP.eB-miniBEND-Cre virus (**Supplementary** Figure 15). Furthermore, only a very few large blood vessels in the lung tissue, few small blood vessels in the spleen tissue and stomach tissue were associated with fluorescence, indicating limited Cre gene expression compared to the other three control groups (**Supplementary** Figure 14 and **Supplementary** Figure 15). We also conducted qPCR experiment to detect the rAAV-miniBEND vector DNA distribution across multiple tissue organs (**Supplementary** Figure 16). These results demonstrate the specificity of the rAAV**-**miniBEND system for gene expression in brainECs, with low expression observed in blood vessels from peripheral organs.

To date, no AAVs are available for efficient brainEC labeling when delivered intracranially. In our observations, both AAV-BR1 and AAV-BI30 viruses carrying Ef1α-EGFP recombinant vectors failed to efficiently label brainECs after viral injection directly into the cerebral cortex (**Fig. 2i**). To assess the rAAV-miniBEND system for gene delivery to brainECs through local injection, we used AAV-BR1, AAV-BI30, AAV-PHP.eB-miniBEND, and AAV-9P13-miniBEND viruses. There was a substantial number of neurons and not detectable brainECs labeled with AAV-BR1-Ef1α-EGFP and AAV-BI30-Ef1α-EGFP. Conversely, AAV-PHP.eB and AAV-9P13 viruses, which carried the miniBEND recombinant vector (mPro723-mCis700), demonstrated efficient labeling of brainECs at adolescent mice, and also adult mice, although some neuronal leakage expression still exist (**Fig. 2i** and **Supplementary** Figure 17). These results demonstrate the suitability of the rAAV-miniBEND system for effective gene delivery to brainECs through intracranial injection.

To further validate the efficacy of specific gene deletion in local brain regions, we injected AAV-PHP.eB-miniBEND-Cre viruses into the cerebral cortex of the left hemisphere of *Glut1^fl/fl^* mice, whereas control viruses with EGFP were injected into the same brain region of the right hemisphere as a control (**Fig. 2j**). Immunostaining with antibodies against CD31 and GluT1 was performed to assess the efficiency of *Glut1* gene deletion. AAV-miniBEND-Cre mediated efficient deletion of *Glut1* in brain vascular endothelial cells, with an average efficiency of 61.65 ± 14.89% (n = 2,328 CD31^+^ endothelial cells in total) in the region with local rAAV-miniBEND virus injection (**Fig. 2k–m**).

To assess the transduction efficacy and specificity of the miniBEND system in brainECs from embryonic and neonatal brains, we locally injected AAV-BR1, AAV-BI30, AAV-PHP.eB, or AAV-PHP.eB with miniBEND-Cre into the lateral ventricles of mouse embryos (E15–18) or neonates (P0–4) (**Fig. 2n, Supplementary** Figure 18a). The combination of rAAV-miniBEND with AAV-BI30 resulted in the highest transduction efficiency during the embryonic stage (**Supplementary** Figure 18b-c) or neonatal stages (**Fig. 2q**). During the neonatal stage, AAV-BR1-CAG-EGFP and AAV-BI30-Ef1α-Cre exhibited notable results: the AAV-BR1-CAG virus infected a substantial number of neurons and astrocytes in the cortex, hippocampus, and striatum (**Fig. 2o–q**). Meanwhile, the AAV-BI30-Ef1α group broadly labeled neurons in these regions (**Fig. 2o–q**). The AAV-PHP.eB-CAG group labeled numerous neurons and astrocytes in these brain regions as well (**Fig. 2r–t**). These outcomes indicated that AAV-BR1, AAV-BI30, and AAV-PHP.eB serotypes alone were insufficient for efficient transduction of brainECs of embryonic or neonatal mice. However, both AAV-BI30-miniBEND and AAV-PHP.eB-miniBEND demonstrated highly specific labeling of brainECs (**Fig. 2o,r**), indicating robust cell-type specificity of these recombinant viruses.

To further evaluate the transduction efficacy and specificity of the miniBEND system in rat brainECs, we retro-orbitally injected viruses—AAV-X1.1-Ef1α-Cre, AAV-BI30-Ef1α-EGFP, or AAV-BI30 with miniBEND-Cre—into both wild-type and transgenic reporter rats (*Rosa26-CAG-LSL-EGFP*). Our observations revealed that both AAV-X1.1 and AAV-BI30 exhibited a certain level of transduction efficiency toward rat neurons but displayed low specificity towards rat brainECs in multiple brain regions. However, the combination of rAAV-miniBEND with AAV-BI30 demonstrated both high transduction efficiency and specificity toward rat brainECs (**Supplementary** Figure 19). These results further underscore the successful extended application of the rAAV-miniBEND system in other rodent species, such as rats.

### Optimization of miniBEND promoter for enhancing gene expression

To assess the strength of the miniBEND promoter for gene overexpression, we initially constructed the AAV-mPro1576-EGFP-mCis700 plasmid, packaged it into the AAV-PHP.eB and AAV-PHP.V1 virus, and administered it into the brain via retro-orbital injection to enable brainEC labeling throughout the entire brain. We observed, however, that EGFP fluorescence was not as strong as that we observed in *Ai* reporter strain (**Fig. 3a, b & Fig. 1d**). The inherent activity of the mPro1576 promoter derived from *Tek* limits its broad application, especially in scenarios requiring the high expression of genes of interest.

**Fig. 3.**
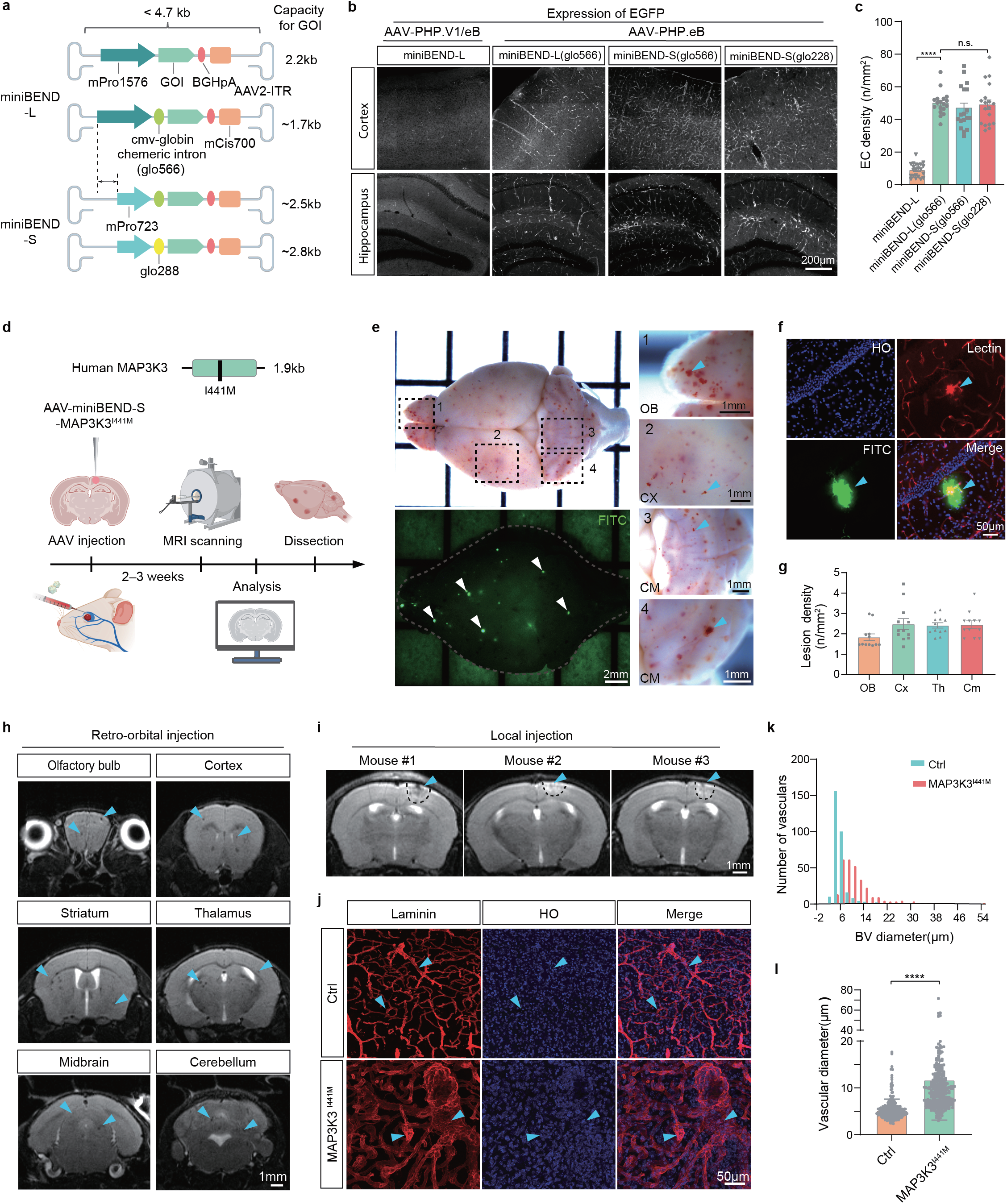
**Development of the rAAV-miniBEND system for modeling CCMs.** **a**, Vector design for the rAAV-miniBEND expression system, including miniBEND promoters (mPro1576 and mPro723), the CMV-globin chimeric intron fused with a globin intron (glo566 or glo228), gene of interest (GOI), Poly A element (bGHpA), miniBEND cis-element (mCis700), internal repeat of recombinant AAV2 vector (AAV2-ITR). **b**, Images of the three expression systems packaged into AAV-PHP.eB or AAV-PHP.V1 serotype virus for miniBEND-L with the original mPro1576 sequence or into AAV-PHP.eB serotype virus for the full-length (L) and shortened (S) mPro723 sequence with the CMV-globin chimeric introns (glo566 or glo228). Expression efficacy was tested in wild-type mice (P60–80) with retro-orbital injection. **c**, The density of brainECs for the mice as described in (b) with 18 regions used for cell counting and statistical analysis per group (n = 18 regions from 3 mice), unpaired two-sided Welch’s t-test, P < 0.0001 (miniBEND-L and miniBEND-L(glo566)), P = 0.769 (miniBEND-L and miniBEND-S(glo228)), \*\*\*\**p* < 0.0001; NS, not significant. **d**, The experimental procedure for CCM modeling with expression of MAP3K3^I441M^. Two delivery strategies were used for AAV administration: retro-orbital injection and local intracranial brain injection. Mouse brains (P60–80) were fixed and dissected after injection of FITC-dextran (2,000 kDa). MRI, magnetic resonance imaging. The diagram was created using BioRender.com. **e**, White-field and fluorescence images of a mouse brain obtained with a stereoscope. Numerous hemorrhagic spots were present on the surface of the olfactory bulb (OB), cerebral cortex (CX), and cerebellum (CM). FITC, green fluorescence observed under blue light excitation. White arrowheads, lesions with FITC leakage. **f**, Representative images from the hippocampus. HO, nuclear staining with Hoechst 33342; lectin, blood vessels stained with Lectin-DyLight 594; FITC, FITC-dextran (2,000 kDa) leaking from blood vessels; blue arrowhead, lesion with leakage. **g**, Density of lesions in different brain regions after retro-orbital administration of AAV-miniBEND- *hMAP3K3*^I441M^, with each group having 12 regions selected for analysis (n = 12 regions from 2 mice). Th, thalamus. **h**, Representative MRI images demonstrating the lesions in the brain after *hMAP3K3*^I441M^ expression mediated by the rAAV-miniBEND system. AAV was administered retro-orbitally. MRI T2 sequence scan shows dark spots within the brain (blue arrowheads), representing bleeding points (i.e., CCM microlesions). **i**, Lesions induced by *hMAP3K3*^I441M^ expression after AAV administration through local brain injection into the cerebral cortex. Focal hemorrhagic phenotype observed under MRI (Blue arrowheads). Representative images from MRI scans of three different mice. **j**, Representative images from brain sections from mice in (i). Ctrl, control region with no lesions. Blue arrowheads, vessels and abnormal vessels in Ctrl and *hMAP3K3*^I441M^ groups, respectively. **k**,**l**, Frequency distribution of blood vessel (BV) diameters (k) and average BV diameter (**l**) in brain regions with or without *hMAP3K3*^I441M^ expression in brainECs delivered by the miniBEND system as in (j). For k, Data distribution is visualized using histograms. For each group, over 150 BVs were selected in (j) for diameter measurements (n = 150 BVs from 5 mice). Data indicate the mean ± s.e.m., unpaired two-sided Welch’s t-test, P <0.0001, \*\*\*\**p* < 0.0001. Image data shown are representative of three (b, e, f), four (h) or five (j) independent experiments.

Hence, further optimization was necessary to enhance the initiation strength of this miniBEND promoter. Recognizing the potential of intron regulatory sequences to enhance promoter initiation strength^42, 44^, we therefore introduced intron regulatory sequences to construct chimeric promoters with enhanced initiation strength. To determine whether these new chimeric promoters maintained endothelial cell specificity, we constructed and packaged AAV-PHP.eB-mPro1576-glo566-intron-EGFP-pA-mCis700 virus, AAV-PHP.eB-mPro723-glo566-intron-EGFP-pA-mCis700 virus, and AAV-PHP.eB-mPro723-glo228-intron-EGFP-pA-mCis700 virus (**Fig. 3a**). The new chimeric promoters exhibited significantly improved initiation strength and endothelial cell–specific expression (**Fig. 3b,c**). Moreover, the truncated version of the chimeric promoters (mPro723-glo566 or glo228) also effectively labeled brainECs and showed little impairment in initiation strength (**Fig. 3b**). Further characterization of the duration of gene expression showed a robust gene expression lasting at least 3 months mediated by rAAV-miniBEND system (**Supplementary** Figure 20). Taken together, the combination of the truncated miniBEND promoter (mPro723) and the smaller intron element (glo228) results in a construct with a total size of 951 bp (less than 1 kb). This allows for the overexpression of larger exogenous genes and mediates robust long-term gene expression.

For gene therapy, the virus is usually administered intravenously. To further evaluate its efficiency, we injected AAV-miniBEND-Cre intravenously with a higher titer (100 μl, 8.7×10^12^ gc/ml) and stained blood vessels in brain sections with anti-Collagen IV. We then calculated the percentage of EGFP^+^/Collagen IV^+^ cells out of the total number of endothelial cells. We observed that the recombination efficiency can reach ∼90–95% in different brain regions (n = 10 regions from 2 mice).(**Supplementary** Figure 21). Further, we intravenously injected AAV-PHPeB-miniBEND-Cre into *Tak1^flox/flox^* transgenic mice. This line has been previously reported to induce regression of blood vessels^45, 46^, Removal of *Tak1* gene from the brain vasculature can increase the regression by 10-fold. After introducing AAV-PHPeB-miniBEND-Cre, we detected blood vessel regression in all major brain regions (including the cerebral cortex, hippocampus, cerebellum, thalamus, hypothalamus, striatum, etc.), and the regression increased dramatically. Additionally, we observed behavioral defects 3–4 weeks after viral infections, similar to the results observed in previous studies. These results indicate that intravenous administration of AAV-miniBEND can efficiently achieve conditional knockout in a whole-brain model(**Supplementary** Figure 22)

### Modeling of CCMs with the rAAV-miniBEND system

The *MAP3K3* c.1323C>G (p.I441M) somatic mutation, initially discovered in patients with peripheral cutaneous vascular malformations^47^, has also been identified in individuals with CCMs^48, 49^. This mutation was subsequently proved to be critical for sporadic CCM formation in mouse models^50, 51^. Leveraging the advantages of the rAAV-miniBEND expression system, we aimed to construct a sporadic CCM disease model in mice. To achieve this, the full-length human version of *MAP3K3* carrying the I441M point mutation (**Fig. 3d**) was cloned into the rAAV-miniBEND vector. The modified vector was then administered either intravenously or locally to enable the expression of the human version of the MAP3K3 kinase protein in vascular endothelial cells throughout the entire brain (intravenously) or in specific brain regions (locally). Expression of human MAP3K3^I441M^ in mouse brain vascular endothelial cells induced widespread brain hemorrhages from malformed vessels (**Fig. 3e–g, Supplementary** Figure 23). This phenotype closely resembled the popcorn-like hemorrhages observed in sporadic CCMs in clinical cases^49, 52^. To further evaluate the blood–brain barrier (BBB) disruption, we injected mice with the high-molecular-weight fluorescent dye fluorescein isothiocyanate (FITC)-dextran (2000 kDa) and then perfused the mice to remove the dye from the circulation system. Diffuse FITC green fluorescence around the hemorrhagic spots in the parenchymal region was observed, indicating the leaking of blood vessels in the lesions (**Fig. 3e–g, Supplementary** Figure 24). This finding was corroborated by endothelial cell staining using Lectin-DyLight 594, a red fluorophore for blood vessel labeling (**Fig. 3f, Supplementary** Figure 24). In additional, we also performed H&E and Prussian blue staining experiments, which detects red blood cells in the perivascular spaces and Hemosiderin deposition in the CCM lesion area (**Supplementary** Figure 25). Our staining results from p62 also showed activation of p62 signaling in the CCMs (**Supplementary** Figure 26). We further analyzed the size and diameter of hundreds of CCM lesions and the density of lesions from brain sections (**Supplementary** Figure 27), we observed enlarged blood vessels with cavernous morphology, and some blood vessels showed leakage of blood cells outside of the vessels throughout brain sections. These results demonstrate the successful establishment of a CCM disease model in mice using the rAAV-miniBEND system.

Magnetic resonance imaging (MRI) has been used to view hemorrhagic lesions in the brains of mice or patients with CCM^49^. In our study, we thus used MRI and observed that expression of MAP3K3^I441M^ in cerebrovascular endothelial cells resulted in a phenotype resembling clinical Grade IV CCM (**Fig. 3h**). When AAV-PHP.eB-miniBEND-MAP3K3^I441M^ viruses were administered intravenously, we obtained a density of CCM lesions throughout the mouse brain similar to what we reported in our previous publications^50^ with AAV-BR1-CAG-MAP3K3^I441M^ (**Supplementary** Figure 28).

We also performed intracranial injections of AAV-PHP.eB-miniBEND-MAP3K3^I441M^ virus locally into the cerebral cortex of mice. After 3 weeks of virus expression, we noticed substantial edema and a hemorrhagic phenotype in the brain regions injected with the virus with MAP3K3^I441M^ (**Fig. 3i**).

Immunostaining results using antibodies against Laminin, which is a marker of blood vessels, indicated vascular distortion in the lesions that resembled cavernous-like dilated structures (**Fig. 3j**). Importantly, the vascular diameters in the CCM lesions were significantly larger than those in the normal control area (**Fig. 3k, l**). These findings provide compelling evidence that the rAAV-miniBEND system can be reliably used for CCM modeling in the mouse brain. We were able to induce focal lesions through local virus injections or global lesions through intravenous injections. Given that AAV-BR1 and AAV-BI30 can label non-brainECs widely in the brain when locally injected **(Fig. 2h, i)**, the miniBEND system emerges as an extremely useful tool for local lesion inductions, especially for time-lapse imaging in mechanistic studies.

### Modeling of bAVM with the rAAV-miniBEND system

To explore the application of the rAAV-miniBEND system for disease modeling of bAVM, which is distinct from CCM^14^, we selected *Braf* as the candidate gene for investigation. A somatic mutation in *Braf* has been identified in two patients with bAVM^53^; however, its role in bAVM development remains unclear. We used rAAV-miniBEND in conditional knock-in mice carrying a *Braf* point mutation (Braf^V600E^) to assess whether this manipulation could mimic the pathological phenotype of human bAVM.

The *Braf-CA*^fl/fl^ mouse model (CA, Cre activated) harbors a point mutation in exon 15 of *Braf* knocked into the mouse genome before the stop codon^54^. Upon Cre recombinase action, the LoxP sites in the *Braf* locus are excised, removing wild-type human exons 15–18 while retaining the mutated form of mouse exon 15 and mouse exons 16–18. Consequently, the downstream fragment carrying the V600E point mutation is transcribed and translated normally, leading to the expression of the Braf^V600E^ mutant protein in the target cells (**Fig. 4a**). Taking advantage of the high efficacy of transduction of brainECs through local injection, we administered AAV-PHP.eB-miniBEND-Cre viruses into the cerebral cortex or hippocampus of mice aged P30–50, inducing the expression of the Braf^V600E^ mutant protein in brainECs in these regions (**Fig. 4b**). We then used MRI to monitor the development of intracranial bAVM. Brain hemorrhages were detected ∼2 weeks after virus injection. By ∼3 weeks the area of cerebral edema had expanded, and at ∼6 weeks the bAVM lesion area exhibited tissue necrosis and compression of the hippocampus (**Fig. 4c, d**). In addition, malformed vasculature was observed with a laser speckle blood flow imaging system transcranially (**Fig. 4e**). With H&E staining, we also observed obvious vascular dilation with feeding/draining vessels having much smaller diameters, typical of AVMs (**Supplementary** Figure 29). We performed immunostaining of these abnormal vessels and they were xx positive. We observed that the microvessels dilated/expanded, and that the feeding vessels (arteries) and draining vessels (veins) had small diameters. Notably, the focal bAVM model mice (i.e., *Braf-CA^fl/fl^* mice with injection of AAV-PHP.eB-miniBEND-Cre virus) began to die on post-injection day 20 (PID20, n = 12 model mice), resulting in a final survival rate of <50% at 3^th^ month after virus injection (**Fig. 4f**). This focal bAVM model in mice resulted in a phenotype similar to human intracranial AVMs^14, 55^.

**Fig. 4.**
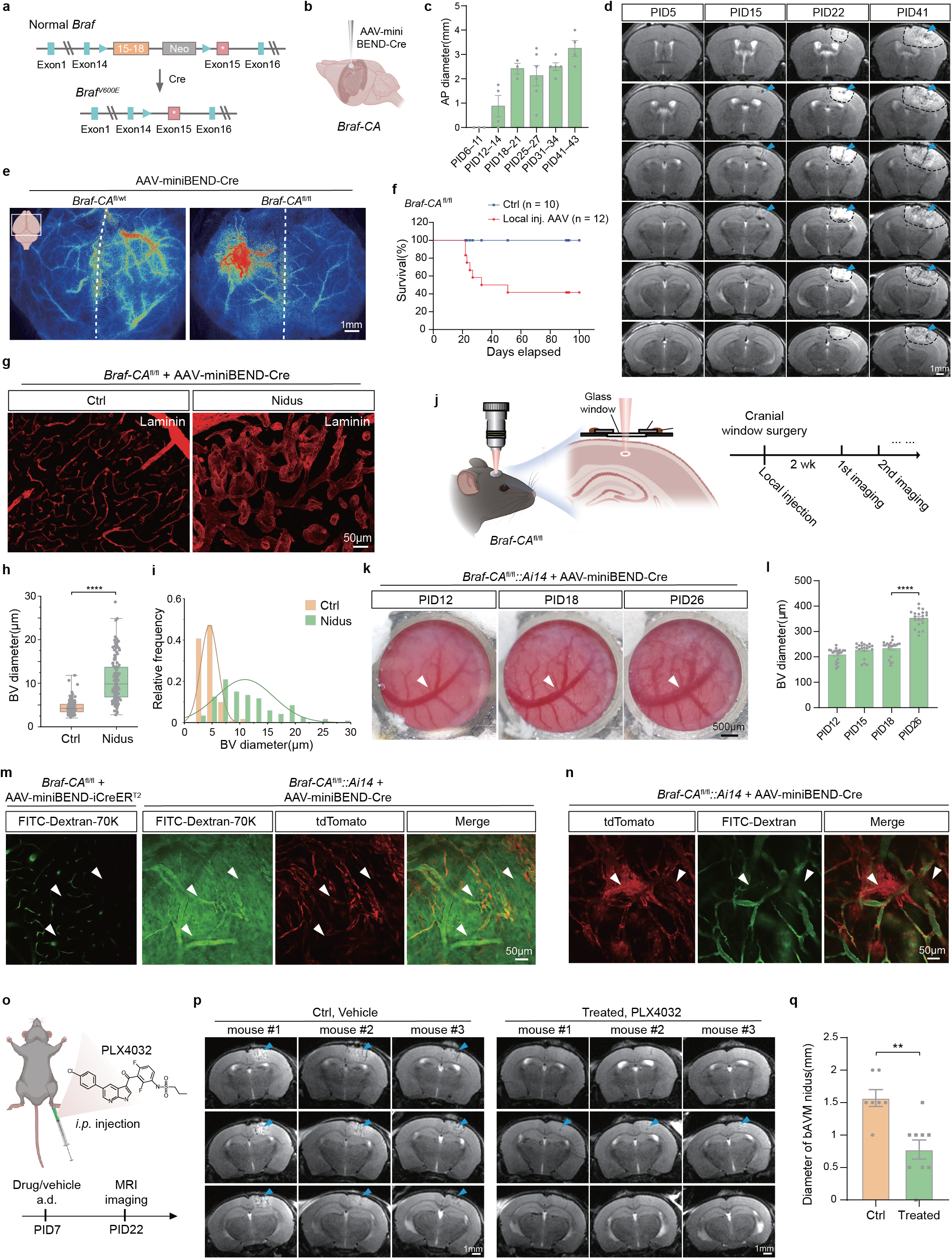
**Modeling bAVM with the rAAV-miniBEND system.** **a,b**, Working principle (a) and experimental schematic (b) for developing a bAVM model with *Braf-CA^fl/fl^* transgenic mice through the rAAV-miniBEND system. The endogenous mouse exons are in blue and wild-type human exons are in orange. Exons 15-18 are the wild-type human exons 15–18, which are transcribed in the absence of Cre recombinase. The red box with a white dot represents mouse exon 15 with the point mutation that results in V600E. **c**, Lesion size detected by MRI at different time points (PID, post-injection days) after local injection of AAV-PHP.eB-miniBEND (mPro723-mCis700)-Cre viruses in *Braf-CA^fl/fl^* mice. AP diameter is the diameter of the AVM nidus in the anterior–posterior direction in mice measured via MRI imaging at different time points: PID6–11 (n = 3 mice), PID12–14 (n = 4 mice), PID18–21 (n = 3 mice), PID25–27 (n = 6 mice), PID31–34 (n = 5 mice), PID41–43 (n = 4 mice); data indicate the mean ± s.e.m. **d**, Representative longitudinal T2 MRI images of mouse brains as in (**c**). Blue arrowhead, bAVM lesion. **e**, Laser speckle blood flow imaging showing the blood flow in the *Braf-*bAVM mouse model PID38 after injection of AAV-PHP.eB-miniBEND-Cre viruses at the right hemisphere. Pseudo-color indicates the blood flow volume index, with warmer colors indicating increased blood flow. The dashed line indicates the midline of the brain, n = 4 mice for each group. **f**, Survival curve of mice with focal bAVM lesions. Experimental group consisted of *Braf-CA*^fl/fl^ mice with local injection of AAV-PHP.eB-miniBEND-Cre (Local inj. AAV, n = 12 mice); the control group consisted of *Braf-CA*^fl/fl^ mice with local injection of AAV-PHP.eB-miniBEND-S(glo566)-EGFP (Ctrl, n = 10 mice). **g**, Representative images of blood vessels stained by antibodies against Laminin in the control region (Ctrl) and AVM lesion area (Nidus). **h,i**, Average blood vessel (BV) diameter (h) and frequency distribution of blood vessel diameters in the control (Ctrl, n = 123 vessels) and AVM model (Nidus, n = 121 vessels) mice. For Ctrl group of h, minima = 2.163, maxima = 11.86, median = 4.257, 25% percentile = 3.455, 75% percentile = 5.197; for Nidus group of h, minima = 2.777, maxima = 28.72, median = 9.868, 25% percentile =6.838, 75% percentile = 13.752, Box plots display the median (middle line), with the box’s lower and upper bounds indicating the 25th and 75th percentiles, respectively. Whiskers extend to the minima and maxima, unpaired two-sided Welch’s t-test, P <0.0001, ****p < 0.0001. For i, data distribution is visualized using histogram, overlaid with the corresponding normal distribution fit. **j**, Time-lapse experiment for imaging AVM development with two-photon laser excitation microscopy. Mice were imaged once every 6–8 days beginning at 2 weeks after local injection. **k**, Representative images captured by a stereoscope showing the gradual enlargement of blood vessels during AVM development. White arrowhead, the enlarged blood vessel. **l**, Data summarizing alterations in the diameter of large blood vessels (BVs) in the superficial cortical region at different time points (PID12, 15, 18, and 26), 20 regions along the main vessel were measured in each group (n = 20), one-way ANOVA with Kruskal-Wallis post-hoc test, P < 0.0001; unpaired Mann-Whitney test between PID18 and PID26, P < 0.0001, \*\*\*\**p* < 0.0001. Data indicate the mean ± s.e.m.. **m**, Using time-lapse two-photon *in vivo* imaging with FITC-dextran (70 kDa) (green) to detect the integrity of the BBB. White arrowheads, leakage of the green fluorescent dye into the brain parenchyma. Red, tdTomato signal. **n**, Images of the EndMT in the indicated mice observed at PID18 through live imaging. White arrowheads, endothelial cells with mesenchymal-like morphology. **o**, Molecular structure of the BRAF^V600E^-selective kinase inhibitor PLX4032 and experimental procedure. *Braf-CA*^fl/fl^ mice were first locally injected with AAV-PHP.eB-mPro723-Cre-mCis700 virus for bAVM disease modeling. PLX4032 dosage: 75 mg/kg body weight per injection every two days for 7 injections. DMSO (vehicle) injections were used as the control. Drug and DMSO were administered via intraperitoneal injection. **p**, Representative longitudinal T2 MRI images of Braf-bAVM mice with or without PLX4032 treatment as in (o). Blue arrowhead, bAVM lesion. Each row represents one mouse. **q**, The diameter of AVM lesions in Braf-bAVM mice with or without PLX4032 treatment as in (o). Treated group (n = 9 mice), control group (n = 7 mice). unpaired two-sided Welch’s t-test, two-tailed P value = 0.0012, \*\**p* < 0.01. Data indicate the mean ± s.e.m., along with individual data points in (c, h, l, q); Image data shown are representative of four (e) or two (g, k, m-n) independent experiments. Diagrams in (a), (b), (j), (o) were created using BioRender.com.

Additionally, mice displayed symptoms such as hemiparesis and occasional seizures (**Supplementary Video 1**), aligning with the clinical manifestations observed in some bAVM patients^55^. Further, we administered AAV-miniBEND-Cre to *Braf-CA^fl/fl^* mice via retro-orbital injection and observed many micro-lesions with vascular malformations in the brain. With H&E staining, we observed the dilation of vessels, leakage of vessels, and a significant increase in the diameter of malformed vessels in the lesion areas (**Supplementary** Figure 30). Additionally, we observed some leakage of dilated blood vessels, but unlike those induced with AAV-miniBEND-MAP3K3^I441M^, the vessels showed a dilated morphology rather than a cavernous shape.

We then carried out TUNEL staining to investigate the cellular phenotypes in bAVM lesions modeled by rAAV-miniBEND with Braf activation. There were significant higher TUNEL^+^ signals in the bAVM lesions as compared to control regions in the contralateral hemisphere **(Supplementary** Figure 31a**, b**). The strongest TUNEL signals were observed near the area of vascular damage, and most of those regions were not co-stained by NeuN **(Supplementary** Figure 31c**, d**), indicating that brain hemorrhage might induce neuronal apoptosis within the lesion region, potentially leading to impaired local brain function and likely contributing to the mortality observed in these mice. We further stained blood vessels with antibodies against Laminin. Our results demonstrated predominantly enlarged and deformed blood vessels in the bAVM lesion, with almost no capillary-sized microvessels present (**Fig. 4g** and **Supplementary Video 2**). The blood vessel diameter in the bAVM lesion area was significantly larger than that in the contralateral control brain region (**Fig. 4h, i**).

To directly examine cellular alterations in brain vasculature during *Braf*^V600E^-mediated bAVM formation, a cranial window was created in the mouse skull at the same time as the local injection, enabling real-time imaging of blood vessels beneath the dura mater, which began at 2 weeks after injection **(Fig. 4j)**. The blood vessels exhibited a progressive increase in diameter within 2–3 weeks after injection of AAV-miniBEND-Cre viruses (**Fig. 4k, l**). Additionally, alterations in functional blood flow within the dilated vessels were examined using laser speckle flowmetry for daily time-lapse imaging. A significant increase in blood flow in the corresponding vessels near the viral injection site was observed (**Supplementary** Figure 32). These results indicated that the somatic variant *Braf*^V600E^ is able to induce vascular dilation, leading to an overall increase in blood flow and affecting the hemodynamics of the local brain region.

Through *in vivo* two-photon imaging experiments, a significant plasma leakage was observed in the circulation of Braf-bAVM mice, indicating impairment of the BBB integrity (**Fig. 4m**). Arteriovenous shunts, enlarged vessel segments and abnormal blood flow were also observed in the endothelial Braf^V600E^ activated vasculatures (**Supplementary Video 3, Supplementary Video 4, and Supplementary** Figure 33).

We perfused the mouse brain vasculature with Latex, which is known to be restricted to arteries/arterioles and blocked before reaching capillaries. We detected AV shunt-like malformations in the brain with local injection of AAV-PHP.eB-miniBEND-Cre or with intravenous injection of a low dose of AAV-PHP.eB-miniBEND-Cre in *Braf-CA^fl/fl^* mice (**Supplementary** Figure 34).

We further performed whole-brain clearing with PEGASOS (a polyethylene glycol (PEG)-associated solvent system) and imaged the entire brain vasculature in *Braf-CA*^fl/fl^*::Ai14* mice using intravenous injection of AAV-PHP.eB-miniBEND-Cre. This approach allowed us to visualize malformations in the blood vessels with a 3D view, showing expansion of vessels in endothelial cells (**Supplementary** Figures 35 and 36).

Furthermore, clusters of mesenchymal-like cells undergoing the endothelial-to-mesenchymal transition (EndMT)^56^ were identified (**Fig. 4n**). These cells were capable of forming blood vessels, as evidenced by the presence of blood flow within the vessel lumen (visualized by FITC-dextran). This phenomenon suggested that EndMT is potentially one of the cellular processes involved in the initiation of bAVM. In addition, we also observed the activation of astrocytes and microglia in lesions induced by Braf^V600E^ after local administration of AAV-PHP.eB-miniBEND-Cre (**Supplementary** Figure 37**, Supplementary** Figure 38). The coverage of GFAP was significantly increased in the nidus region. The processes of microglia were shortened and conjugated near the malformed blood vessels.

To investigate the therapeutic potential of this model, PLX4032, a small-molecule blocker known to inhibit the kinase activity of BRAF^V600E^ in mice^57^, was administered to mice 7 days after AAV-miniBEND-Cre (AAV-PHP.eB-mPro723-Cre-mCis700 virus) viral injection. An equal volume of the vehicle dimethyl sulfoxide (DMSO) was given to control mice. Time-lapse imaging with MRI commenced on PID22 (**Fig. 4o**). The diameter of bAVM lesions, the density of malformed vessels, and the diameter of malformed vessels was significantly reduced in the drug-treated mice as compared to the control group (**Fig. 4p,q, Supplementary** Figure 39), indicating that activating BRAF kinase and its downstream signaling are sufficient for bAVM pathogenesis and progression. Additionally, these results suggested that Vemurafenib, a drug derived from PLX4032 that is used to treat malignant melanoma, could potentially be repurposed as a treatment for bAVM patients. The Braf-bAVM mouse model developed in this study could serve as a valuable platform for therapeutic drug screening in the context of bAVM.

### Mechanistic studies of bAVM initiation and progression with the rAAV-miniBEND system

To elucidate the molecular mechanisms underlying the initiation and development of bAVM, single-nucleus RNA sequencing (snRNAseq) of tissues from bAVM lesions was conducted in *Braf-CA*^fl/fl^ mice at two time points (PID15 and 27) after local injection of AAV-miniBEND-Cre viruses. Given that AAV starts expression about 1 week after injection, and the mice start dying after 3 weeks, we consider PID15 and 27 as early and late developmental stages of AVMs. The control samples were collected from the contralateral hemisphere of the same brain (**Fig. 5a**). Through clustering analysis and basic cell type annotation, we identified multiple cell clusters in the single-nucleus transcriptomes of bAVM lesion areas at both early stage (PID 15) and late stage (PID 27) (**Fig. 5b,c**).

**Fig. 5.**
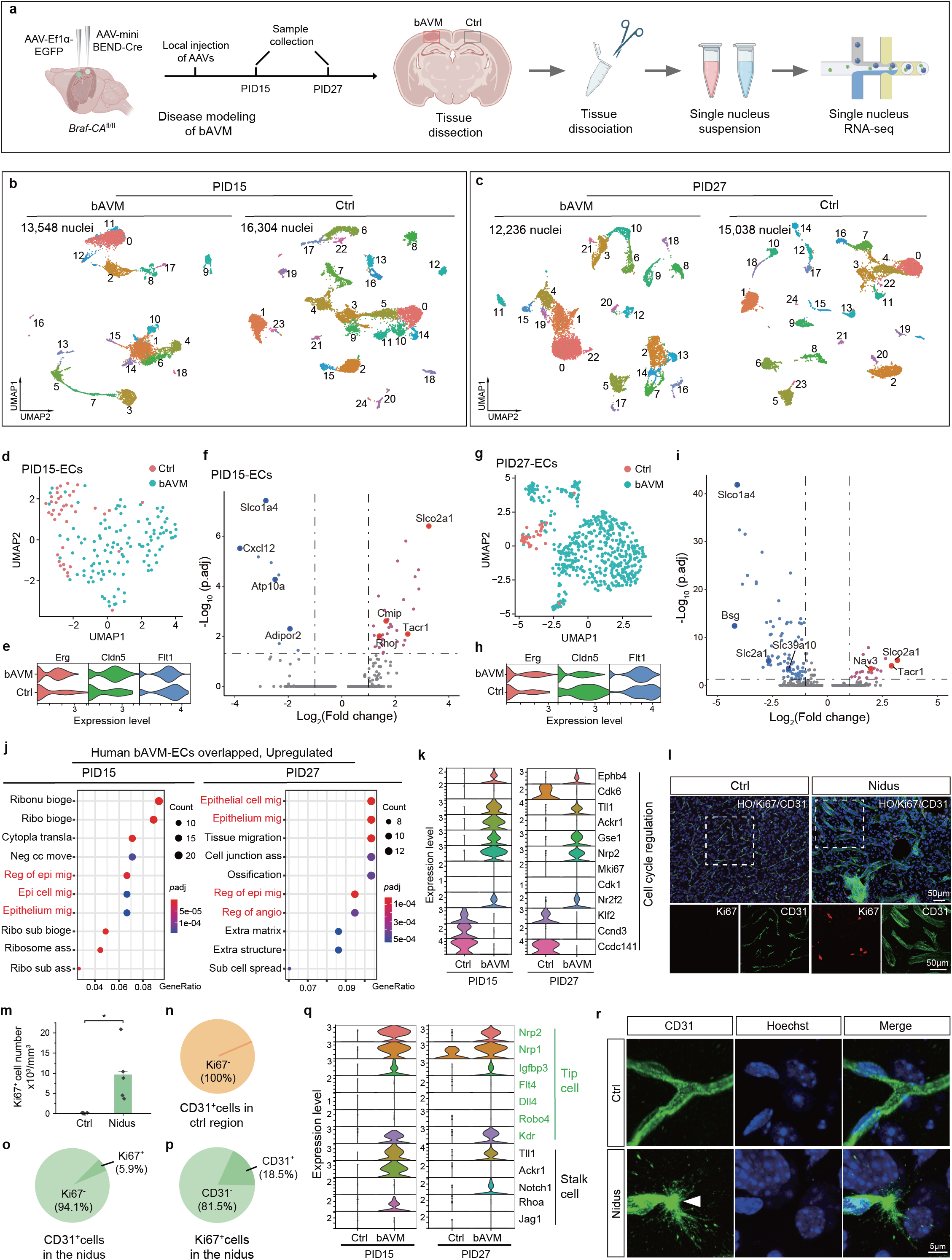
**rAAV-miniBEND and snRNAseq reveal the molecular landscape of bAVM initiation and progression.** **a**, bAVM disease modeling and workflow of single nucleus transcriptome (snRNAseq) analysis. an injection shown for AAV-EF1α-EGFP represents the control contralateral hemisphere, AAV-miniBEND-Cre represents the modeling of bAVM in one hemisphere. Diagram was created using BioRender.com. **b,c**, UMAP plot after dimension reduction clustering analysis of single-nucleus transcriptome data from both nidus (bAVM) and control (Ctrl) regions at PID15 (b) and PID27 (c). All clusters are labeled with numeric identifiers in the graph, which correspond to cells listed here. PID15-bAVM region: Neurons (clusters 1, 4, 6, 10, 14, 18); ECs (15); Microglia (0, 2, 11,12); Astrocyte (13); Oligodendrocyte (3, 7); Immune cells (8, 9, 17); NG2 glia (8); PID15-Ctrl region: Neurons (0, 2, 3, 4, 5, 7, 9, 10, 11, 12, 13, 14, 15, 16, 22); ECs (24); Microglia (6, 17); Astrocyte (21); Oligodendrocyte (1, 18); Immune cells (19); NG2 glia (8); PID27-bAVM region: Neurons (2, 5, 7,12, 13,14,16,17,20); ECs (8); Microglia (0,1,22); Astrocyte (9); Oligodendrocyte (6,10); Immune cells (4,11,15,19); NG2 glia (3, 21); PID27-Ctrl region: Neurons (0,3,4,7,11,12,13,14,16,17,22); ECs (24); Microglia (2,8,20); Astrocyte (9); Oligodendrocyte (1,18,19); Immune cells (6,21,23); NG2 glia (10); **d,g**, UMAP two-dimensional plot after dimension reduction, representing the clustering of combined single-nucleus transcriptomic data from all endothelial cells in the bAVM and control regions at PID15 (d) and PID27 (g). **e,h**, Violin plot representing the expression level of endothelial cell gene markers (*Erg*, *Cldn5*, *Flt1*) in both clusters of ECs (bAVM and Ctrl) at PID15 (e) and PID27 (h). **f,i**, Volcano plot representing the differentially expressed genes in endothelial cells from the bAVM lesion and control brain region at PID15 (f) and PID27 (i). Significance cutoff: |log_2_(fold change) ≥ 1 and p*adj* ≤ 0.05. Upregulated genes highlighted: *Slco2a1*, *Cmip*, *Rhoj*, and *Tacr1* at PID15 and *Slco2a1*, *Nav3*, and *Tacr1* at PID27; downregulated genes highlighted: *Slco1a4*, *Cxcl12*, *Atp10a*, and *Adipor2* at PID15 and *Slco1a4*, *Bsg*, *Slc2a1*, and *Slc39a10* at PID27. Two-sided Wald test was used, and P values were adjusted for multiple testing using the Benjamini-Hochberg method (adjusted P < 0.05). **j**, GO enrichment analysis of upregulated genes of endothelial cells (ECs) in bAVM lesions at PID15 and PID27 that overlapped with human bAVM-EC DEGs (human bAVM DEGs of EC clusters were from ref. 55). 249 genes overlapped at PID15 and 120 genes overlapped at PID27, cutoff: log_2_(fold change) > 0.5. Ribonu bioge, Ribonucleoprotein complex biogenesis; ribo bioge, ribonucleoprotein complex biogenesis; cytopla transla, cytoplasmic translation; neg cc move, negative regulation of cellular component movement; reg of epi mig, regulation of epithelial cell migration; epi cell mig, epithelial cell migration; epithelium mig, epithelium migration; ribo sub bioge, ribosomal small subunit biogenesis; ribosome ass, ribosome assembly; ribo sub ass, ribosomal small subunit assembly; epithelial cell mig, epithelial cell migration; epithelium mig, epithelium migration; cell junction ass, cell junction assembly; reg of epi mig, regulation of epithelial cell migration; reg of angio, regulation of angiogenesis; extra matrix, extracellular matrix organization; extra structure, extracellular matrix organization; sub cell spread, substrate adhesion-dependent cell spreading. **k**, Expression profile analysis of DEGs encoding cell cycle–related regulatory proteins in endothelial cell clusters from the bAVM lesion area and control brain region during both early AVM development (PID15, left) and late AVM development (PID27, right). **l**, Representative images from immunostaining with antibodies against CD31 and Ki67 in the control brain area (Ctrl) and the bAVM lesion area (Nidus). HO, nuclear staining with Hoechst 33342. **m**, The density of Ki67^+^ cells per square millimeter in the control (Ctrl) and nidus regions (Nidus). Data indicate the mean ± s.e.m. along with individual data points, unpaired two-sided Welch’s t-test, P = 0.0361, \**p* < 0.05. **n,o**, The dividing endothelial cells (CD31^+^Ki67^+^ cells/CD31^+^ cells) in the control region (n) and bAVM region (o). **p**, The proportion of dividing endothelial cells among all dividing cells (CD31^+^Ki67^+^ cells/Ki67^+^ cells) in the nidus region. **q**, Marker genes related to tip cells and stalk cells in endothelial cell clusters among DEGs from both early (PID15) and late (PID27) developmental stages of AVMs. **r**, Representative image of an individual tip cell in the bAVM lesion area. White arrowhead indicates a filopodia extending from the tip cell. Tip cells were labeled with antibodies against CD31, and nuclei were stained with Hoechst 33342. Image data shown are representative of three (l, r) independent experiments.

To identify differentially expressed genes (DEGs) in endothelial cells between the control and bAVM groups, the expression matrix of endothelial cell subtypes was analyzed (**Fig. 5d,g**). Multiple genes, including *Slco2a1, Tacr1, Cmip* and *Rhoj* were significantly upregulated in the bAVM group, whereas other genes such as *Slco1a4*, *Cxcl12, Atp10a, Bsg, Slc1a4* and *Slc39a10* were dramatically downregulated (**Fig. 5f,i**). Notably, downregulated genes *Slco1a4*, *Spock2*, and *Slc39a10* were consistent with results from single-nucleus transcriptome sequencing of brainECs from affected tissues of bAVM patients (*SLCO1A2*, *SPOCK3*, and *SLC39A10*, respectively); similarly, upregulated genes *Tll1* and *Ackr1* were consistent with upregulated genes *TLL1* and *ACKR1* in these patients^58^. These results suggested that the Braf-bAVM model replicated the molecular alterations observed in bAVM lesions in patients.

Expression profiling analysis of genes that encode cell cycle–related regulatory proteins in all endothelial cell clusters revealed higher expression of proliferation-related genes such as *Mki67*, *Cdk1*, *Cdk6*, *Ccnd3*, *Gse1*, and *Nrp2* in endothelial cell clusters from bAVM lesions, whereas these genes were expressed at lower levels in endothelial cells from the control group (**Fig. 5k**). Both tip cells and stalk cells play crucial roles in angiogenesis. Tip cells lead blood vessel sprouts, followed by more proliferative stalk cells. To confirm endothelial cell proliferation in bAVM lesions, brain sections were stained with an antibody against Ki67, a marker for proliferating cells. There were significantly more proliferating cells in the nidus than in the control contralateral brain tissue, although only 18.5% of Ki67^+^ cells were CD31^+^ in lesions (**Fig. 5l–p**). Thus, a substantial portion of dividing cells were not endothelial cells. We also observed the upregulation of tip cell marker genes (*Flt4*, *Dll4*, *Robo4*, *Kdr*, *Nrp2*, *Nrp1*) and stalk cell marker genes (*Notch1*, *Jag1*, *Rhoa*) in brain vascular endothelial cells from bAVM lesions (**Fig. 5q**). This was confirmed by immunostaining brain sections with anti-CD31, which revealed numerous tip cells in bAVM lesions as compared to the absence of tip cells in the control brain region (**Fig. 5r**).

In our staining results, we observed the EndMT phenomenon in AVM lesions (**Fig. 4n**). Our gene set enrichment analysis (GSEA) showed that EndMT-related genes were upregulated in endothelial cell clusters within the Braf-bAVM lesion area (**Supplementary** Figure 40), which is similar to the result observed in human bAVM single cell transcriptome data^58^. An immunostaining experiment further confirmed that αSMA, a marker protein for mesenchymal cells^56, 59^, is upregulated in bAVM lesion areas and is co-localized with CD31 expression (**Supplementary** Figure 40), indicating that endothelial cells are undergoing the EndMT process during bAVM initiation.

To obtain more endothelial cells for further analysis and elucidate the molecular mechanisms of Braf^V600E^-mediated bAVM, we performed single-cell RNA sequencing of endothelial cells collected from the cerebral cortex in *Braf-CA^fl/fl^* mice after they were injected with AAV-miniBEND-Cre (AAV-PHP.eB-mPro723-Cre-pA-mCis700, **Fig. 6a**). The control group without bAVM was collected at the same age and injected with control virus (AAV-PHP.eB-mPro723-EGFP-pA-mCis700). After extracting recognized endothelial cells from a total of 28,954 brain cells, we obtained 5,646 endothelial cells in total, with 2,624 cells from the bAVM group and 3,022 cells from the control group. Clustering analysis and cell subpopulation annotation revealed the presence of arterial, venous, and capillary endothelial cells (aECs, vECs, and cECs, respectively) (**Fig. 6b–d**), based on well-accepted markers^60^ (*Pecam1* for all ECs, *Slc7a5* for cECs, *Ackr1* for vECs, and *Bmx* for aECs). From cell proportion and MELD analysis between the bAVM and control groups, we observed significant alterations in the cell proportion of vECs. vECs likely represent the most relevant EC subpopulation responsible for bAVM formation (**Fig. 6e, f**).

**Fig. 6.**
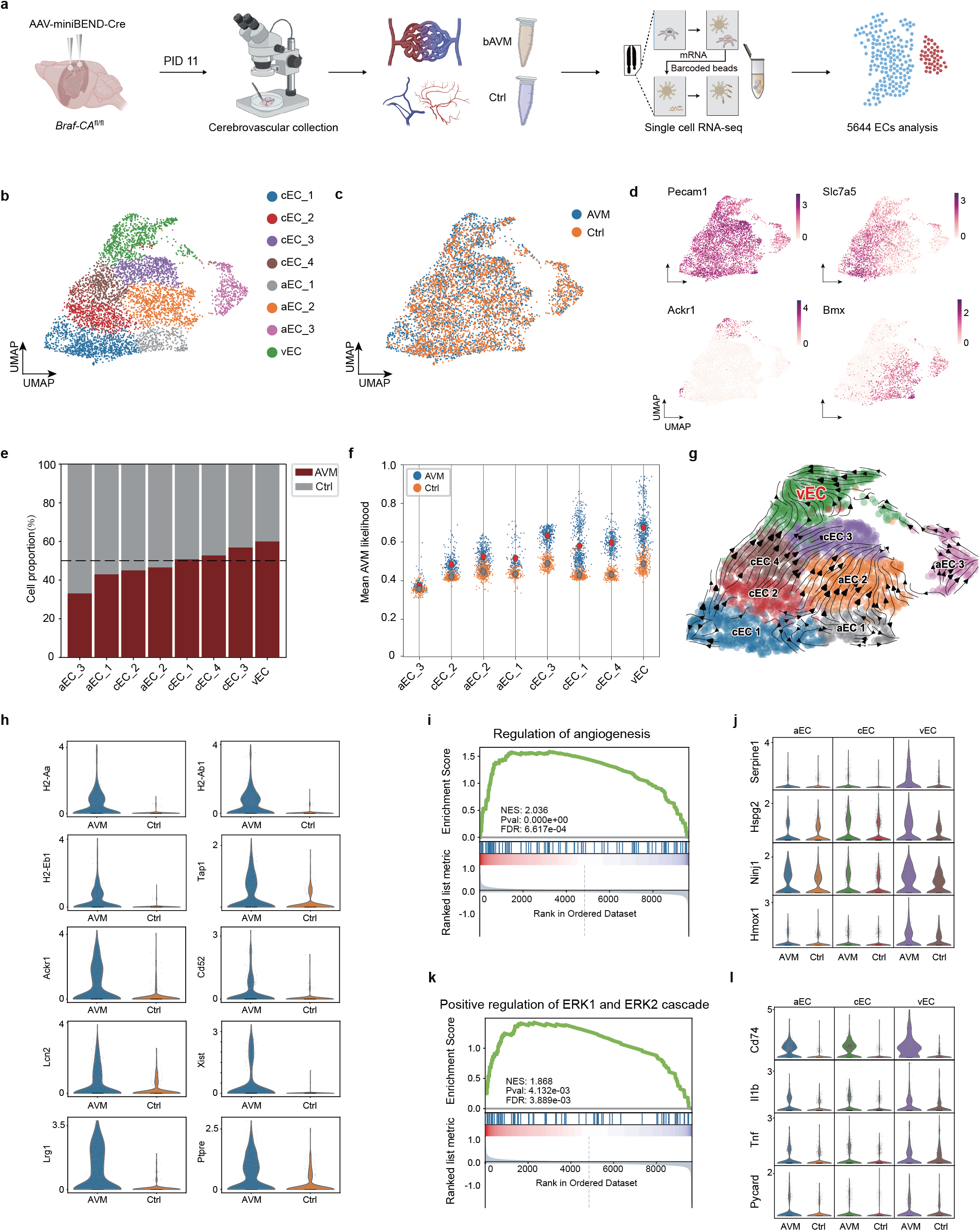
**scRNAseq analysis of brainECs from Braf^V600E^-mediated bAVM** **a**, Schematic illustration of bAVM disease modeling, endothelial cell collection, and the workflow of single-cell transcriptome (scRNA-seq) analysis of endothelial cells from control and bAVM groups. Mice from control group were injected with AAV-PHP.eB-miniBEND-L(glo566)-EGFP virus, Diagram was created using BioRender.com. **b**, UMAP two-dimensional plot after dimension reduction, representing the subclusters from all endothelial cells in the bAVM and control mouse cortex at PID11. (aEC, arterial endothelial cell; vEC, venous endothelial cell; cEC, capillary endothelial cell.) **c**, The UMAP two-dimensional plot of dimensionality reduction clustering analysis of single-cell transcriptome data from both the nidus (bAVM) and control brain (Ctrl) at PID11 shows distinct clustering patterns for endothelial cells in each group. In the bAVM group, a total of 2,624 endothelial cells were identified, while the Ctrl group comprised a total of 3,022 endothelial cells. **d**, The UMAP two-dimensional plot after dimension reduction represents the expression levels of endothelial cell gene markers, including Pecam1 for whole endothelial cells, Slc7a5 for capillary endothelial cells, Ackr1 for venous endothelial cells, and Bmx for arterial endothelial cells, in both clusters of endothelial cells (ECs) from the bAVM and Ctrl groups. **e**, The bar plot represents the proportion of endothelial cell subtypes isolated from both the bAVM and Ctrl groups. **f**, The jitter plot illustrates the likelihood of bAVM association with each endothelial cell type. Blue values indicate the average likelihood of the cell being isolated from bAVM, while orange values represent the average likelihood of the cell being isolated from Ctrl. **g**, The UMAP visualization depicts the RNA velocity of endothelial cell states in bAVM, with colors representing different subjects. **h**, The violin plot illustrates the expression level of up-regulated genes in vascular endothelial cells (vECs), including *H2-Aa, H2-Ab1, H2-Eb1, Tap1, Ackr1, Cd52, Lcn2, Xist, Lrg1*, and *Ptpre*, associated with inflammation, endothelial proliferation, and dysfunction in both clusters (bAVM and Ctrl) of endothelial cells. **i**, The Gene Set Enrichment Analysis (GSEA) demonstrates the enrichment of the “regulation of angiogenesis” gene set in vascular endothelial cells (vECs) from bAVM samples. **j**, The violin plot illustrates the expression levels of genes enriched in the regulation of angiogenesis pathway (*Serpine1, Hspg2, Ninj1, Hmox1*) in arterial endothelial cells (aECs), capillary endothelial cells (cECs), and venous endothelial cells (vECs) from both bAVM and Ctrl clusters. **k**, The Gene Set Enrichment Analysis (GSEA) demonstrates the enrichment of the positive regulation of the ERK1 and ERK2 cascade in venous endothelial cells (vECs) from bAVM samples. **l**, The violin plot illustrates the expression levels of genes enriched in the positive regulation of the ERK1 and ERK2 cascade pathway (*Cd74, Il1b, Tnf, Pycard*) across both clusters (bAVM and Ctrl) of arterial endothelial cells (aECs), capillary endothelial cells (cECs), and venous endothelial cells (vECs). For i and k, Gene Set Enrichment Analysis was performed using permutation testing, two-sided P values were calculated, and FDR correction was applied.

To further elucidate the alterations in vECs, we computed the differentially regulated genes across conditions (bAVM vs. control). Based on expression level alterations and gene function, we identified a list of genes involved in vascular inflammation, endothelial proliferation, and dysfunction (**Fig. 6h**). This suggests that vECs in bAVM lesions were active in vascular inflammation and might promote angiogenesis. GSEA analysis showed that related genes from two pathways, Regulation of Angiogenesis and Positive Regulation of ERK1 and ERK2 cascade, were mainly upregulated in vEC clusters in the bAVM group (**Fig. 6i** and **k**). By comparing the expression levels of different genes in aECs, cECs, and vECs in these two pathways, we observed that vECs exhibited more substantial alterations, and the genes showed higher expression levels in response to vascular malformation, suggesting a more critical role of vECs in the development of bAVM (**Fig. 6j** and **i**). In the future, more sophisticated investigations are required to elucidate the underlying molecular mechanisms.

## Discussion

The development of the rAAV-miniBEND system, as presented in this study, addresses several crucial challenges in the field of gene therapy and disease modeling, particularly in the context of brain vascular diseases. The optimized miniature promoters and cis-regulatory elements have significantly reduced the space required for exogenous gene insertion, enabling the delivery of large protein-encoding genes (2.2– 2.5 kb) within the limited payload capacity of AAV viral genomes. This advancement is particularly valuable in the context of *in vivo* genome editing and treatment of neurological disorders, for which the size of therapeutic genes can be substantial^61^. Moreover, the specificity and efficiency of the rAAV-miniBEND system in delivering genes to brainECs at various developmental stages and its ability to achieve efficient local gene delivery represent notable advancements. Previous AAV systems, such as AAV-BR1^6^ and AAV-PHP.eB^7^, faced challenges in achieving such precise and effective targeting. The enhanced biosafety profile, with minimal ectopic expression in peripheral vasculature and limited immune response, adds another layer of reliability to this system. This is crucial for ensuring the safety and efficacy of gene therapy approaches, especially those involving the central nervous system^62^.

The optimization of the miniBEND promoter, although currently of moderate strength, represents an important step toward overcoming the limitations associated with promoter strength. While there are caveats with this system, especially in cases requiring exceptionally high gene expression, the progress described here provides a foundation for future improvements in promoter design for gene therapy applications.

The rAAV-miniBEND system exhibits high labeling efficiency in developing rodent brains (including mouse and rat), demonstrating higher specificity than other AAV systems in developing mouse brains (including those of neonatal mice). It allows highly efficient gene delivery within the brain vasculature when administered intracranially/locally, a capability unmatched by existing AAV systems. Although a promoter can be used in rAAV vectors, the limited space for the exogenous gene and promoter (i.e., <4.7 kb) still hampers disease modeling and gene therapy applications. Our truncated elements save 2.5 kb of space for the gene(s) of interest.

The establishment of animal models for cerebrovascular diseases, such as bAVMs and CCMs, is of importance for understanding the underlying mechanisms and for developing potential therapies. The successful creation of a stable mouse model for focal AVMs in the brain, coupled with detailed characterization using various imaging and molecular techniques, demonstrates the applicability and reliability of the rAAV-miniBEND system in disease modeling. The real-time observation of cellular morphology and hemodynamics within the focal lesion area using advanced imaging techniques provides valuable insights into disease progression and pathology. The potential applications of the rAAV-miniBEND system extend beyond rodents, with the prospect of establishing preclinical large-animal disease models in species such as pigs and non-human primates, such as marmosets or macaques^63–65^.

These models are important for bridging the gap between basic research and clinical trials, providing a more comprehensive understanding of disease mechanisms and allowing for thorough preclinical evaluations of candidate therapeutic interventions^66^.

Researcher previously utilized AAV-BR1 to overexpress Kras^G12V^ in endothelial cells through retro-orbital venous sinus injection in 5–6-week-old male wild-type mice, resulting in a brain AVM phenotype for further study^67^. Fish and colleagues employed a brain-endothelial cell-specific Cre/CreER line (*Slco1c1-CreER*) to induce brain AVMs in a lox-stop-lox Kras conditional mouse strain (referred to as *Kras^G12D^*)^22^. Initiation of mutant Kras^G12D^ expression at P1 or in adult mice (2–4 months old) required up to 8 weeks to induce bAVMs in *Slco1c1-CreER::Kras^G12D^* animals, with brain AVMs occurring in only about half of all animals examined (*Slco1c1-CreER::Kras^G12D^*), posing challenges for further treatment or live imaging studies. Our strategy to use AAV-miniBEND to induce brain AVMs is similar to previous studies but also has differences. Notably, the AAV-miniBEND-Cre exhibits specificity and high efficacy for labeling blood vessels through local injections, providing a unique advantage compared to other systems (AAV-BR1 infection causes a lot of non-specific labeling, as shown in our results). We successfully induced local brain AVMs by locally injecting AAV-miniBEND-Cre into *LSL-Braf*^V600E^ mice. Moreover, based on our experience, we can induce brain AVMs through retro-orbital injection of AAV-miniBEND, with a very high success rate and a timeframe of only about 2 weeks to observe a typical AVM phenotype.

The rAAV-miniBEND system presented in this study represents a marked advancement in the field of gene therapy and disease modeling. Its ability to achieve specific and efficient gene delivery, coupled with enhanced biosafety features and applicability to various disease models, positions it as a powerful tool for researchers and clinicians working on neurological disorders and vascular diseases. The ongoing optimization and expansion of this system hold great promise for the future of gene therapy and translational research.

## Supplementary Figure Legends

**Supplementary Figure 1**

***Tek* gene expression is restricted to endothelial cells of both mouse and human brains.**

**a,** Expression profile of *Tek* in the single-cell transcriptome of the mouse brain vasculature system (data sourced from a public database^68^). PC, Pericytes; vSMC, venous SMC; aaSMC, arteriolar SMC; aSMC, arterial SMC; MG, Microglia; FB1, fibroblast-like type 1; FB2, fibroblast-like type 2; OL, Oligodendrocytes; EC1, EC type 1; EC2, EC type 2; EC3, EC type 3; vEC, venous EC; capilEC, capillary EC; aEC, arterial EC; AC, Astrocyte.

**b,** Expression profile of *Tek* in the single-cell transcriptome of the human brain vasculature system (data sourced from a public database^69^). ART: Arterial; CAP, Capillary; VEN, Venous; aSMC, Vascular Smooth Muscle Cell; aaSMC, Arteriolar Smooth Muscle Cell; T-PC, Solute transport-Pericyte; M-PC, ECM-regulating Pericyte; P.FB, Perivascular Fibroblast; M.FB, Meningeal Fibroblast; TC, T cell; EPEN, Ependymal; AST-Hpc, Astrocyte-hippocampus; AST-Ctx, Astrocyte-Cortex; PM, Perivascular Macrophage; MG, Microglia; OL, Oligodendrocyte; OPC, Oligodendrocyte Precursor Cell; NEU, Neuron.

**Supplementary Figure 2**

**BLAST analysis of transcriptional regulatory sequences of *Tek* from different species and truncation analysis of miniBEND elements in the human gene version.**

**a,** DNA multiple sequence alignment analysis of the promoter region and 5’ untranslated region (5’ UTR) of *Tek* across multiple species, including human (*Homo sapiens*), macaque (*Macaca fasicularis*), marmoset (*Callithrix jacchus*), pig (*Sus scrofa*), cow (*Bos taurus*), rat (*Rattus norvegicus*), and mouse (*Mus musculus*).

**b,** Evolutionary tree analysis of sequences from different species, showing a closer evolutionary relationship for the promoter region sequences between pig and cattle, human and macaque/marmoset, and mouse and rat.

**c,** DNA multiple sequence alignment analysis of the genomic region corresponding to the first intron of

*Tek* across the same species as in (a).

**d,** Evolutionary tree analysis of sequences from different species, showing a closer evolutionary relationship for the first intron sequences between pig and cattle, human and macaque/marmoset, and mouse and rat.

**Supplementary Figure 3**

Truncation and *in vivo* characterization of the human *Tek* promoter and cis-element.

**a,b,** Schematic representation of the truncation strategy for the promoter (**a**) and the cis-element in intron 1 (b) of human *Tek*.

**c,** Representative images of cortical and hippocampal mouse tissue showing the restricted gene expression effect of truncated versions of the human *Tek*–derived rAAV-miniBEND system.

**d,** Assessment of combination of promoter and cis-element constructs from different species in mouse brains. Upper, hPro1612 (human) with marmoset-Cis700; middle, hPro1612 with pig-Cis700; lower, rat-Pro1600 with rat-Cis737.

**Supplementary Figure 4**

**Analysis of neuronal infection with AAV-PHP.eB-miniBEND-EGFP in adult mice.**

The infection of neurons was analyzed in the brains of adult mice (P60–90) following retro-orbital administration of AAV-PHP.eB-miniBEND-mPro723-EGFP viruses. Neurons were stained with antibodies against NeuN. Representative images showed that no EGFP-positive neurons were observed in the olfactory bulb, cerebral cortex, hippocampus, striatum, thalamus, or corpus callosum.

**Supplementary Figure 5**

**Analysis of astrocyte infection with AAV-PHP.eB-miniBEND-EGFP in adult mice.**

The infection of astrocytes was analyzed in the brains of adult mice (P60–90) following retro-orbital administration of AAV-PHP.eB-miniBEND-mPro723-EGFP viruses. Astrocytes were stained with antibodies against GFAP. Representative images showed that no EGFP-positive astrocytes were observed in the hippocampus (upper, CA1 region; lower, dentate gyrus).

**Supplementary Figure 6**

**Analysis of microglial infection with AAV-PHP.eB-miniBEND-EGFP in adult mice.** Microglial infection was analyzed in the brains of adult mice (P60–90) following retro-orbital administration of AAV-PHP.eB-miniBEND-mPro723-EGFP viruses. Microglia were stained with antibodies against Iba1. Representative images showed that no EGFP-positive microglia were observed in the olfactory bulb, cerebral cortex, hippocampus, striatum, or thalamus.

**Supplementary Figure 7**

**Analysis of pericyte infection with AAV-PHP.eB-miniBEND-EGFP in adult mice.**

The infection of pericytes was analyzed in the brain through retro-orbital administration of AAV-PHP.eB-miniBEND (mPro723-glo566-mCis700)-EGFP viruses to adult *Pdgfrb-Cre::Ai14* mice (P60–90).

Pericytes were labeled with tdTomato. Representative images showed that no tdTomato-positive pericytes were observed in the brain section.

**Supplementary Figure 8**

**rAAV-miniBEND system enables efficient and specific gene delivery to brainECs throughout the whole brain.**

**a,** Representative sagittal section image of the mouse brain demonstrating the efficient and specific transduction of brainECs in the whole brain, using the optimized mPro723-mCis700 version of the rAAV-miniBEND system in transgenic reporter mice *Ai14* after intravenous injection of AAV-PHP.eB-mPro723-Cre-mCis700. Boxed regions show four ROIs, which are shown at higher magnification in the lower images.

**b,** Co-localization of GluT1 staining (red) and EGFP fluorescence (green) in mouse brainECs labeled with the rAAV-miniBEND system. Results from cortex, hippocampus, and thalamus were obtained from *Ai47* mice injected with AAV-PHP.eB-miniBEND(mPro973)-Cre-mCis400. The cerebellum staining results are from *Ai47* mice injected with AAV-PHP.eB-miniBEND(mPro973)-Cre-mCis303. HO, nuclear staining with Hoechst 33342. White arrowhead, blood vessels.

**Supplementary Figure 9**

**Expression pattern of AAV-PHP.eB-Ple261-iCre in *Ai14* reporter line.**

The expression of tdTomato signals in brain sections from a P56 *Ai14* mouse after retro-orbital injection of AAV-PHP.eB-Ple261-iCre (50 μl, 6.5×10^12^ gc/ml). A high percentage of neurons and glial cells (arrowheads) were labeled in different brain regions, including the cerebral cortex, hippocampus, and thalamus.

**Supplementary Figure 10**

**rAAV-miniBEND system enables efficient gene delivery to retinal vascular endothelial cells.**

**a,** Representative images of retinas at 15 days after retro-orbital injection of rAAV-miniBEND for gene delivery to retinal vascular endothelial cells in a P46 mouse. For *NG2DsRedBAC::Ai47* double reporter mice, the red fluorescence from DsRed served as an aid in distinguishing arterial and venous segments in the retinal blood vasculature. AAV-miniBEND-Cre, AAV-PHP.eB-mPro973-Cre-mCis700; SMC, smooth muscle cell. White arrowhead indicates vascular smooth muscle cells in the left row and pericyte in the right row.

**b,** Schematic diagram of intravitreal injection of rAAV-miniBEND system for local gene delivery to vascular endothelial cells. Diagram was created using BioRender.com.

**c,** Representative images showing labeled endothelial cells in the retina of an *Ai* mouse at 14 days after injection (PID14) of 1.5 μl of AAV-9P13-miniBEND-Cre or AAV-PHP.eB-miniBEND-Cre. White arrowheads, vessels and abnormal vessels in 9P13-miniBEND-Cre (in *Ai14* mice) and PHP.eB-miniBEND-Cre (in *Ai47* mice) groups, respectively.

**d,** Representative images of retinas were captured at 21 days post retro-orbital injection of AAV-BI30 for gene delivery to retinal vascular endothelial cells in P3 rats. Specifically, AAV-BI30-Ef1α-EGFP virus was injected into wild-type rats, while AAV-BI30-miniBEND-Cre virus (AAV-BI30-mPro723-Cre-mCis700) was injected into transgenic rats Tg(*Rosa26-CAG-LSL-EGFP*).

The white arrowhead indicates neurons (the left panel) or vascular endothelial cells (the right panel).

**Supplementary Figure 11**

**Characterization of AAV-BI30 and AAV-BR1 capsid expression patterns in the adult mouse brain.** The expression patterns of EGFP in *Ai47* mice following retro-orbital injection of AAV-BI30-Ef1α-Cre viruses (upper) and AAV-BR1-Ef1α-Cre viruses (lower) are shown. Both viruses exhibit high infection efficiency in endothelial cells of the adult brain, but neurons were also labeled in the AAV-BR1-Ef1α-Cre group.

**Supplementary Figure 12**

Figure 1: Comparative analysis of AAV-BI30 and AAV-miniBEND-mediated gene expression in *Ai* reporter mice

**a**,Experimental workflow: *Ai14* reporter mice at P5–10 were injected with either AAV-BI30-CAG-Cre or AAV-PHP.eB-miniBEND(mPro723-mCis700)-Cre. Brain tissues were harvested and sagittally sectioned along the midline. Immunostaining was performed to visualize expression patterns. Additional analyses included cell isolation, FACS, and quantification. Diagram was created using BioRender.com.

**b**, Representative immunofluorescence images: Brain sections from *Ai14* mice injected with AAV-BI30-CAG-Cre (left) and AAV-PHP.eB-miniBEND-Cre (right) are shown. DAPI (blue) marks nuclei, while tdTomato (red) indicates Cre-induced fluorescence in reporter-positive cells. Enlarged insets (yellow boxes) highlight vascular structure and cellular localization. Scale bars: 1 mm (top row), 50 µm (bottom row).

**c**, Pie charts of CD31+ cell distribution: Comparison of CD31+ endothelial cell percentages between AAV-BI30-CAG-Cre and AAV-PHP.eB-miniBEND-Cre. AAV-miniBEND-Cre achieves nearly 100% CD31+ cell targeting (n=649 red cells), indicating higher endothelial specificity than the 75.1% (n=1268 red cells) achieved by AAV-BI30-CAG-Cre.

**d**, Flow cytometry plots: FACS analysis of brain samples from AAV-BI30-CAG-Cre (top row) and AAV-PHP.eB-miniBEND-Cre (bottom row) treated groups. Gated populations are shown for DAPI (nuclei), EGFP or tdTomato (Cre activity reporters), and CD31 (endothelial marker). *Ai + AAV-BI30-CAG-Cre* displays mixed EGFP/ tdTomato and CD31 expression, while *Ai + AAV-miniBEND-Cre* shows clear EGFP/tdTomato expression with a highly enriched CD31+ endothelial population.

**e**, Quantification of CD31+ cell percentages: Bar plot summarizing CD31+ cell proportions in both groups (BI30 and miniBEND), showing AAV-miniBEND-Cre’s superior specificity for endothelial cells. Data are presented as mean ± SEM, with a significant difference between groups (unpaired two-sided Welch’s t-test, P = 0.0038, \*\**p* < 0.01).

**Supplementary Figure 13**

**rAAV-miniBEND system enhances transduction specificity for brainECs with multiple AAV capsid serotype variants.**

Representative images of multiple brain regions show cell type–specific transduction in brainECs. AAV viruses were administered via retro-orbital injection. mPro723-mCis700, rAAV-mPro723-Cre-pA-mCis700 vector; AAV1-rh.10, AAV1-rh.10 capsid; AAV-DJ/8, AAV-DJ/8 capsid; AAV-9P13, AAV9-9P13 capsid; AAV-9P36, AAV9-9P36 capsid.

**Supplementary Figure 14**

**EGFP expression in peripheral organs of *Ai47* reporter mice infected with AAV-BI30-Cre.**

EGFP signals were measured in sections of different peripheral organs 20 days after intravenously administering AAV-BI30-Cre to *Ai47* reporter mice. Strong EGFP signals were detected in sections from the liver, spleen, and lung. No EGFP signals were detected in the vascular cells of the heart.

**Supplementary Figure 15**

**EGFP expression in peripheral organs of *Ai47* reporter mice infected with AAV-miniBEND-Cre.** EGFP signals were measured in sections of different peripheral organs 20 days after intravenously administering AAV-PHP.eB-miniBEND-Cre to *Ai47* reporter mice. Rare EGFP signals were detected in sections from the liver, lung, stomach, kidney, intestine, heart, and esophagus. However, very few vascular cells with EGFP expression were detected in sections from the spleen.

**Supplementary Figure 16**

**Distribution of AAV-miniBEND-EGFP vectors in different organs.**

AAV genome copies were measured in different organs (not endothelial cells) and quantified by qPCR using EGFP primers (n = 4 mice). Vector copy numbers were normalized to 100 ng of total DNA. Mice were retro-orbitally injected with AAV-PHP.eB-miniBEND (mPro723-glo566-mCis700)-EGFP viruses, and tissues were collected 20 days after virus injection. Tissues from five organs were analyzed: brain, heart, kidney, liver, and lung. DNA was subsequently extracted using the DNeasy tissue kit.

**Supplementary Figure 17**

**The rAAV-miniBEND system demonstrates enhanced brainEC specificity via local injection in adult and young mouse brains.**

**a**, AAV-BR1-Ef1α-Cre-P2A-EGFP was locally injected into the lateral septum or hippocampus (900 nl) of adult *Ai14* mice. White arrowheads, labeled neurons.

**b**, The mPro723 vector exhibits higher brainEC transduction efficiency and specificity as compared to the mPro973 group. AAV-PHP.eB-mPro973-Cre-mCis700 and AAV-PHP.eB-mPro723-Cre-mCis700 were locally injected (1 μl) into the indicated region of *Ai14* (left) and *Ai47* (right) mice at the indicated ages. PFC, prefrontal cortex. White arrowheads, labeled neurons and brain ECs.

**c**, The rAAV-miniBEND system demonstrates reduced brainEC specificity via local injection in older mice. AAV-PHP.eB-mPro723-Cre-mCis700 was locally injected (1 μl) into the indicated region of *Ai47* and *Ai14* mice at the indicated ages. White arrowheads, labeled brainECs.

**Supplementary Figure 18**

rAAV-miniBEND enables highly efficient gene delivery to brainECs through *in utero* intracerebroventricular (i.c.v.) injection.

**a**, Schematic diagram illustrating *in utero* i.c.v. injection of AAVs. Diagram was created using BioRender.com.

**b**, Representative images showing endothelial labeling efficiency in different brain regions at PID16 after injection with AAV-BR1-CAG-mGL, AAV-PHP.eB-CAG-mScarlet, AAV-PHP.V1-miniBEND (mPro1576-mCis700)-EGFP, and AAV-BI30-miniBEND (mPro723-mCis700)-Cre in wild-type (WT) or *Ai47* mouse strains. Blue arrowheads, representative neurons (a’–d’, f’–h’, j’) or endothelial cells (e’, i’, n’, o’–r’) that were labeled.

**c**, The transduction efficiency of AAV-BI30-miniBEND (mPro723-mCis700)-Cre for brainECs in different brain regions. n = 3 mice, with 5 ROIs analyzed for each brain region from each mouse.

**Supplementary Figure 19**

**The rAAV-miniBEND system facilitates highly efficient gene delivery to rat brainECs through retro-orbital injection.**

Representative images show the endothelial labeling efficiency and specificity in various brain regions at PID21 following injection with AAV-X1.1-Ef1α-Cre, AAV-BI30-Ef1α-EGFP, and AAV-BI30- miniBEND (mPro723-mCis700)-Cre in both wild-type (WT) and transgenic rat strains, *Tg(Rosa26-CAG-LSL-EGFP)*.

**Supplementary Figure 20**

**Measurement of the duration of gene expression mediated by rAAV-miniBEND system.**

**a.** Representative images of EGFP expression in brain endothelial cells mediated by the rAAV-miniBEND system. Wild-type mice were injected retro-orbitally with AAV-PHP.eB-mPro723-glo566-EGFP-pA-mCis700 viruses. Brains were harvested on PID22 (post-injection day 22) and PID90 for subsequent sectioning and imaging.
**b.** Summary of results showing the density of endothelial cells with EGFP expression on PID22 (n = 2), PID32 (n = 2), PID60 (n = 2), and PID90 (n = 2 mice). We selected 5-7 ROIs per mouse for analysis. The duration of gene expression was quantified by measuring the length density of EGFP^+^ vessels in a limited brain region as shown in (**a**).

**Supplementary Figure 21**

**Evaluation of labeling efficiency of rAAV-miniBEND system in the mouse brain**

**a.** Adult mice (P60–90) were administered AAV-PHP.eB-miniBEND-S-(mPro723-glo228-mCis700)-EGFP viruses (8.7×10^12^ gc/ml, 100μl). The expression efficiency was evaluated by calculating the percentage of EGFP-positive cells (green) among all endothelial cells. Blood vessels were labeled using antibodies against Collagen IV (red).
**b.** Summarized results of the percentage calculated from multiple brain regions (8 ROIs from 2 mouse brains). The regions includes Cerebellum (CB), Cortex (CTX), Hippocampus (HIP) and Olfactory bulb (OB).

**Supplementary Figure 22**

**Pattern of regressive blood vessels in *Tak1^fl/fl^* mice infected with AAV-PHP.eB-miniBEND-Cre.**

**a**, **b**. Representative images (**a**) and a summary of the density (**b**) of regressive blood vessels (arrowheads) in *Tak1^fl/fl^* mice after administration of PBS (top row, **a**) or AAV-PHP.eB-miniBEND (mPro723-mCis700)-Cre (bottom row, **a**). 12 regions from 2 mice in each group were selected for analysis, unpaired two-sided Welch’s t-test, P < 0.0001, \*\*\*\**p*<0.0001.

**Supplementary Figure 23**

**Characterization of sporadic CCM lesions induced by human *MAP3K3^I441M^* via rAAV-miniBEND system.**

**a.** H&E staining of a sagittal brain section from a wild-type mouse injected retro-orbitally with AAV-miniBEND-S (mPro723-glo566-mCis700)-hMAP3K3^I441M^ viruses.
**b.** Leakage of red blood cells (upper panel, RBC lesion, arrowheads) and enlarged vessels (lower panel, caverns, arrowheads) were observed in the lesion area.

**Supplementary Figure 24**

**Characterization of sporadic CCM lesions induced by human *MAP3K3^I441M^* via rAAV-miniBEND system.**

Wild-type mice were retro-orbitally injected with AAV-PHP.eB-miniBEND-S (mPro723-glo566-mCis700)-hMAP3K3^I441M^ viruses, followed by subsequent injection with Dextran-FITC-70KDa and Lectin-DyLight594 dye before perfusion. Blood occlusion and leakage of the blood-brain barrier (BBB) were observed in these cavernous vessels. In the left panel, cavernous vessels are indicated by arrowheads, while in the right panel, leaking blood vessels from the cavern are indicated by arrowheads.

**Supplementary Figure 25**

**H&E and Prussian blue staining of brain sections with CCM lesions induced by human MAP3K3^I441M^.**

Brain sections were collected from mice injected with AAV-miniBEND-S (mPro723-glo566-mCis700)-MAP3K3^I441M^. Leakage of blood vessels in the lesion was observed in sections stained with H&E (upper) and Prussian blue (lower). Red blood cells (upper, arrowheads) were detected using H&E staining in the perivascular spaces of the lesioned areas in different brain regions (cerebral cortex, hippocampus, midbrain, and cerebellum). Hemosiderin deposition from Prussian blue staining (lower, arrowheads) was also observed in the CCM lesion areas of these brain regions.

**Supplementary Figure 26**

**P62 staining in lesions with human *MAP3K3^I441M^* expression.**

Representative images of staining with antibodies against p62 (green) and Laminin (red) from the control and brain region infected with AAV-PHP.eB-miniBEND-S (mPro723-glo560-mCis700)-hMAP3K3^I441M^ viruses. Blue indicates nuclei stained with Hoechst33342 (Hoechst).

**Supplementary Figure 27**

**CCM lesions in brain sections from mice infected with AAV-miniBEND-MAP3K3^I441M^.**

**a, b.** Representative images of CCM lesions in the hippocampus and cerebellum. Insets in **a** and **b** show four lesions from these two brain regions.

**a. c.** Summary of lesion density across the whole brain section.

**d, e.** Summary of the diameters and areas of CCM lesions in whole brain sections of mice infected with AAV-miniBEND-S-MAP3K3^I441M^.

**Supplementary Figure 28**

**Distribution of CCM lesions mediated by *MAP3K3^I441M^* expression across the whole brain.**

**a.** Representative MRI images of CCM lesions from a mouse injected retro-orbitally with AAV-miniBEND-S (mPro273-glo566-mCis700)-hMAP3K3^I441M^ (1×10^10^vg). The brains were imaged 7 days (left, PID7) and 22 days (right, PID22) after virus injection.
**b.** Summary of CCM lesion density across different brain regions (n = 4 mice). The regions include: BS (brain stem), CB (cerebellum), CTX (cerebral cortex), HIP (hippocampus), OB (olfactory bulb), and TH/MB/HY (thalamus, midbrain, and hypothalamus).

**Supplementary Figure 29**

**Abnormalities of intracranial vascular structure mediated in *Braf-CA-fl/fl* mice after administration of the AAV-miniBEND-Cre viruses.**

H&E Staining: Malformed vessels in *Braf-CA^E^-fl/fl* mice were observed after local injection of the AAV-PHP.eB-miniBEND (mPro723-mCis700)-Cre viruses (n = 2 mice). The vessels showed enlarged vascular lumens (ROI1) and hemorrhage (ROI2).

**Supplementary Figure 30**

**Lesions in brain sections of *Braf-CA****-fl/fl* **mice infected with AAV-miniBEND-Cre.**

**a, b.** Representative images of lesions in the hippocampus and cerebellum. Insets in **a** and **b** show four lesions from these two brain regions.

a. **c.** Summary of lesion density across the whole brain section.

**d, e.** Summary of the diameters and areas of CCM lesions in whole brain sections of *Braf-CA-fl/fl* mice infected with AAV-miniBEND-Cre.

**Supplementary Figure 31**

**Detection of cellular apoptosis in brain regions with lesions in the mouse model of bAVM.**

**a–d,** In the brain sections with bAVM lesions, TUNEL staining, CD31 staining (a,b), and NeuN staining (c,d) were performed to assess endothelial cell or neuronal apoptosis in the region with AVM lesion and the control region from the contralateral hemisphere.

TUNEL staining (red) identifies apoptotic cells, CD31 staining (green) was used to label endothelial cells, and NeuN staining (green) was used to label neurons. Nuclei (blue) were stained with Hoechst 33342. For both group, five regions were selected for cell counting, unpaired two-sided Welch’s *t*-test, P = 0.0053 (b), P = 0.0011 (d), *p*<0.01**.

**Supplementary Figure 32**

**Laser speckle blood flow imaging of the Braf-bAVM mouse model.**

**a**, Bright-field (upper) and laser speckle flowmetry (lower) images from a *Braf-CA*^fl/fl^*::Ai14* mouse with a cranial window at the indicated time points. The color scale indicates the range of blood flow: red for maximum flow and blue for minimum flow.

**b-d**, Blood flow index analysis of the ROI1 (b), ROI2 (**c**), and ROI3 (d) as shown in (a). PU, perfusion units. Data indicate the mean ± s.e.m., with individual data points shown as well. unpaired two-sided Welch’s t-test, P < 0.0001, \*\*\*\**p* < 0.0001.

**Supplementary Figure 33**

**Abnormalities of intracranial vascular structure mediated in *Braf-CA-fl/fl* mice after administration of the AAV-miniBEND-Cre viruses.**

Two-photon excitation microscopy: Images taken using two-photon excitation microscopy showed arteriovenous shunts in *Braf-CA-fl/fl::Ai14* mice induced by the injection of AAV-PHP.eB-miniBEND (mPro723-mCis700)-Cre. The malformed vessels displayed enlargement in segments and abnormal blood flow (arrowheads, also see Supplementary Video 3 and 4,). Red signals indicate tdTomato (tdT) expression, and green signals indicate Dextran-FITC.

**Supplementary Figure 34**

**Representative images of malformed vasculature in the brain of a *Braf-CA-fl/fl* mouse perfused with latex dye.**

a. Whole-brain and high-magnification images of brain vasculature in a *Braf-CA^fl/fl^* mouse following local injection of AAV-miniBEND-Cre into the right cortex. The left hemisphere received control viruses (AAV-miniBEND-EGFP) without Cre. Malformed vessels are indicated by arrowheads.
b. Whole-brain and high-magnification images of latex dye-perfused brain vasculature in a control mouse and a *Braf-CA^fl/fl^* mouse after retro-orbital venous injection of AAV-miniBEND-Cre. Malformed vessels are indicated by arrowheads.

**Supplementary Figure 35**

**Images of whole-brain vasculature and malformed vessels from the cleared brain of a *Braf-CA-fl/fl::Ai14* mouse infected with AAV-miniBEND-Cre.**

**a)** Schematic illustrating the AAV injection process and whole-brain clearing method using PEGASOS. Whole-brain vasculature was visualized with light-sheet microscopy. A full view of the whole-brain vasculature image is shown at right. Diagram was created using BioRender.com.
**b)** Representative regions with malformed vessels indicated by arrowheads. Two sample frames and a 3D image from the corresponding brain block are included. Three videos derived from these images are included in the supplementary materials.

**Supplementary Figure 36**

**Confocal imaging of malformed vessels in a cleared brain block from a *Braf-CA^fl/fl^::Ai14* mouse.**

A *Braf-CA^fl/fl^::Ai14* mouse was intravenously injected with AAV-PHP.eB-miniBEND-Cre, and its whole brain was cleared using the PEGASOS protocol. A 1 mm thick block of this cleared brain was imaged with confocal microscopy, enabling high-resolution visualization of the malformed vessels. Images show three representative malformed vessels (labeled b–d, arrows) which were selected from a section of the cleared brain.

**Supplementary Figure 37**

**Activation of astrocytes in lesions induced by *Braf^V600E^* after administration of AAV-PHP.eB-miniBEND-Cre.**

**a.** Staining of control and nidus region with antibodies against GFAP and Laminin. Nuclei were labeled with Hoechst33342. *Braf-CA-fl/fl* mice were injected with AAV-PHP.eB-miniBEND (mPro723-mCis700)-Cre (local injection, cortex, unilateral; 900 nl; PID76).
**b.** Representative morphology of astrocytes in control and nidus region. Normal: normal control brain area, Nidus: lesion area.
**c.** Summary of GFAP-positive area. Astrocytes were significantly activated in the lesion area. GFAP, astrocytes. 10 regions in each group were selected for analysis, unpaired two-sided Welch’s t-test, P = 0.0031, \*\**p*<0.01.

**Supplementary Figure 38**

**Activation of microglia in lesions induced by *Braf^V600E^* after administration of AAV-PHP.eB-miniBEND-Cre.**

**a.** Staining of control and nidus region with antibodies against Iba1 and Laminin. Nuclei were labeled with Hoechst33342. *Braf-CA^fl/fl^* mice were injected with AAV-PHP.eB-miniBEND (mPro723-mCis700)-Cre (local injection, cortex, unilateral; 900 nl; PID76).
**b.** Representative morphology of microglia in control and nidus region. Normal: normal control brain area, Nidus: lesion area.

**Supplementary Figure 39**

**Measurement of lesions in bAVM mice treated with PLX4032.**

**a, b.** Representative images of H&E staining of brain sections from the cerebral cortex of *Braf-CA-fl/fl* mice after local injection of AAV-PHP.eB-miniBEND-Cre. **a** shows the control group (DMSO), while **b** shows the PLX4032-treated group. Insets show two representative lesions from the control group (**a**) and the treated group (**b**).

**c, d.** Density (c) and diameter (d) of malformed vessels in the control group and the PLX4032-treated group . For c, five regions were selected for analysis; for d, DMSO group (n = 561 vessels), PLX4032-treated group (n = 64 vessels), unpaired two-sided Welch’s t-test, two-tailed P = 0.0013 (c), P < 0.0001 (d), \*\**p* < 0.01, \*\*\*\**p* < 0.0001.

**Supplementary Figure 40**

**The EndMT process in the Braf-bAVM mouse model.**

**a,** GSEA was performed with the EndMT gene set (which is included in Epithelial-Mesenchymal Transition Geneset) on endothelial cell (EC) expression data from the Braf-bAVM mouse model and human bAVM single-cell transcriptomic data. FDR, false discovery rate; NES, normalized enrichment score. The y-axis represents the enrichment score of the Gene Set Enrichment Analysis (GSEA) result. GSEA was performed using permutation testing, two-sided P values were calculated, and FDR correction was applied.

**b,** Immunofluorescence co-staining images depict the expression of αSMA and CD31 in control brain regions and nidus regions in a PID76 mouse brain. HO, nuclear staining with Hoechst 33342.

**Supplementary Video 1**

**The Braf-bAVM mouse model exhibits hemiplegic seizures.**

This video corresponds to Figure 4a and b. *Braf-CA*^fl/fl^ mice, after receiving a local injection of the AAV-PHP.eB-miniBEND-Cre viruses (PID32) in the cortex and displaying abnormal behavior, were placed in an open field and recorded for ∼1 hour. This video clip serves as a representative example for Braf-bAVM mice, with a hemiplegic seizure rate of 13 of 21 mice (61.9%).

**Supplementary Video 2**

**Three-dimensional whole brain visualization of a bAVM nidus.**

This video corresponds to Figure 4a and b but we performed local injection of AAV-PHP.eB-miniBEND-Cre viruses in *Braf-CA*^fl/wt^*::Ai14* mice. Mice were then subjected to retro-orbital injection of Lectin-DyLight 488 to label blood vessels before fixation with paraformaldehyde. Subsequently, the brain was dissected and processed using the PEGASOS tissue clearing procedure. Whole-brain imaging of the cleared brain was conducted using lightsheet microscopy, allowing for comprehensive visualization of the bAVM nidus in three dimensions.

**Supplementary Video 3**

**Observation of arteriovenous shunts in the abnormal vessels in bAVM lesions under two-photon excitation microscopy**.

AAV-PHP.eB-mPro723-Cre-mCis700 viruses (800 nl) were locally injected into *Braf-CA^fl/fl^::Ai14* transgenic mice and imaged at PID 18. Blood flow is illustrated with FITC-Dextran (20 kDa), and the red signal indicates endothelial cells with tdTomato expression.

**Supplementary Video 4**

**Observation of arteriovenous shunts in the abnormal vessels in bAVM lesions under two-photon excitation microscopy**.

AAV-PHP.eB-mPro723-Cre-mCis700 viruses (800 nl) were locally injected into *Braf-CA^fl/fl^* mice and imaged at PID18. Blood flow was illustrated with FITC-Dextran (20kDa). No leakage was detected in the region we imaged.

**Supplementary Video 5**

**Representative images of malformed vessels from the cleared brain of a *Braf-CA-fl/fl::Ai14* mouse infected with AAV-PHP.eB-miniBEND-Cre.**

Whole-brain vasculature was visualized with light-sheet microscopy. A 3D image of one representative region with malformed vessels from the corresponding brain block in Supplementary Figure 35.

**Supplementary Video 6**

**Representative images of malformed vessels from the cleared brain of a *Braf-CA-fl/fl::Ai14* mouse infected with AAV-PHP.eB-miniBEND-Cre.**

Whole-brain vasculature was visualized with light-sheet microscopy. A stack image of one representative region with malformed vessels from the corresponding brain block in Supplementary Figure 35.

**Supplementary Video 7**

**Representative images of malformed vessels from the cleared brain of a *Braf-CA-fl/fl::Ai14* mouse infected with AAV-PHP.eB-miniBEND-Cre.**

Whole-brain vasculature was visualized with light-sheet microscopy. A stack image of one representative region with malformed vessels from the corresponding brain block in Supplementary Figure 35.

**Supplementary Video 8**

**Representative images of malformed vessels from the cleared brain of a *Braf-CA-fl/fl::Ai14* mouse infected with AAV-PHP.eB-miniBEND-Cre.**

Whole-brain vasculature was visualized with light-sheet microscopy. A stack image of one representative region with malformed vessels from the corresponding brain block in Supplementary Figure 35.

## Methods

### Animals

Adult or pregnant C57BL/6 mice were purchased from the Beijing Vital River Laboratory Animal Technology (stock #027). Rats and other transgenic mice were bred in our laboratory. Animals were housed in a 12-h light/12-h dark environment and were provided food and water *ad libitum*. Ambient temperature was kept at 25 °C with humidity at 50%. All animals were euthanized using Avertin or isoflurane (5%). All procedures for animal surgery and maintenance were in accordance with protocols approved by the Institutional Animal Care and Use Committee at the Chinese Institute for Brain Research (CIBR, Beijing, China). *NG2BacDsRedtg* mice were originally from Akiko Nishiyama’s laboratory at the University of Connecticut (also available at The Jackson Laboratory, Cat# 008241). *Pdgfr*β*-Cre* mice were originally from Volkhard Lindner’s laboratory (MaineHealth Institute for Research, Scarborough, United States). *Cdh5-CreER* mice were acquired from Ralf H. Adams’s laboratory (University of Münster, Germany). *Ai14* mice (Cat# 007914) were obtained from The Jackson Laboratory. *Ai47* mice were acquired from the laboratory of Hongkui Zeng (Allen Brain Institute). Adult mice (10–16 weeks) of either gender were used for virus injection. Pregnant mice were used for embryonic lateral ventricle virus injection at the indicated time points. The wild-type rat (Sprague-Dawley) and the transgenic reporter rat (*Rosa26-CAG-LSL-EGFP-pA*, with a Sprague-Dawley background) strain were supplied by the Genetic Manipulation Core Facility at the Chinese Institute for Brain Research.

For the *Braf-CA* transgenic mice (The Jackson Laboratory, Cat# 017837), a homologous recombination knock-in vector was designed to preserve the normal expression of endogenous *Braf* and production of intact BRAF protein sequence, albeit with human exons 15–18. It contains a LoxP site, a DNA fragment from exon 15 to exon 18 of human *BRAF*, the mouse *Braf* polyA element, and a *Neo* selection cassette, followed by the second LoxP site. This second LoxP site is upstream of the exon 15 coding region of mouse *Braf*, in which nucleotide 1799 has been mutated from a T to an A (c.1799 T to A), followed by mouse exons 16–18. This results in a valine (V) to glutamic acid (E) mutation at amino acid 600 in the encoded protein in the presence of Cre recombinase activity.

### Plasmid preparation

Molecular cloning was carried out using the ClonExpress II One Step Cloning Kit (Vazyme, C112-01) with oligo primers of ∼20 overlapping bases (GENEWIZ, Azenta) and DNA polymerase (FastPfu, TransGen; Phanta Mix, Vazyme; or PrimeSTAR, Takara Bio) and was verified by Sanger sequencing (Tsingke, Beijing). A standardized rAAV expression vector (rAAV-Promoter-Cre-BGHpA-Cis-element) was designed and constructed for this study (Fig. 1a). The upstream and downstream regions of the promoter were flanked by restriction enzyme sites *Spe*I and *Age*I to allow replacement of truncated versions of the *Tek* promoter. After the Kozak sequence, the NLS-Cre-HA gene expression cassette was inserted, followed by the polyA element BGHpA. Downstream of the polyA element, truncated versions of the cis-regulatory element of interest were individually introduced with single restriction enzyme sites *Kpn*I and *Not*I at either end, which facilitated molecular cloning. The chimeric miniBEND promoter was generated by fusion polymerase chain reaction (PCR) of the mPro1576 fragment and cmv-globin intron fragments. *MAP3K3*^I441M^ cDNA fragments were clone from a cDNA library of HEK293T cells, point-mutated through a PCR strategy, and subcloned into rAAV-miniBEND vector (Fig. 3d). mGL refers to the mGreenLantern fluorescent protein^70^.

### Regulatory region selection and promoter sequence retrieval

The UCSC Genome Browser (http://genome.ucsc.edu/) can be used to browse, extract, and compare genomic sequences of many model species, including the species mentioned in this study. Multiple sequence alignment and evolutionary tree analysis was conducted through “Align” module with ClustalW algorithm in Vector NTI.

### rAAV packaging and viral titration

rAAVs were packaged in the vector core facility at CIBR. The AAV vectors were packaged as described^50, 71–73^. In brief, the AAV vectors and the AAV helper plasmids were co-transfected into HEK293T cells. Cells were collected 96 h after transfection, and viral particles were released from cells by freeze–thaw cycling and sonication. The virus was purified using cesium chloride density-gradient ultracentrifugation and was dialyzed against phosphate-buffered saline (PBS). The viral titer was analyzed using quantitative PCR (qPCR)^74^.

For qPCR viral titration, we first generated a standard curve using a linearized plasmid DNA template. A suitable restriction enzyme site was chosen, and the digested plasmid DNA was purified, its concentration measured, and the copy number calculated. Tenfold serial dilutions of the template were prepared to establish a five-point standard curve.

For rAAV samples, 49.5 µl of lysis buffer was added to 0.5 µl of each rAAV preparation, then lysed at 98°C for 10 minutes, followed by addition of 49.5 µl of neutralization buffer. Using the same primers for both the standard curve and rAAV sample, qPCR was performed with Hieff® qPCR SYBRGreen Master Mix (No Rox) (YEASEN Biotechnology, China). The qPCR program was as follows: an initial cycle of 95°C for 5 minutes, followed by 40 cycles of 95°C for 10 seconds, 55°C for 20 seconds, and 72°C for 20 seconds. The reactions were conducted using a BioRad CFX96 Real Time PCR Thermal Cycler.

### Tamoxifen preparation and treatment

Tamoxifen was dissolved with corn oil (10 mg/ml). Mice were administered tamoxifen (100 mg/kg body weight per day) for 3 consecutive days after two-photon live imaging of basal phase.

### Surgical procedures and virus injection

We used an established method for *in utero* intracerebroventricular (i.c.v.) injection of AAVs^75^. Briefly, pregnant C57BL/6 or *Ai47* mice at 15–18 days gestation were placed under isoflurane anesthesia, and a midline laparotomy was conducted to expose the uterus. Generally, ∼1 μl of AAVs was administered to each fetus via transuterine injection directed toward the anterior horn of the lateral ventricle on the right side of the brain (Extended Data Fig. 9).

We also used established protocols for lateral ventricle injection of postnatal mice and retro-orbital injection of adolescent or adult mice^76^. Briefly, for the latter, the injection location corresponded to the lateral canthus of the left orbit. A 3/10-ml insulin syringe attached to a 31-G needle was gently inserted into the mouse’s orbital venous sinus to a depth of ∼2–3 mm, with the needle’s bevel pointing upward at a 45^°^angle. For stereotaxic injection into the lateral ventricle, mice were anesthetized with a gas mixture containing 5% isoflurane and 95% oxygen and then were placed on a stereotaxic instrument (Keaotong, China). This stereotaxic instrument has a mouse mask that is connected to an anesthesia machine (RWD, China). Injections were performed using a microsyringe pump and a commercial controller (Keaotong, China).

For AAV injection was performed with a stereomicroscope (Leica M125C) during the surgical procedure, and the AAV-miniBEND-Cre virus was injected into the subdural space at a depth of 0.4 mm during the cranial window surgery, with the injection site located in the center-right region of the cranial window.

Intravitreal injection into adult mice adhered to an established protocol (Extended Data Fig. 5b)^38^. Briefly, mice were anesthetized with isoflurane and placed on a stereotaxic instrument. For pupil dilation, Tropicamide-Phenylephrine ophthalmic solution (SANTEN Pharmaceutical, China) was used. An incision was made into the sclera (1–2 mm posterior to the superior limbus) using a sterile, sharp 31-G needle (insulin syringe, BD). After removing the 31-G needle, a blunt needle attached to a microliter Hamilton syringe (10 µl) was carefully inserted into the same incision and 1–1.5 µl of viral suspension was slowly injected into the vitreous body.

### Assessing BBB breakdown and quantification

The integrity of the blood–brain barrier (BBB) was determined by quantification of hemorrhagic lesion points. Each mouse received an injection of FITC-dextran at a rate of 10 μl/g body weight. After 5 min, the brains were dissected and preserved in preparation for whole-mount or sectional fluorescence imaging. Hemorrhagic lesions, characterized by distinct green fluorescent dye leakage, were selected and counted. The density of hemorrhagic lesions was then determined for various brain regions, including the olfactory bulb, cortex, hypothalamus, and cerebellum. In some experiments, BBB leakage or hemorrhage were also detected by Prussian blue staining or through detecting red blood cells in extra vessel spaces and identifying hemosiderin deposition on H&E-stained sections.

### Flow cytometry of brain cells

Mice aged P30–45 were anesthetized with isoflurane and perfused with precooled PBS (HyClone) via the heart. Brains were isolated and immediately immersed in ice-cold HBSS (5 mM KCl, 5 mM NaOH, 5 mM NaH[PO[, 0.5 mM MgCl[, 20 mM sodium pyruvate, 5.5 mM glucose, 200 mM sorbitol, pH 7.3– 7.4, filtered with a 0.22 μm filter). Each brain hemisphere was diced into small pieces and incubated in HBSS with 2 mg/ml papain (Sigma) and 5 μg/ml DNase I (Sigma) at 37°C for 30 minutes. A fire-polished Pasteur pipette (BrainBits) was used to dissociate the tissue into a single-cell suspension.

The cell suspension was then added to an Optiprep gradient^77^ and centrifuged at 1000 g for 15 minutes. The top debris layer was removed, and specific layers were collected. Cells from each layer were washed with HBSS and pelleted at 700 g for 10 minutes. After discarding the supernatant, cells were incubated in 1 ml of HBSS containing Alexa Fluor® 647 Rat Anti-Mouse CD144 (BD) at room temperature for 40 minutes in the dark. The cells were then centrifuged at 300 g for 5 minutes, the supernatant was discarded, and the cells were resuspended in 1 ml HBSS for sorting.

### Single-cell RNAseq library construction and sequencing

After brains from our bAVM model mice were harvested, cell count and viability was estimated using a fluorescence Cell Analyzer (Countstar Rigel S2). We used the contralateral control tissue as the control samples. Fresh cells were resuspended at 1 × 10^6^ cells/ml in PBS containing 0.04% bovine serum albumin. Single-cell RNAseq libraries were prepared using SeekOne MM Single Cell 3’ library preparation kit (SeekGene, Cat# SO01V3.1). Briefly, an appropriate number of cells was loaded into the flow channel of a SeekOne MM chip, which has 170,000 microwells, and the cells were allowed to settle into the microwells by gravity. After the unsettled cells were removed, sufficient Cell Barcoded Magnetic Beads (CBBs) were pipetted into the flow channel and were also allowed to settle into the microwells with the help of a magnetic field. Next, excess CBBs were rinsed out, and the cells in the MM chip microwells were lysed to release their RNA, which was captured by the CBB in each microwell. Then all CBBs were collected and reverse transcription was performed at 37 °C for 30 min to label the cDNAs with cell barcodes on the beads. Further Exonuclease I treatment removed unused primer on the CBBs. Subsequently, barcoded cDNA on the CBBs was hybridized with random primer, which had read 2 SeqPrimer sequence on the 5’ end and could extend to form the second-strand cDNA with the cell barcode on its 3’ end. The resulting double-stranded cDNA was denatured to release the CBBs, was purified, and then was amplified in a PCR reaction. The amplified cDNA product was then cleaned to remove unwanted fragments and was added to a full-length sequencing adapter and sample index by indexed PCR. The indexed sequencing libraries were cleaned up with SPRI beads, quantified by qPCR (KAPA Biosystems KK4824), and then sequenced on an Illumina NovaSeq 6000 with a paired-end read length of 150 bp.

### Single-cell RNAseq data analysis

Single-cell sequencing data obtained pre-processed data of AAV-miniBEND and *Braf-CA* transgenic mice based focal bAVM disease model. Fastq files were processed with the commercial scRNAseq data analysis software SeekSoulTools (SeekGene) and were organized into standard expression matrices according to online manuals (http://seeksoul.seekgene.com/en/v1.2.0/2.tutorial/1.rna.html). The single-cell transcriptome expression matrix data were converted into a Seurat object by the R package Seurat (version 4.1.3)^78^ using the software RStudio Desktop v2022.02.3 (Posit Software, PBC) and then were filtered to remove cells with <200 or >4000 unique gene counts. Cells with >5% mitochondrial gene counts were also removed. The global-scaling normalization method “LogNormalize” was then used to normalize the feature expression measurements for each cell relative to the total expression; this was multiplied by a scale factor of 10,000, and the result was log-transformed. Over 95% of high-quality cells with >25,000 protein-coding genes in total remained after the filtering and normalization for downstream processing.

### Identification of cell types and gene expression by dimension reduction

A standard analysis workflow was carried out as per the Seurat manual available online (https://satijalab.org/seurat/). In brief, NormalizeData() was used with the parameter “normalization.method” set to “LogNormalize” and “scalefactor” set to 10,000. After the normalization, subsets of feature genes were generated that exhibited high cell-to-cell variation in the dataset. Data normalization is performed after the NormalizeData() step. The ScaleData() function confines the data distribution to a specified range for easier comparison. Next, a linear transformation (scaling data) and subsequent linear dimensional reduction were performed. Then, cells were clustered by feature gene profiles. The annotation of each cell cluster was confirmed by the expression of canonical marker genes such as *Snap25*, *Syt1*, *Slc32a1*, and *Slc17a7* for neurons; *Flt1*, *Erg*, and *Cldh5* for endothelial cells; *Cx3cr1*, *Tmem119*, and *C1qa* for microglia; *Gfap*, *Slc1a3*, and *Aldh1l1* for astrocytes; and *Oligo1*, *Mbp*, and *Plp1* for oligodendrocytes^79^. The “resolution” parameter was set to “0.5”, which was determined by the modular optimizer developed by Ludo Waltman and Nees Jan Van Eck (Modularity Optimizer v1.3.0) . Finally, dimension reduction was performed on the single-cell gene expression matrix using the RunUMAP() function.

### Differential expression analysis and GSEA analysis

Differential expression analysis was performed using DESeq2^80^ with a cutoff of FDR < 0.05 and **|**log_2_(fold change)**|** > 1. Two-sided Wald test was used, and P values were adjusted for multiple testing using the Benjamini-Hochberg method (adjusted P < 0.05). Volcano plots, scatter plots, and heatmaps were generated using R packages (ggplot2; pheatmap) implemented in RStudio Desktop. For gene set enrichment analysis (GSEA)^81^, we generated a KEGG_2019_Mouse geneset based on a database file from the Enrichr online library (https://amp.pharm.mssm.edu/Enrichr/)^82^. For GSEA analysis in Supplementary Figure 37, we downloaded the Seurat file containing merged EC clusters from https://cells.ucsc.edu/?ds=adult-brain-vasc. Differentially expressed genes from malformed and normal human bAVM EC cell clusters were analyzed using the R Seurat package. Genes were pre-ranked through the metrics algorithm log_2_(fold change) × –log_10_(*p*-value[not adjusted *p*-val]) according to the statistical results from DESeq2. A pre-ranked (.rnk) file and custom geneset were used as input for GSEA v4.0.3 (https://www.gsea-msigdb.org/gsea/index.jsp). The number of permutations was set at 1000, and enrichment statistics were set at “weighted”. For the general significance threshold, FDR q-val < 0.25 and |NES| > 1.5 were considered as significant enrichment.

### Laser speckle blood flow imaging

The laser speckle blood flow imaging system (Wuhan SIM Opto-technology Co., Wuhan, China) consists of an Olympus ZS61 microscope, a continuous-wavelength laser diode (k = 785 nm), a charge-coupled-device camera, and a computer. The observation region was illuminated using the continuous-wavelength laser light source. Speckle signals were continuously acquired by the camera with a 10-ms exposure time and were subsequently transmitted to the computer for downstream analysis. The region of interest (ROI) was selected by a tool provided by the computer software, and the value obtained was the mean cerebral blood flow (CBF) in the region. Three ROIs were selected for each mouse, and the CBF was recorded for 100 cycles.

### Magnetic resonance imaging (MRI) of mice

We performed MRI as described^50^. Briefly, coronal T2-weighted magnetic resonance images were collected on a Bruker 7.0 T MRI scanner (Bruker Pharmascan 70/16) using 43.5–45 MHz, a 23-mm surface coil, and a 12-cm (in diameter) self-shielded gradient system for acquiring signals. The user interface was Paravision 5.1 software (Bruker BioSpin) and a Linux PC running Topspin 2.0. A mixture of 5% isoflurane and 95% O_2_ was used to induce anesthesia in a Plexiglas chamber, and 2% isoflurane/98% O_2_ was used for maintenance. The respiratory rate was closely monitored by an animal physiological guarding system. T2-weighted imaging was performed with a rapid acquisition method with relaxation enhancement (RARE). The detailed parameters were as follows: repetition time, 3500 ms; echo time, 33 ms; RARE factor, 4; field of view, 21 × 21 mm; acquisition matrix, 256 × 256; and slice thickness, 0.5 mm. The number of lesions and area of focal regions were quantified from the T2-weighted images with 3D Slicer(v4.11). We also used FLASH sequence for imaging in some mice. The imaging parameters for the FLASH sequence were as follows: Field of View (FOV) = 2.5cm × 2.5cm, Matrix (MATRX) = 256 × 256, Repetition Time (TR) = 600ms, Echo Time (TE) = 8ms, Flip Angle = 40deg, Slice Thickness = 0.5mm, Slice Distance = 0mm, Number of Slices = 25, Acquisition Time (TA) = 5m45s, Number of Averages = 3.

### Cranial window surgery and two-photon live imaging

Mice were anaesthetized with 1–2% isoflurane in oxygen delivered by an anesthetic machine (RWD Life Science Co., Shenzhen, China). We performed a craniotomy (3 mm in diameter) over the cerebral cortex, centered 2–3 mm posterior to the bregma and 2–3 mm from the midline. Mice were fixed in a stereotaxic frame, a mini-heating pad made in house was used to keep the body temperature at 37°C, and eye ointment (Rugby) was applied to prevent the eyes from drying out. A low dose of dexamethasone (0.1 ml at 1 mg/ml) was administered prior to surgery to minimize brain swelling. To minimize inflammatory phenomena that may occur after surgery, Ceftiofur sodium (100 μl, 0.1 g/ml in saline) was injected intraperitoneally on each of the following 3 days. Observation was usually performed 2 weeks after cranial window surgery.

For live imaging of the brain vasculature, head-fixed mice were placed on a customized microscope stage. To exclude the effect of anesthesia on cerebral blood flow, mice were imaged in a state of consciousness. Mice were accustomed to the platform for at least 30 min each day for 3 days before imaging. To observe blood flow, FITC-dextran dye (2,000 kDa, Sigma) at a dose of 100 μl (1% w/v, saline) was injected through the retro-orbital vein. Cerebral vasculature was observed by an upright two-photon laser scanning microscope (TPLSM, FVMPE-RS, Olympus) controlled by software (F31S-SW, Olympus) and a Mai Tai HP Ti:Sapphire laser (InSight X3TM, Spectra Physics) with a 25×, 1.05 NA water immersion objective lens (Olympus) at a resolution of 1024 × 1024 pixels, with galvanometric scanners. The optimal excitation wavelength was set as 990 nm for both FITC and DsRed. The emitted fluorescence from FITC and DsRed was detected with a non-descanned GaAsp detector (Olympus), and emission bandpass filters were BA575-645 (red) and BA495-540 (green). Z-stack images were collected at a 2-μm step size with a volume of 509 × 509 × 500 μm. For the measurement of red blood cell velocity in each capillary, X-T line scanning was conducted along a 15- to 50-μm range for 2,000 cycles.

### Image acquisition and quantification

Slide scanner imaging was performed on an Olympus VS 120 slide scanner (10×, 0.4 NA). Confocal imaging was performed on Leica SP8 laser scanning confocal microscopes with different objectives (20×, 0.75 NA; 40×, 1.3 NA; 63×, 1.4 NA). Cranial window images were captured through a stereo microscope (Olympus SZ61) with a smartphone camera. Images were processed using Aivia (Leica), LAS X (Leica), Olyvia (Olympus), and FIJI (NIH). Cell counting was performed using QuPath and Imaris. Labeling density was defined as the number of cells or blood vessels labeled over an area of 1 mm^2^ from brain section with 50μm thickness. To quantify the labeling efficiency of the AAV-miniBEND system, we calculated the percentage of EGFP^+^ cells over a ROI in the cortex and hippocampus (see Fig. 1) using the Hoechst channel as a reference for individual cell counts. Endothelial cells and non-endothelial cellswere counted, and we calculated the number of cells over a ROI corresponding to a rectangular or circular shape (area = 300,000 μm^2^). Briefly, to quantify conditional knockout efficiency, we calculated the density of Glut1^+^ and CD31^+^ (i.e., AAV-miniBEND–labeled cells) over a volume of 1 mm^3^. Cell counting was done using the “Draw counter” tool in the Leica microscope LAS X software Annotation panel. The acquired sample data consisted of multi-layer z-axis scanning confocal images of multiple ROIs and included three fluorescence channels: Hoechst (cell nuclei), Alexa 546 (GluT1), and Alexa 647 (CD31). Initially, the number of double-positive cells for CD31 and Hoechst was calculated to determine the total number of brain vascular endothelial cells within individual ROIs. Then, the number of double-positive cells for GluT1 and Hoechst was computed. The ratio of GluT1^−^ brain vascular endothelial cells to total brain vascular endothelial cells within each ROI was used to determine the knockout efficiency.

### Immunostaining

Mice were anesthetized by Avertin and perfused transcardially with PBS followed by 4% (w/v) paraformaldehyde. The brain was dissected out and fixed in cold paraformaldehyde (4 °C) for 2 h, washed thoroughly in PBS overnight, and dehydrated sequentially in 10%, 20%, and 30% sucrose in PBS. Fixed brains were sectioned at 40–60 μm with a cryostat (model CM3050S, Leica) or 60–70 μm with a Vibratome (VT1000S, Leica). Sections were stained as described (Ge et al., 2012). Briefly, sections were permeabilized with 0.25% Triton X-100 and then blocked with 5% BSA and 3% normal goat serum with 0.25% Triton X-100 for 2 h. The following primary antibodies were incubated with the brain sections for 24–48 h at 4 °C: PDGFRβ (1:300; rat, eBioscience, 14-1402-81), NG2 (1:300; rabbit, Millipore, AB5320), Ki67 (1:200; rabbit, Abcam, Ab15580), αSMA (1:200; rabbit, Abcam, Ab5694), GluT1 (1:100; rabbit, EMD Millipore, 07-1401), and CD31 (1:50; rat, BD Pharmingen, BD550274). After three washes in PBS together with Hoechst 33342 or DAPI (Life Technologies), sections were incubated with appropriate secondary antibodies conjugated with Alexa 488, Alexa 546, or Alexa 633/647 (1:500, Life Technologies) for 2 h at 22–25 °C. After three washes in PBS, sections were mounted with anti-fade mounting medium (Vector Laboratories) or a custom-made solution of 60% glycerol with Hoechst 33342). Whole-brain section images were taken with the tiling scan function on the confocal microscope (Leica SP8) with 10% overlap of adjacent tiles for automatic stitching. TUNEL staining (One Step TUNEL Apoptosis Assay Kit, Beyotime, C1089) was used to detect cell apoptosis with the immunostaining protocol.

### Vascular labeling with latex dye

Mice were anesthetized with 2.5% avertin (14 µl/g body weight) via intraperitoneal injection. The left ventricle was perfused with 0.1 M phosphate buffer until the liver appeared pale, followed by 4 ml of latex dye solution. The latex dye, which cannot cross capillary beds, was retained within arterial branches in the brain. Brains were then fixed in 10% formalin for 12 hours, dehydrated through a methanol gradient, and cleared for transparency. Latex dye was observed lodged in arterial and venous branches after clearing.

### Tissue clearing and lightsheet imaging

Mouse brains were cleared using the polyethylene glycol (PEG)-associated solvent system (PEGASOS) method^83^. Briefly, retro-orbital injection of 100 μl Lectin–DyLight 488 (Vector Laboratories, DL-1174-1) was performed 5 min before fixation. Brains were washed with PBS (three times, 10 min each time) after fixation with 4% paraformaldehyde solution for 4 h. Brains were then treated with Quadrol decolorization solution for 2 days at 37 °C, at which point they were immersed in gradient delipidation solution in a 37 °C shaker for 1–2 days, followed by dehydration solution treatment for 1–2 days and BB-PEG clearing medium treatment for at least 1 day until reaching transparency. Brains were then preserved in the clearing medium at 4 °C for ∼3 days before imaging with a lightsheet microscope (LaVision BioTec, UltraMicroscope II). Images were stitched using Imaris Stitcher, and an animated video of each brain was made with Imaris (9.0) software.

### Whole-brain vasculature imaging

All samples were processed using the PEGASOS tissue clearing kit (Leads Bio-Tech, Catalogue number: PSK100N). Brains from miniBEND-infected *Braf ^V600E^fl/fl* mice were perfused with PBS and fixed in 4% paraformaldehyde (PFA) at 4°C overnight. Samples were decolorized in Solution 2 (25% Quadrol in H[O) at 37°C for two days, followed by serial delipidation at 37°C using Solutions 3, 4, and 5 (30%, 50%, and 70% tert-butanol in H[O) each for one day. The samples were then dehydrated in Solution 6 (70% tert-butanol and 30% Quadrol) at 37°C for two days, followed by immersion in Solution 7 (BB-PEG clearing medium) at 37°C until fully transparent, typically within one day. A minimum of 30 ml of solution was used for each step per sample.

Imaging was conducted using a 10X objective (N.A. = 0.6, WD = 8 mm) on a LiTone XL Light-sheet Microscope (Light Innovation Technology Limited). Images were stitched with LiTScan 3.3.0 and processed with Imaris 9.0.

### Flow cytometry of brain cells

Mice aged P30–45 were anesthetized with isoflurane and perfused with precooled PBS (HyClone) via the heart. Brains were isolated and immediately immersed in ice-cold HBSS (5 mM KCl, 5 mM NaOH, 5 mM NaH[PO[, 0.5 mM MgCl[, 20 mM sodium pyruvate, 5.5 mM glucose, 200 mM sorbitol, pH 7.3– 7.4, filtered with a 0.22 μm filter). Each brain hemisphere was diced into small pieces and incubated in HBSS with 2 mg/ml papain (Sigma) and 5 μg/ml DNase I (Sigma) at 37°C for 30 minutes. A fire-polished Pasteur pipette (BrainBits) was used to dissociate the tissue into a single-cell suspension.

The cell suspension was then added to an Optiprep gradient^77^ and centrifuged at 1000 g for 15 minutes. The top debris layer was removed, and specific layers were collected. Cells from each layer were washed with HBSS and pelleted at 700 g for 10 minutes. After discarding the supernatant, cells were incubated in 1 ml of HBSS containing Alexa Fluor® 647 Rat Anti-Mouse CD144 (BD) at room temperature for 40 minutes in the dark. The cells were then centrifuged at 300 g for 5 minutes, the supernatant was discarded, and the cells were resuspended in 1 ml HBSS for sorting.

### Single-cell RNA-seq data analysis of harvested endothelial cells

Single-cell sequencing data were obtained and pre-processed from AAV-miniBEND and Braf-CA transgenic mice 11 days post-injection, based on a focal bAVM disease model. The single-cell transcriptome expression matrix data were merged using Anndata, following instructions in the SCANPY tutorial^84^. During the quality control step, cells with a minimum gene count of 300 and a minimum cell count of 50 with respect to the expression of the gene of interest were retained. Cells with more than 15% mitochondrial gene counts were filtered out. Subsequently, the count matrix was log-transformed. After these preprocessing steps, a total of 28,954 cells were obtained for downstream analysis. For annotation and collection of endothelial cells, manual evaluation of classic marker genes was used (PECAM1 and CLDN5). The bAVM group comprised a total of 2,624 endothelial cells, and the Ctrl group comprised a total of 3,022 endothelial cells, which were harvested for subsequent analysis. The MELD algorithm^85^ was used to estimate the differences in each cell under bAVM and Ctrl conditions. Harmony^86^, with default parameters, was used for batch effect removal, and Louvain clustering was performed with a resolution parameter set to 1.0. A Wilcoxon test with tie correction was used to obtain ranked marker genes. Gene set enrichment analysis was performed using GSEAPY. Gene Set Enrichment Analysis was performed using permutation testing and Two-sided P values were calculated, and FDR correction was applied.

### Statistics and reproducibility

Animals were randomly assigned to groups during the experiments. QuPath, Imaris, and NIH ImageJ software were used to quantify the data from confocal microscopy images after immunostaining. All data were analyzed with Prism 8 (GraphPad) software and are presented as the mean ± s.e.m. Statistical significance was evaluated using Prism 8. *p*-Values were calculated using an unpaired two-sided Student’s t-test or Welch’s t-test. Two-tailed *p* < 0.05 was considered statistically significant; values are denoted as follows: \**p* < 0.05; \*\**p* < 0.01; \*\*\**p* < 0.001; **** *p* < 0.0001. All fluorescence intensity, cell counting and immunostaining experiments were independently performed ≥2 times with similar results.

## Data availability

The authors confirm that the data supporting the findings of this study are available within the article and its supplementary material. Derived data supporting the findings of this study are available from the corresponding authors on request.

## Acknowledgements

We thank M.S. Xinwei Gao, Mingyue Jia, and Dr. Qingchun Guo from the CIBR Imaging facility for providing microscopic imaging and data analysis service and from the CIBR vector core for providing AAV packaging services; Shufang Huang and Dr. Wenlong Li from the CIBR Laboratory Animal Resource Center for their support; and our colleagues for critical reading of the manuscript. We thank members of the Ge and Sun Laboratories and other colleagues from CIBR for their feedback on this work. This project is financially supported by funds CAMS Innovation Fund for Medical Sciences (CIFMS, 2024-I2M-ZD-012) to W-P.G., the STI2030-Major Projects (2022ZD0204700) to W.S. and W-P.G., and funds from CIBR, Changping Laboratory, the Feng Foundation of Biomedical Research, and National Natural Science Foundation of China (No. 32170964) to W-P.G.

## Author contributions

J.L.L. and W.-P.G. conceived the project. W.-P.G. and W.S. supervised the whole project and helped to design the experiments. J.L.L. and Z.Y.B. performed the experiments in Fig. 1, Fig. 2, and Fig. 4 (with the exception of staining) and made all of the constructs used in this study. J.L.L. and X.J.C. performed disease modeling for CCM and bAVM with input from W.-P.G. J.L.L., X.J.C. performed the staining and imaging in Fig. 4 and most Supplementary Figures. Z.Y.B. and T.Y.M. assisted in staining and immunohistochemistry data analysis in Fig. 4. J.L.L. performed *in vivo* imaging of bAVM mice with the help of B.S.Q. and X.J.C. X.J.C., J.Y.W. and S.L. performed tissue clearing and lightsheet imaging of bAVM mouse brains with technical input from H.Z… Z.Y.F. provided help with mouse breeding and maintenance. F.Z.L. provided help with FACS sorting. J.H. generated transgenic rat, J.L.L. and Y.P.L. performed rat experiments, F.Z. provided help with AAV packaging. T.Y.M., J.L.L. and D.S.A. performed snRNA-seq data analysis. Z.J.W. and T.T.Z provided help in MRI data acquisition and analysis, respectively. J.L.L. and X.J.C. performed all MRI experiments with the help of Z.J.W.. W.-P.G., W.S., F.Z., Y.L.W. provided reagents and necessary instruments. J.L.L., Z.Y.B. and W.-P.G. wrote the manuscript. All authors reviewed and edited the manuscript.

