## supplemental Files for "rAAV-miniBEND: A targeted vector for brain endothelial cell gene delivery and cerebrovascular malformation modeling"

#### Supplementary Figure 1

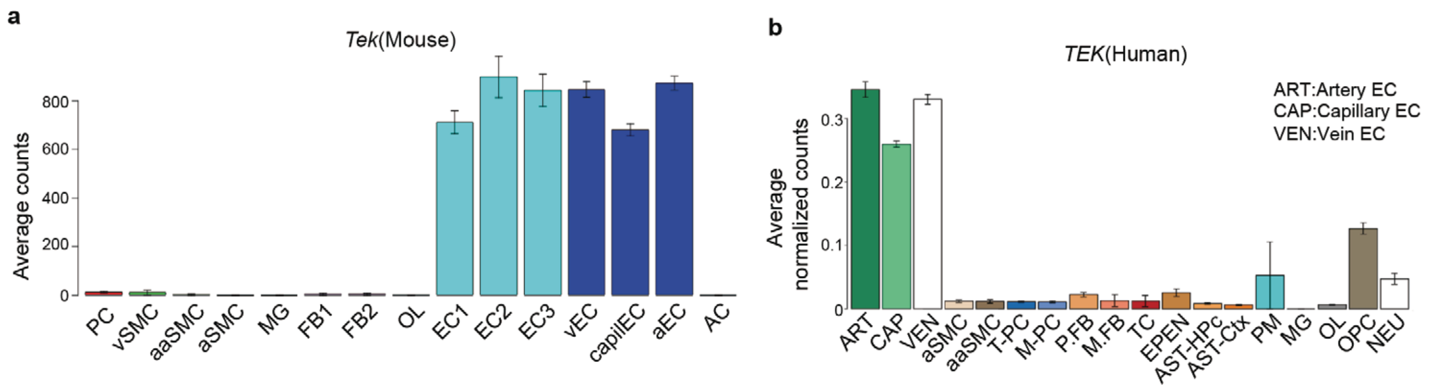

**Supplementary Figure 2**

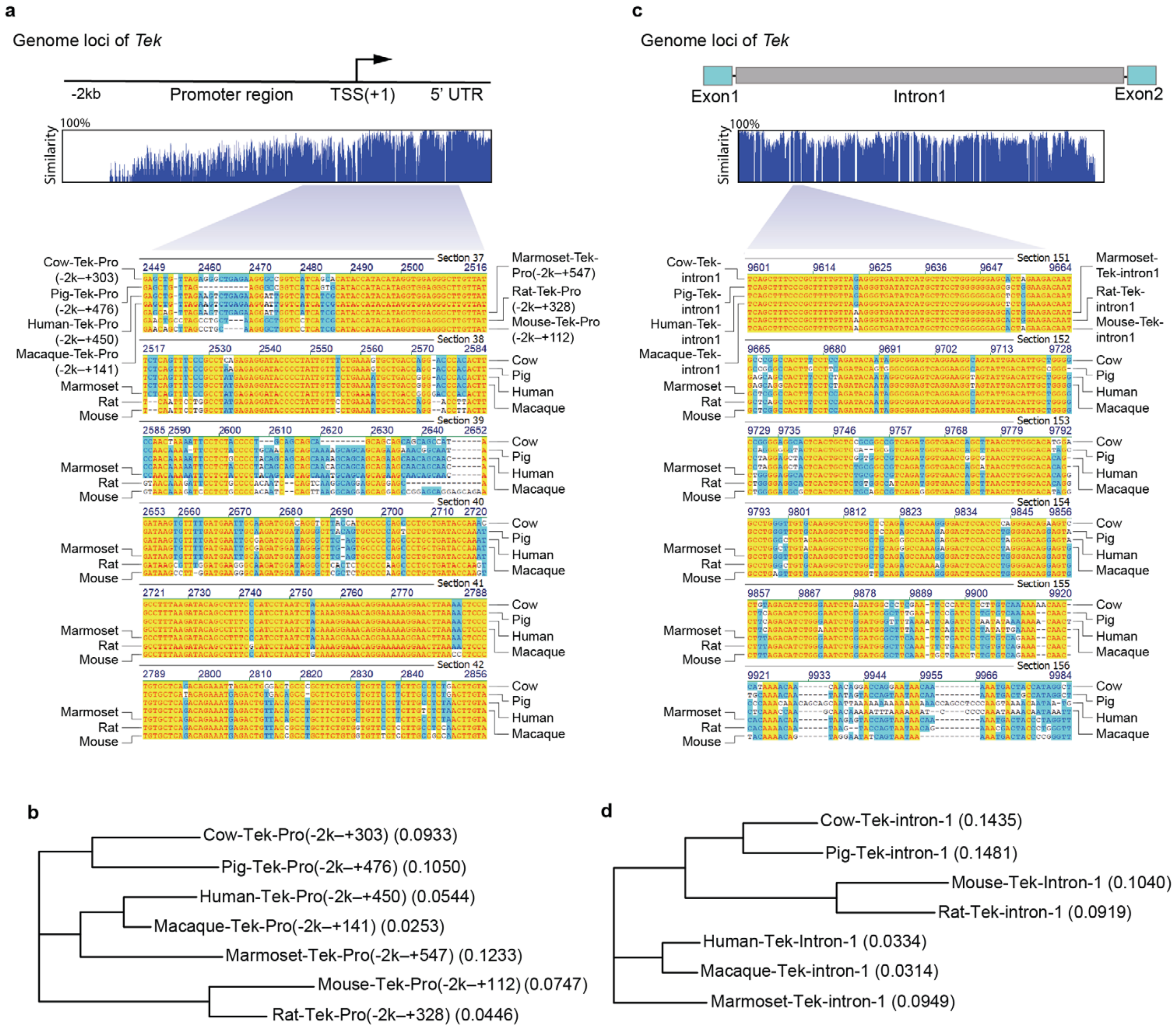

**Supplementary Figure 3**

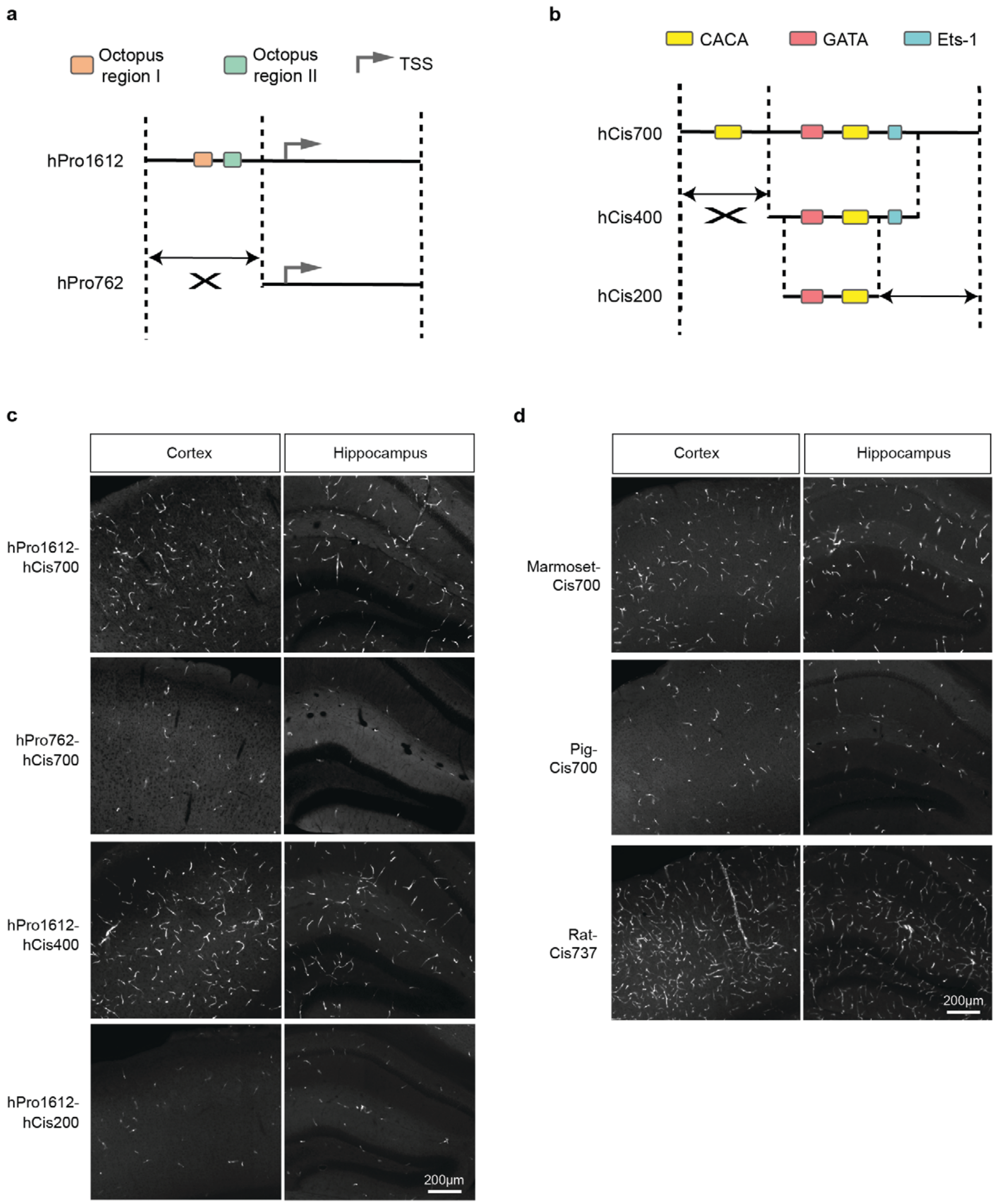

**Supplementary Figure 4**

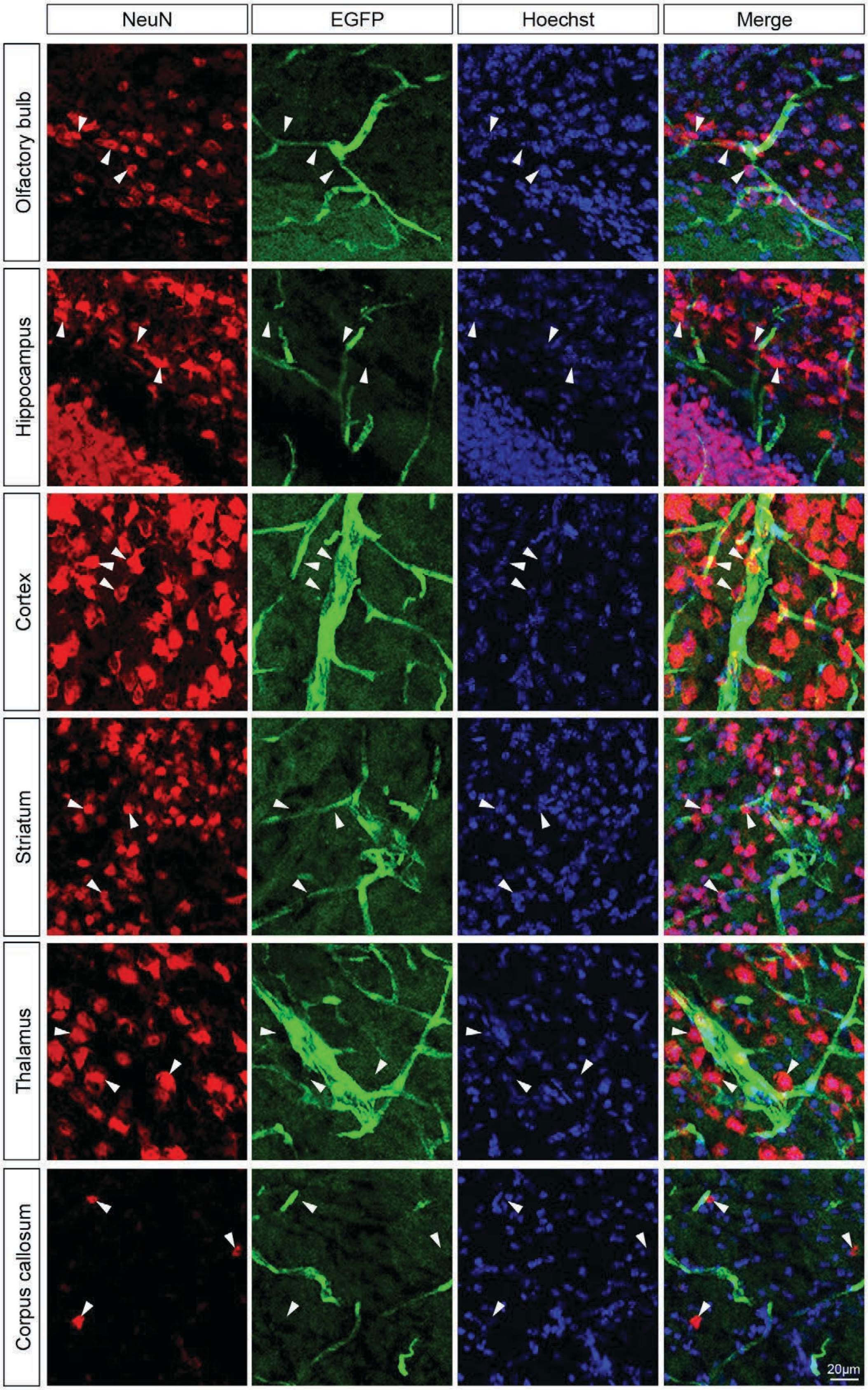

**Supplementary Figure 5**

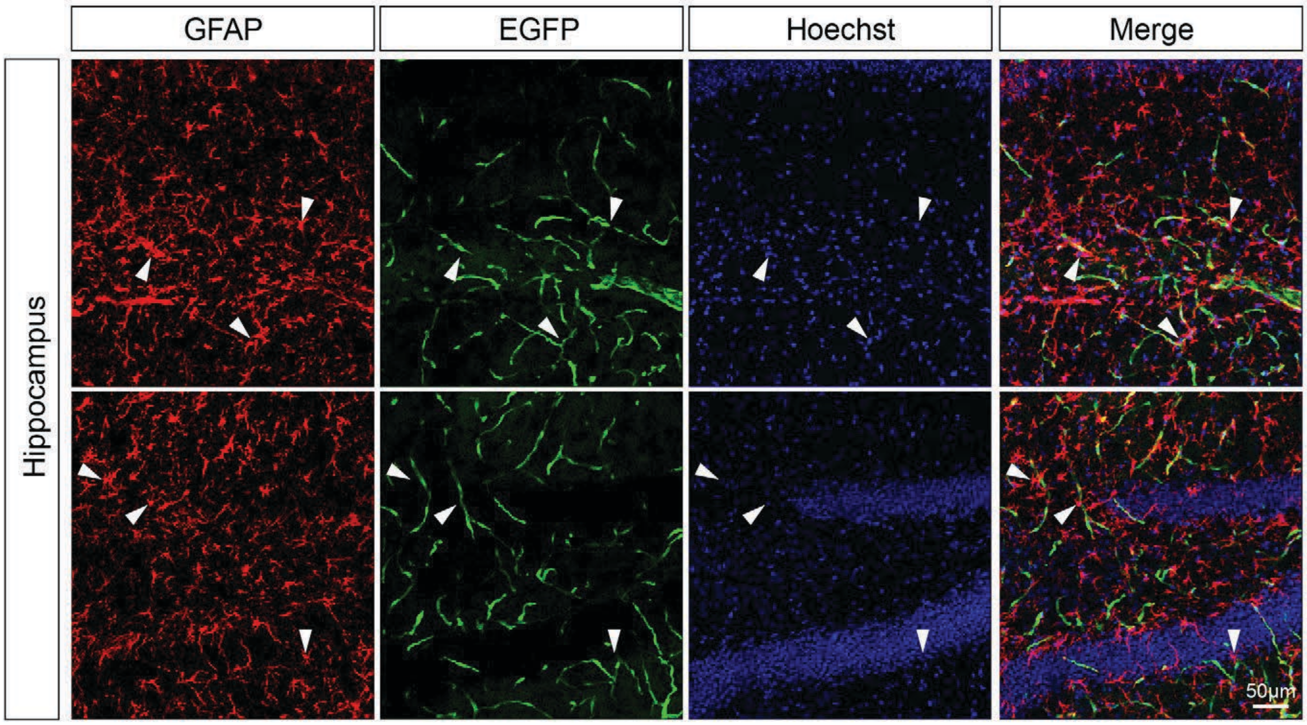

**Supplementary Figure 6**

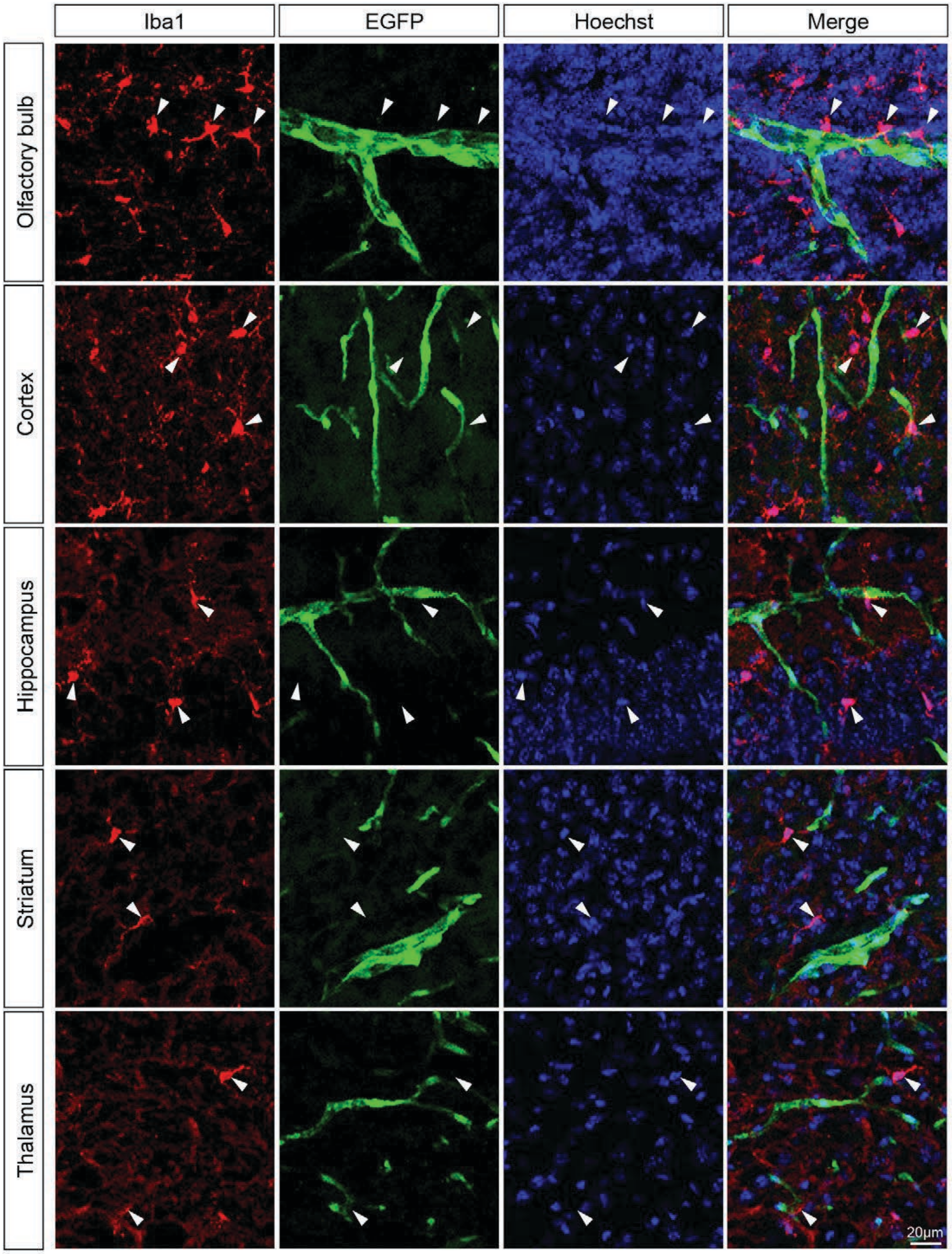

**Supplementary Figure 7**

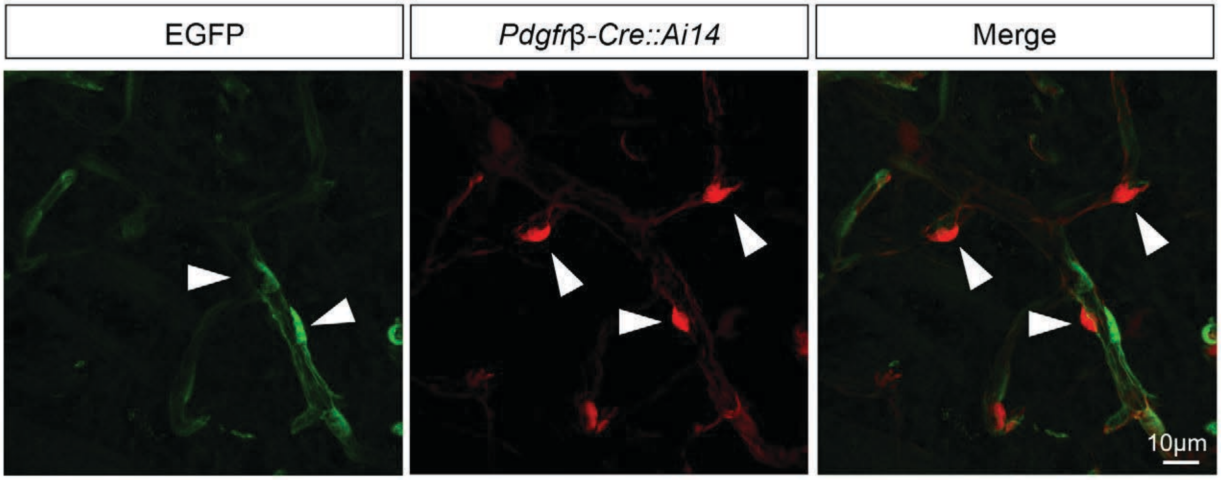

#### Supplementary Figure 8

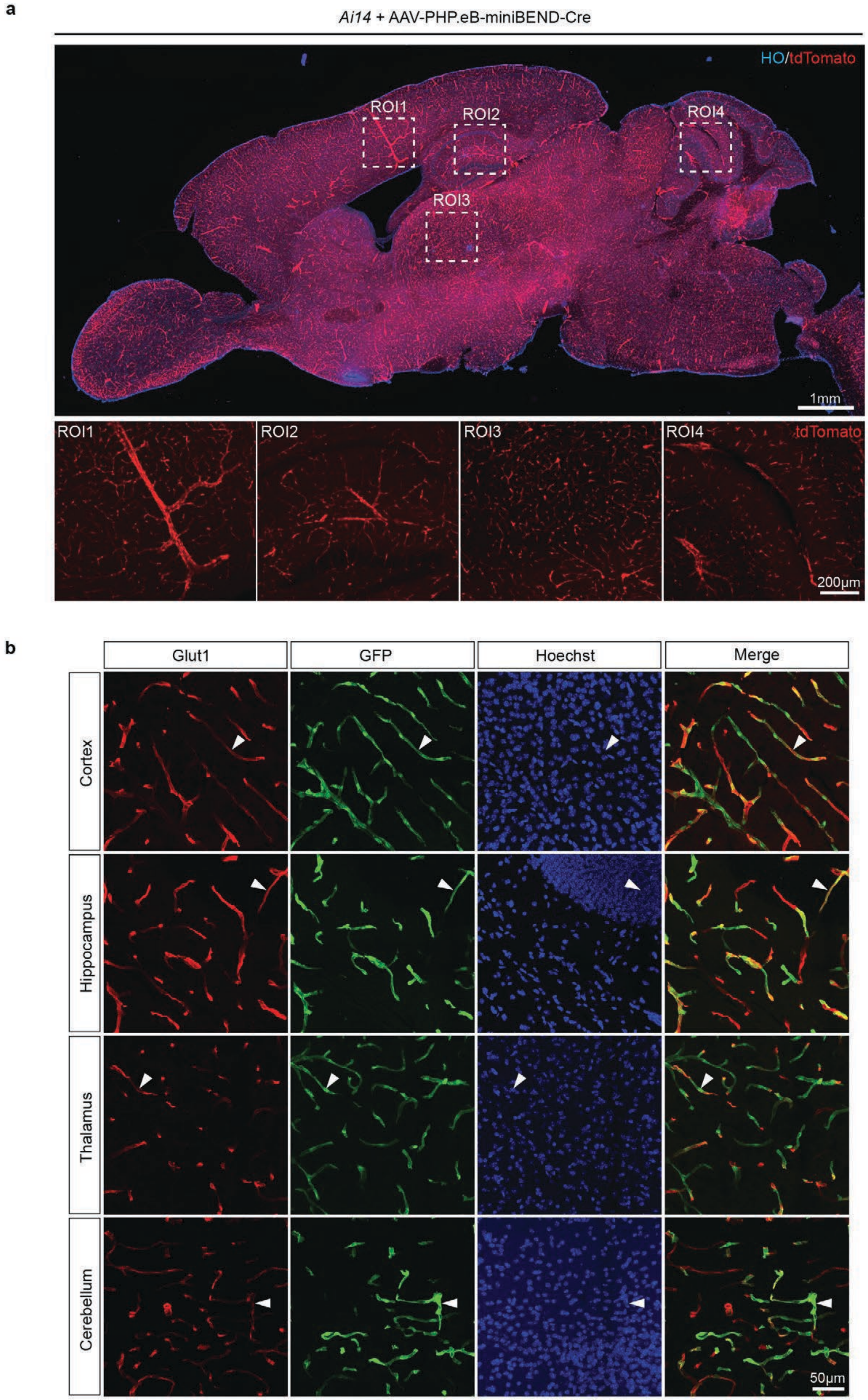

**Supplementary Figure 9**

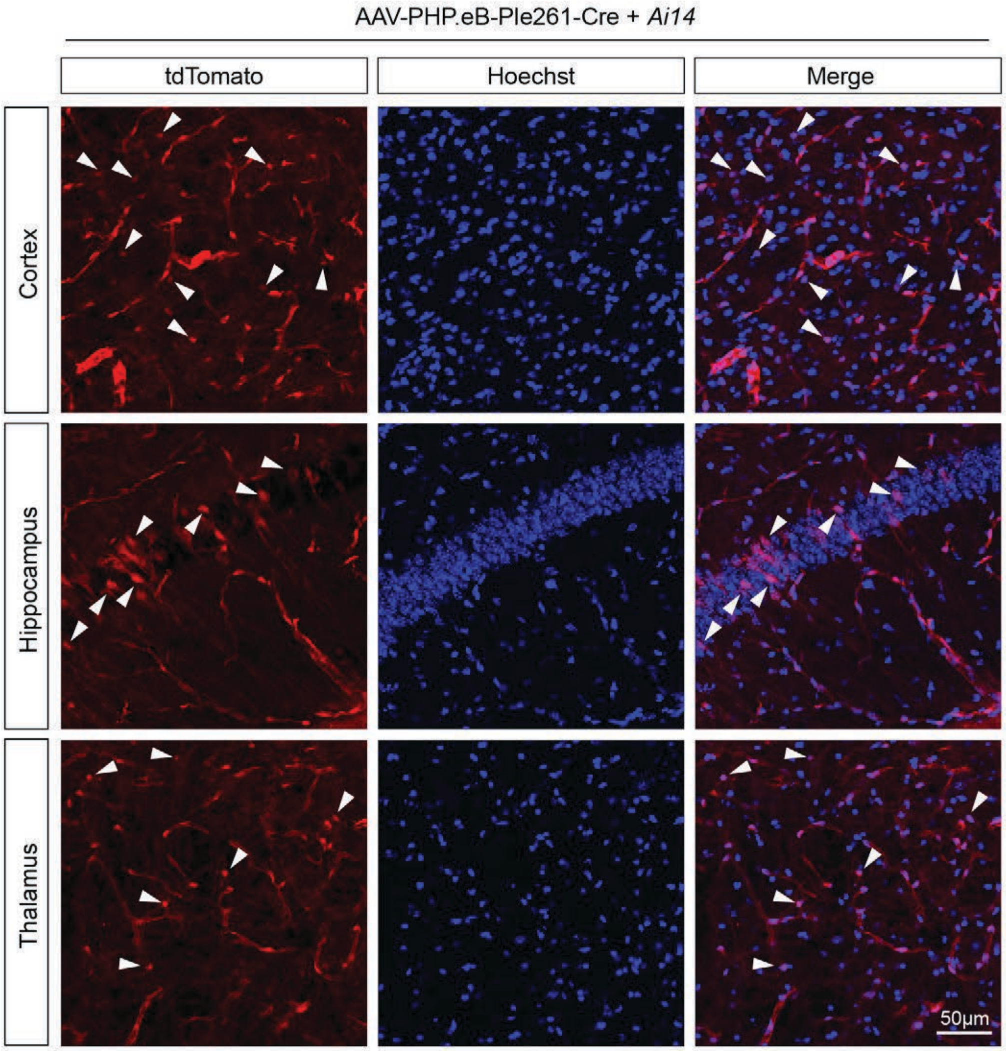

**Supplementary Figure 10**

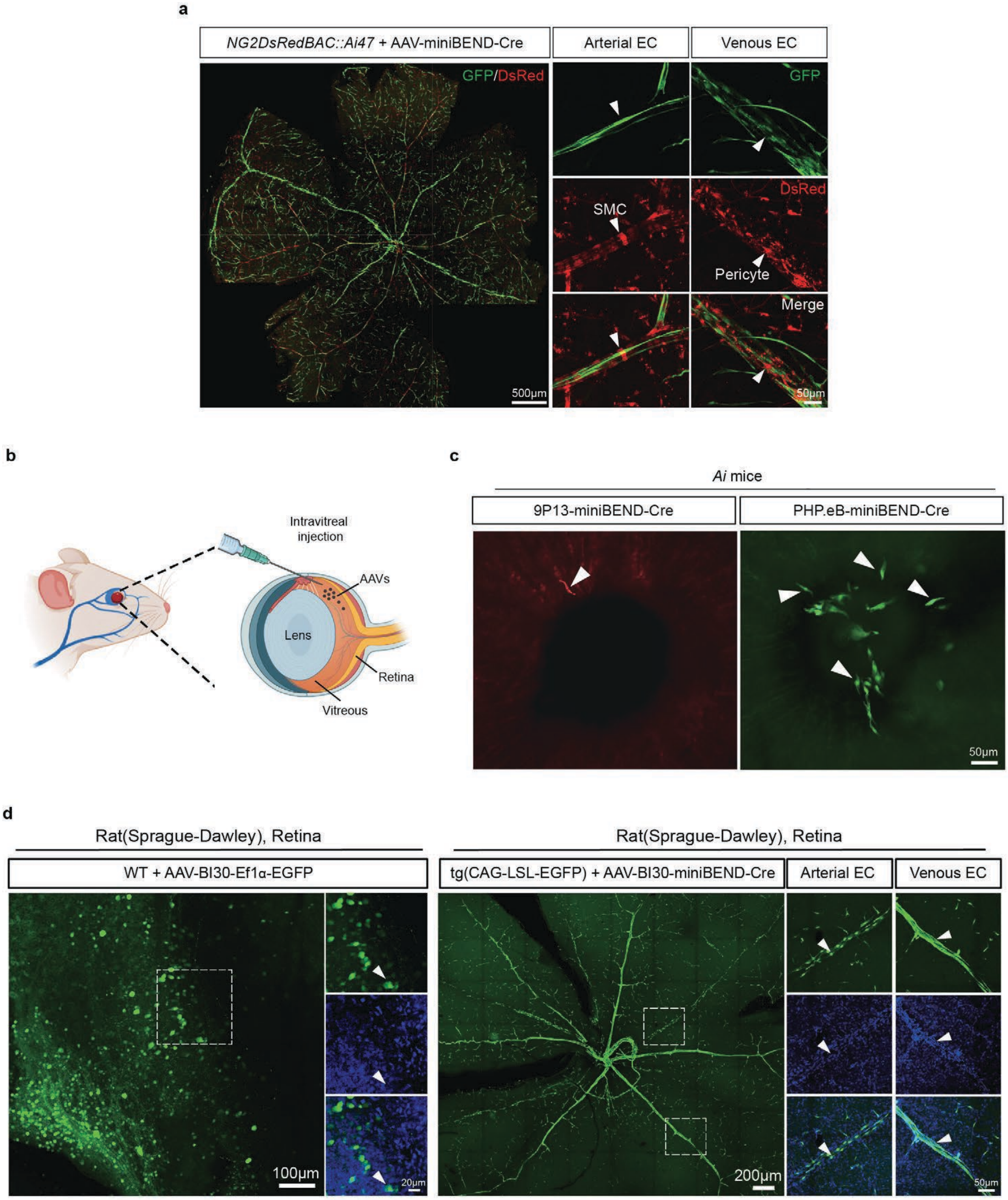

**Supplementary Figure 11**

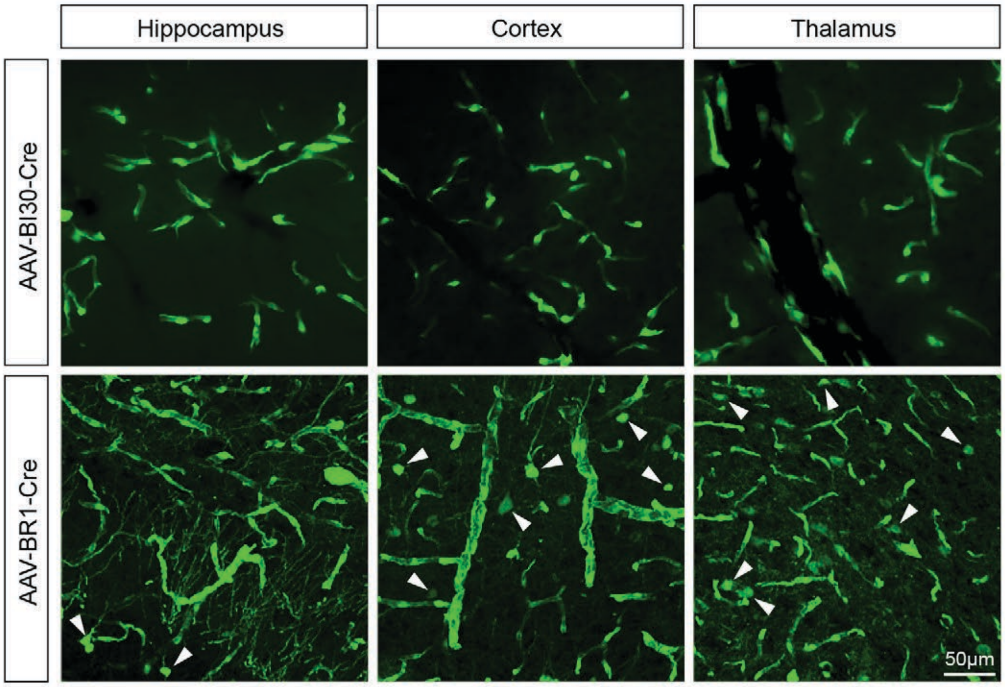

#### Supplementary Figure 12

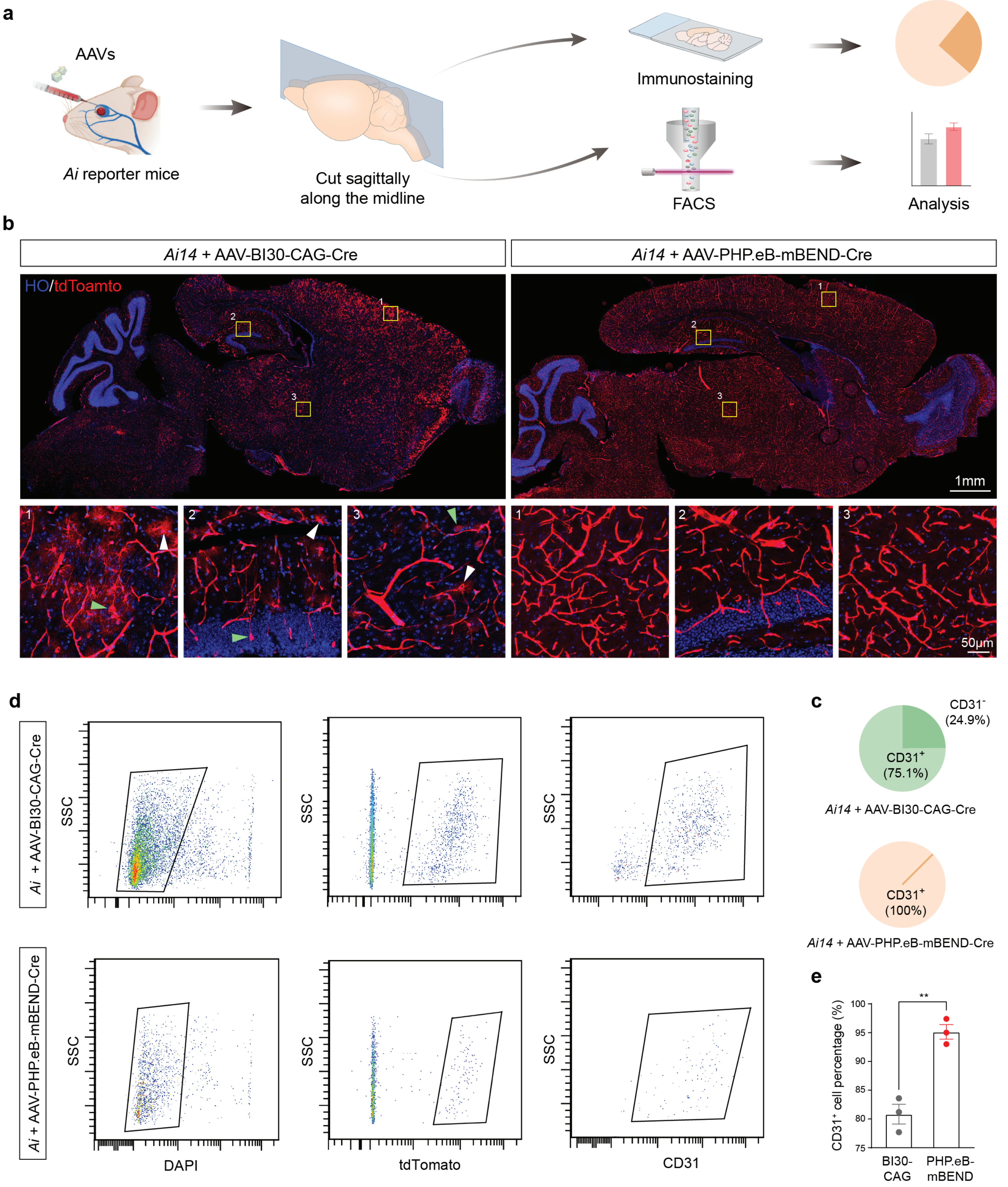

**Supplementary Figure 13**

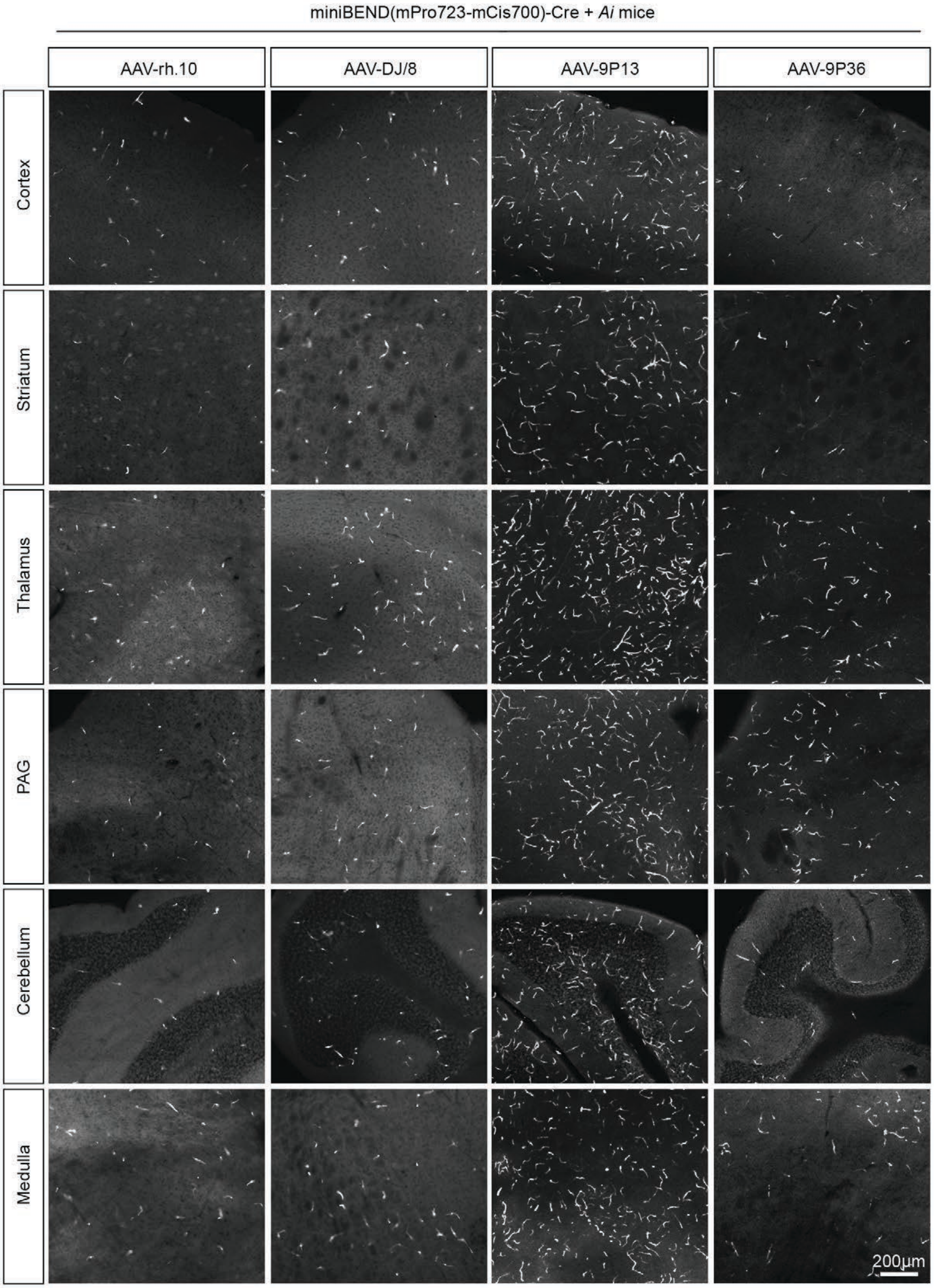

**Supplementary Figure 14**

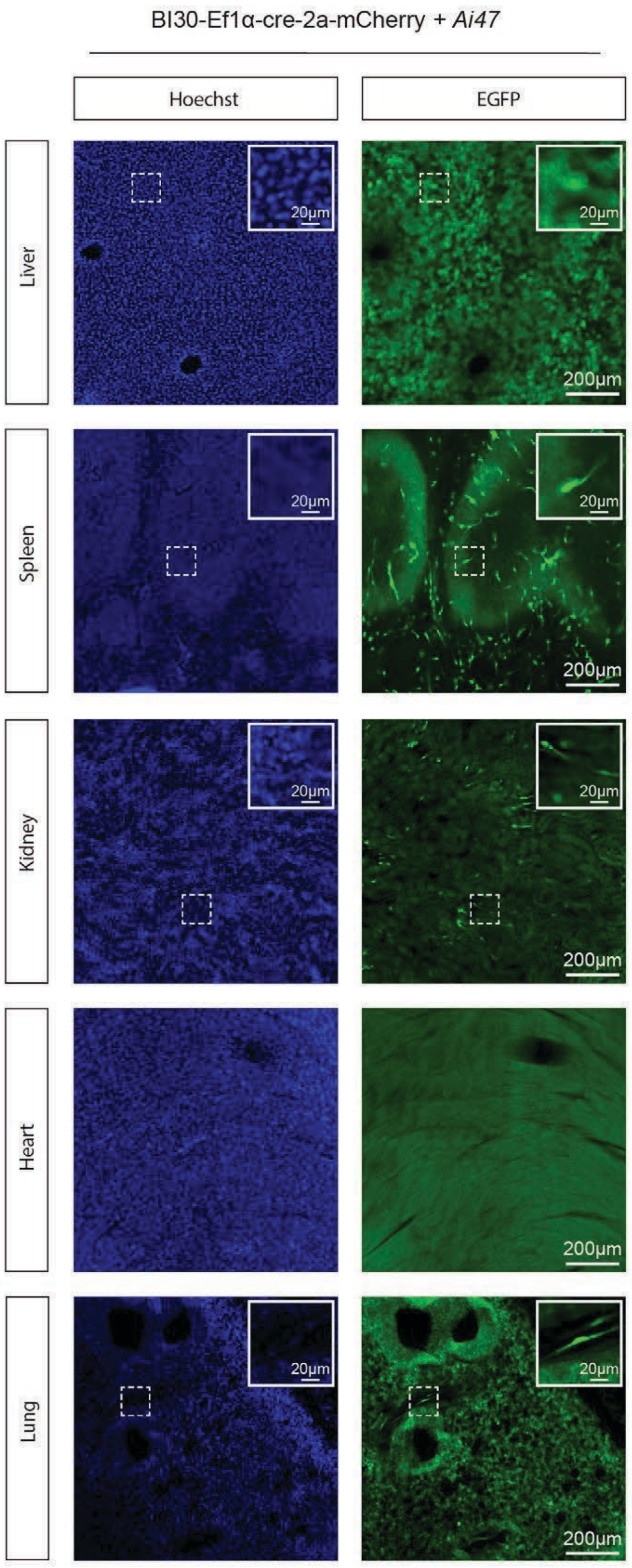

**Supplementary Figure 15**

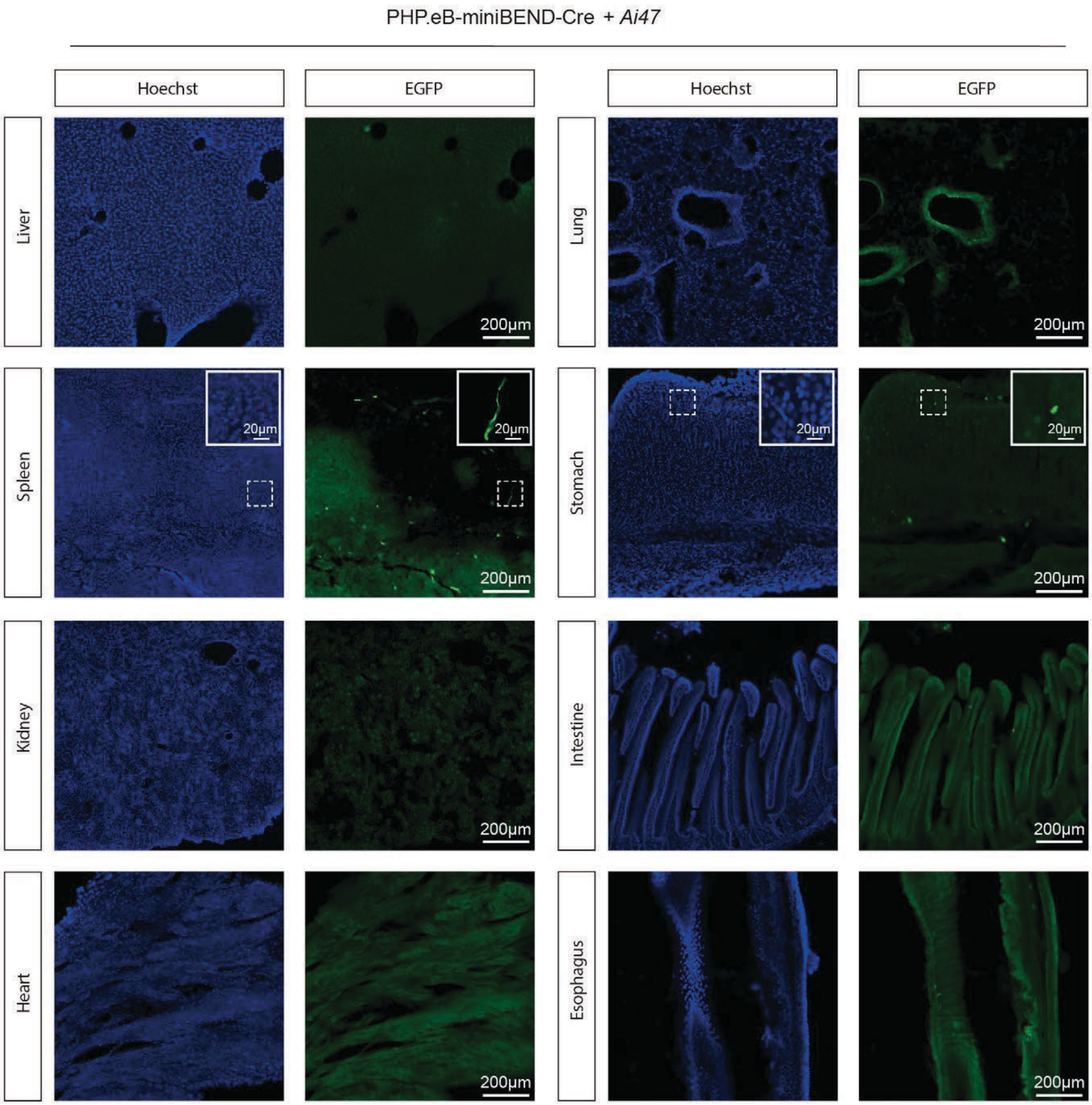

**Supplementary Figure 16**

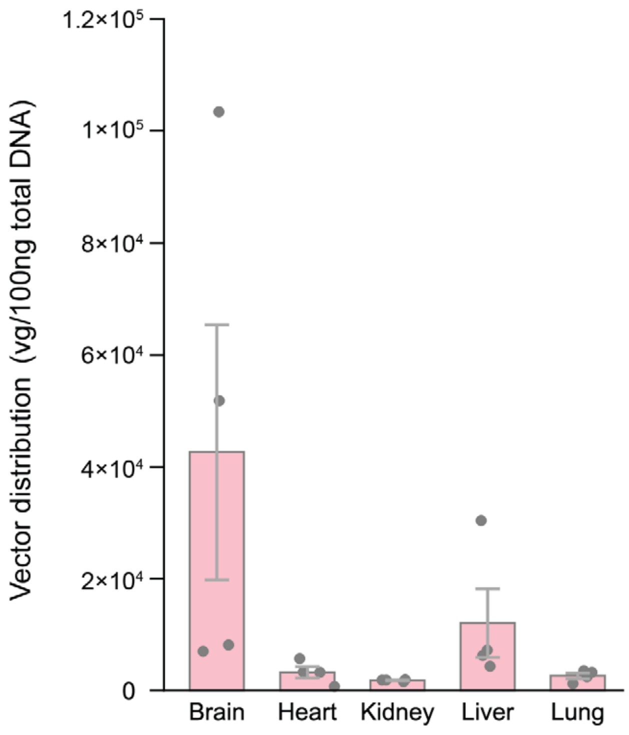

#### Supplementary Figure 17

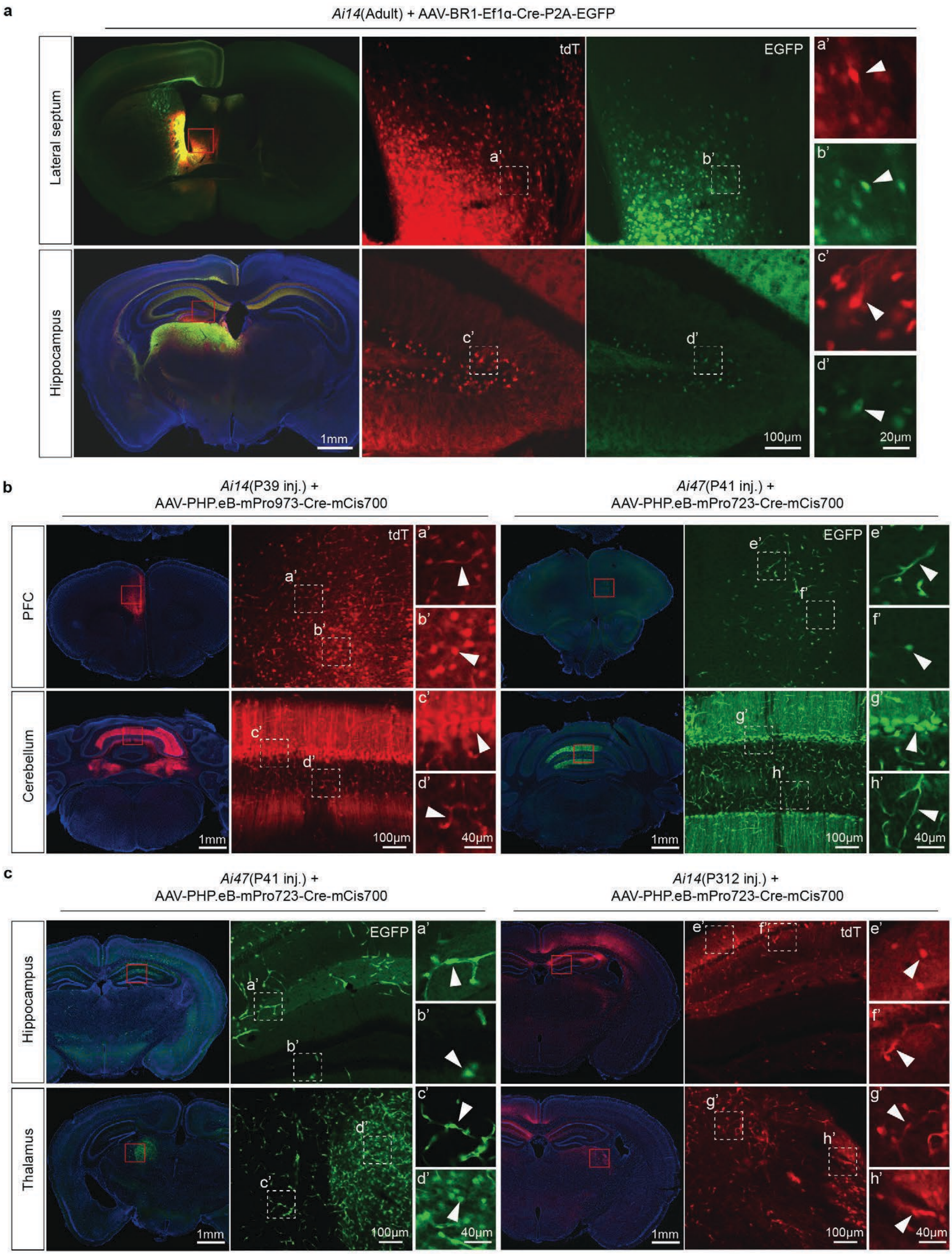

**Supplementary Figure 18**

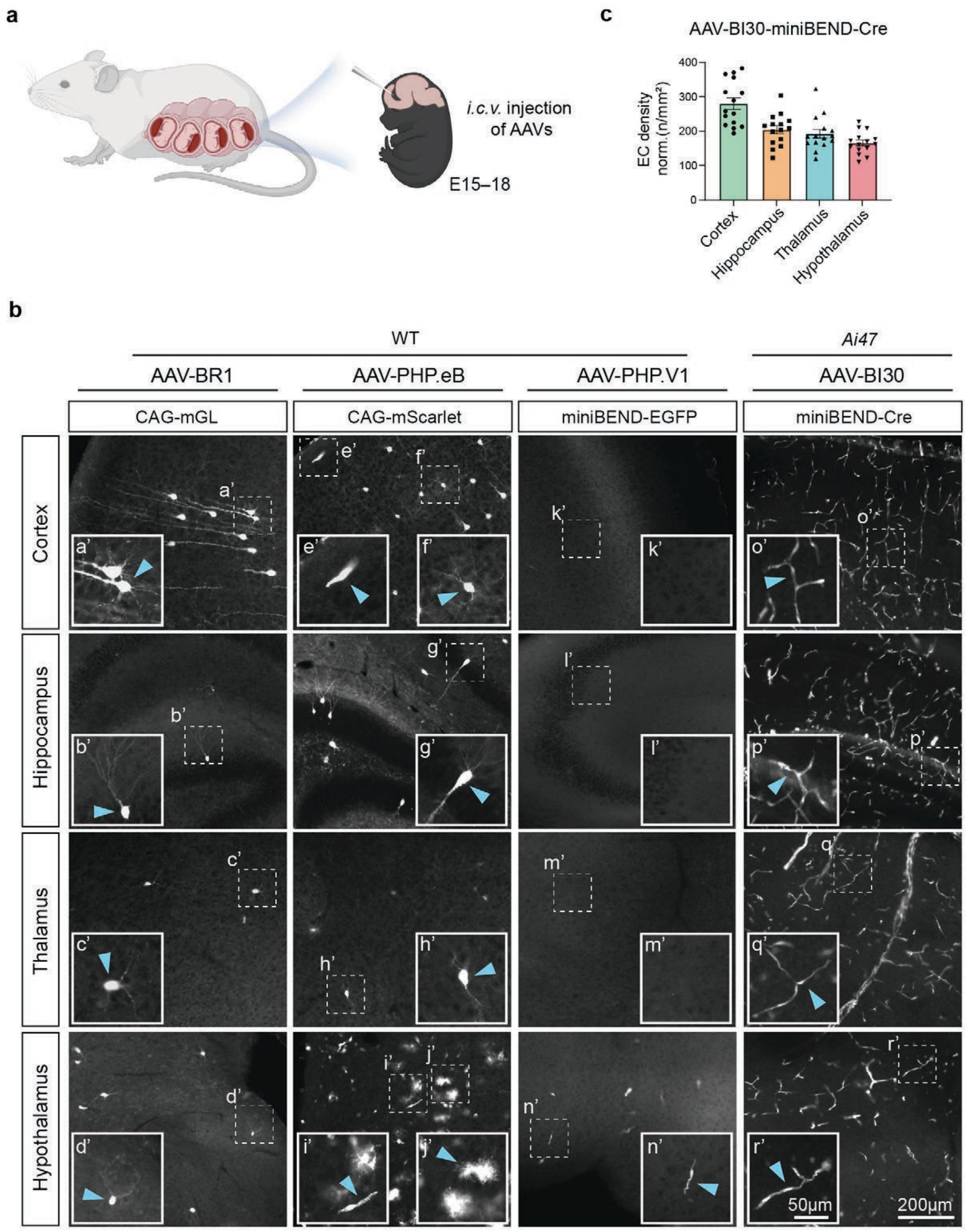

**Supplementary Figure 19**

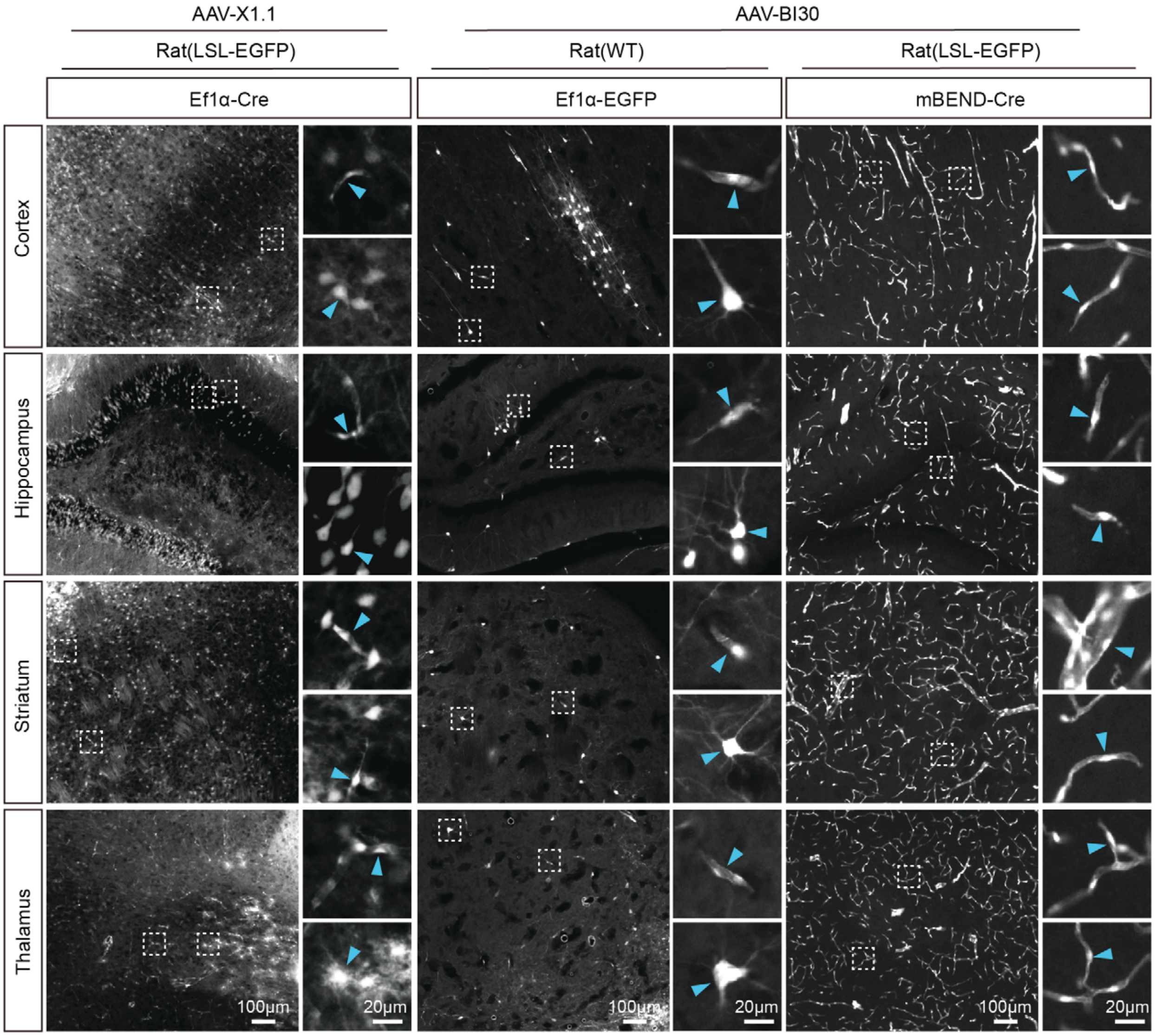

**Supplementary Figure 20**

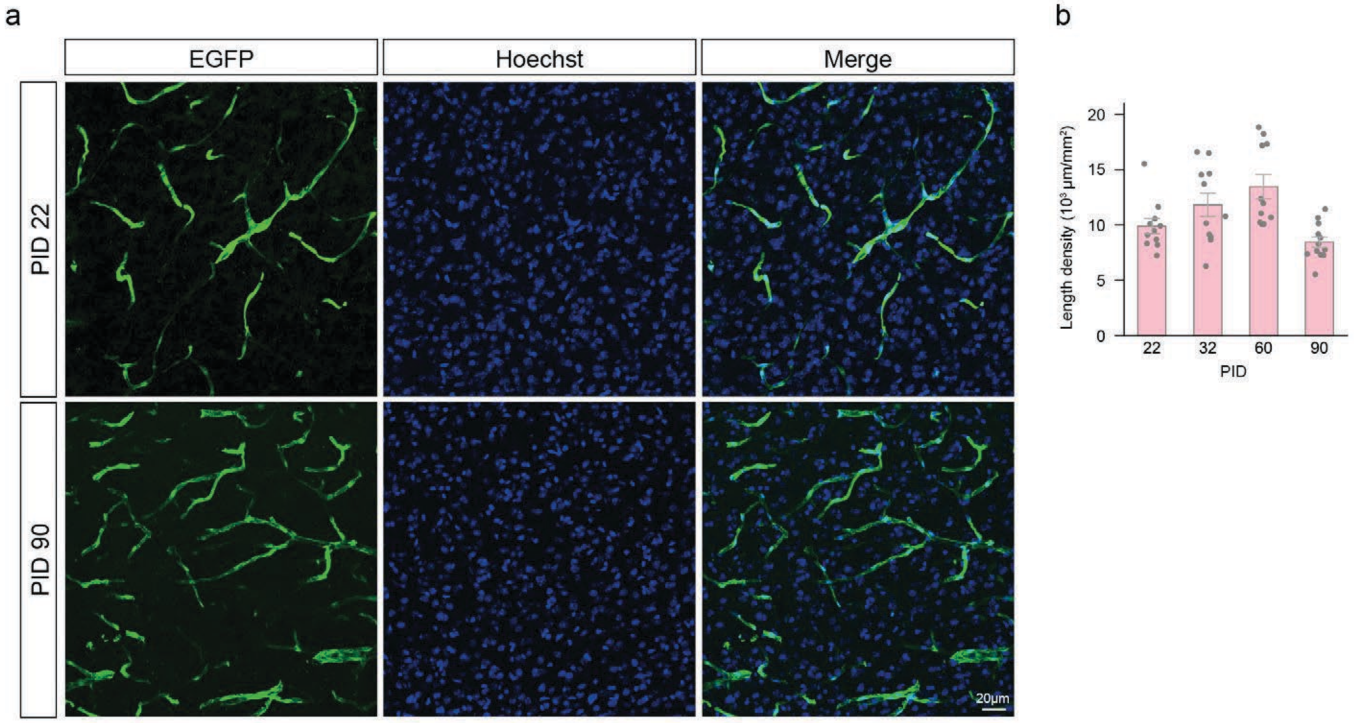

**Supplementary Figure 21**

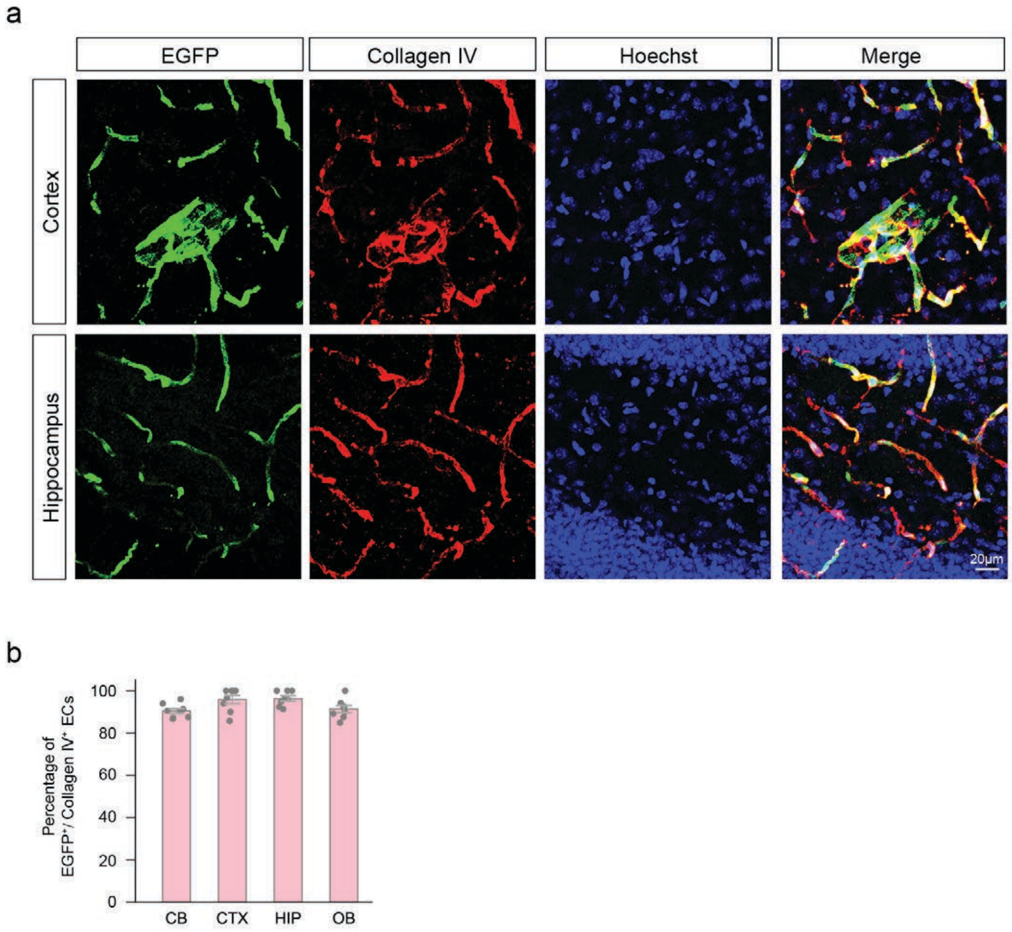

#### Supplementary Figure 22

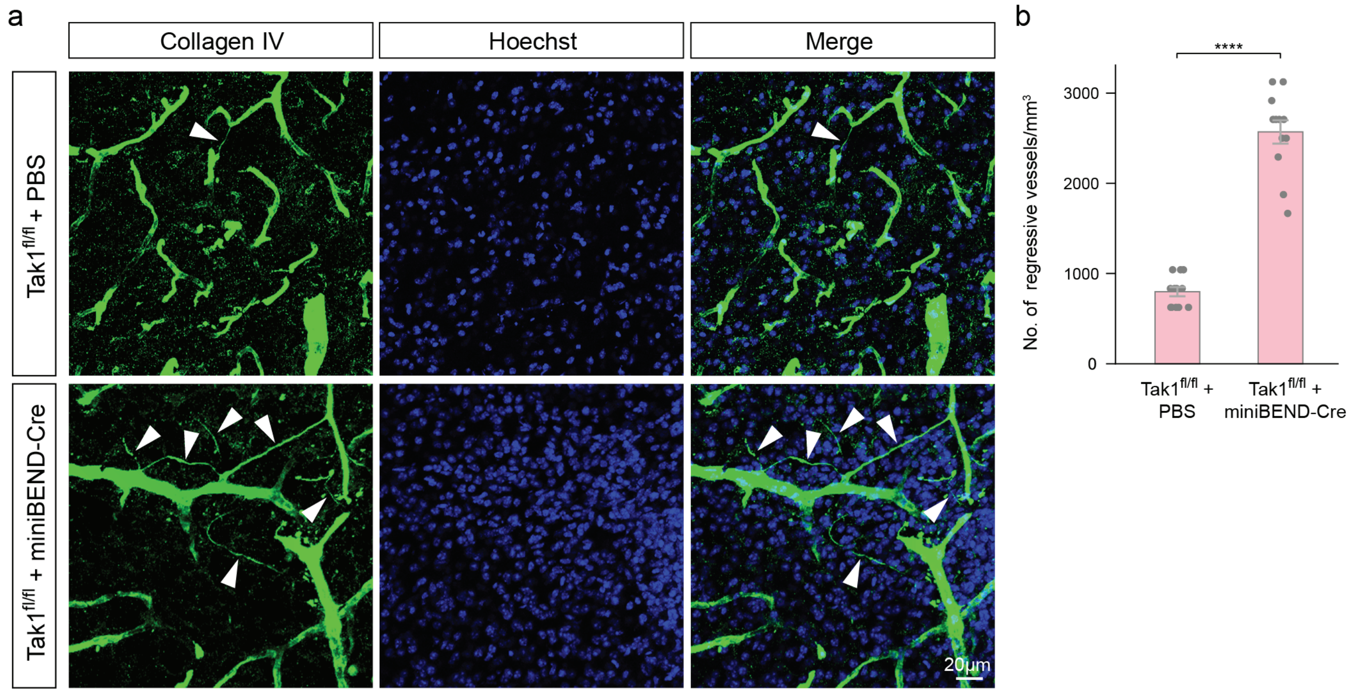

**Supplementary Figure 23**

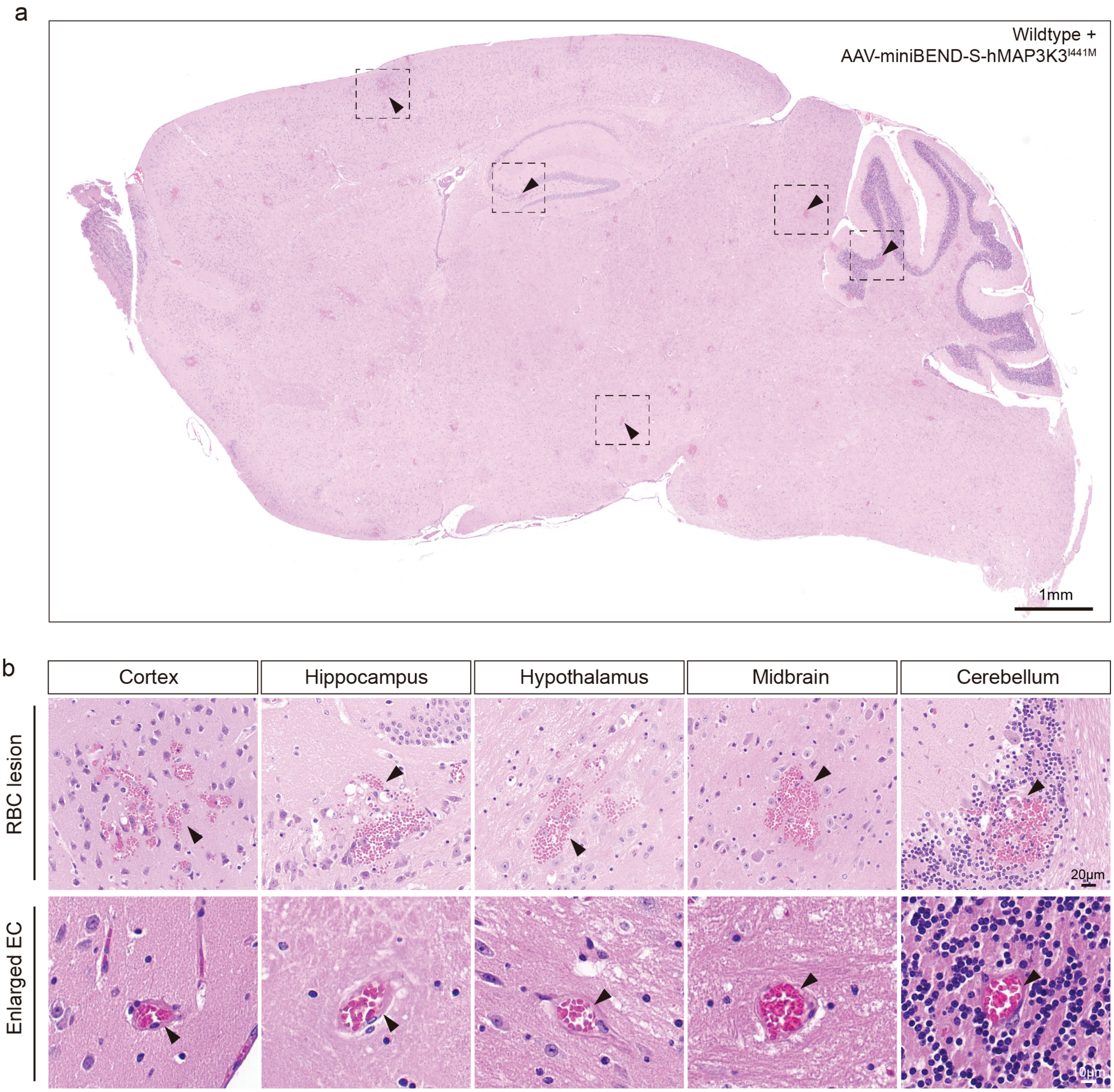

**Supplementary Figure 24**

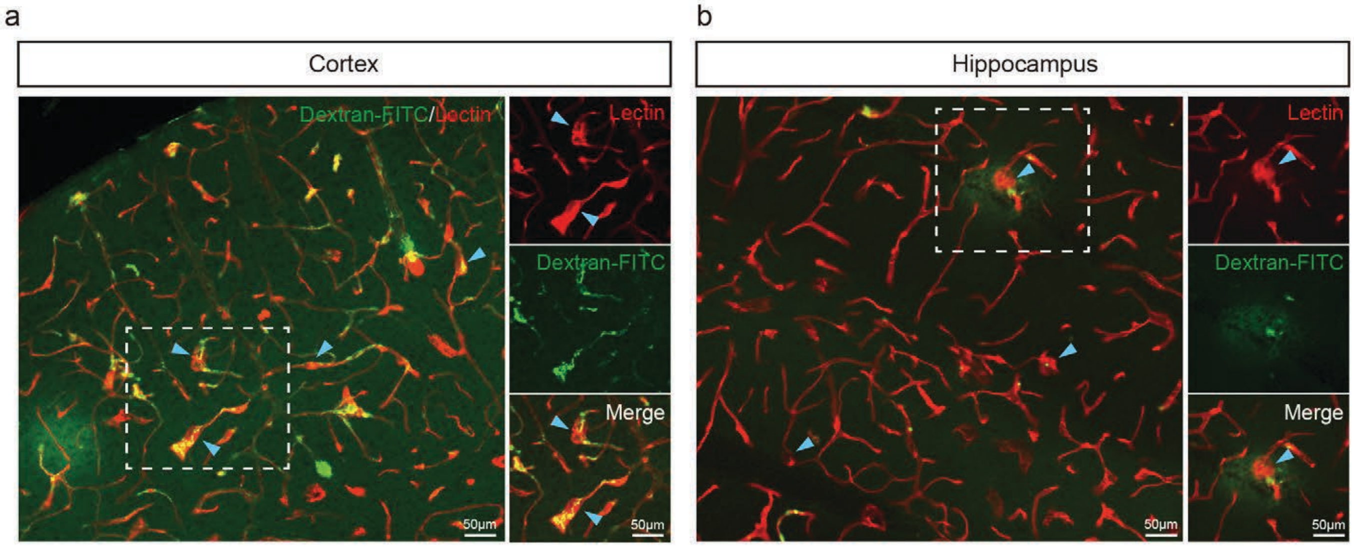

**Supplementary Figure 25**

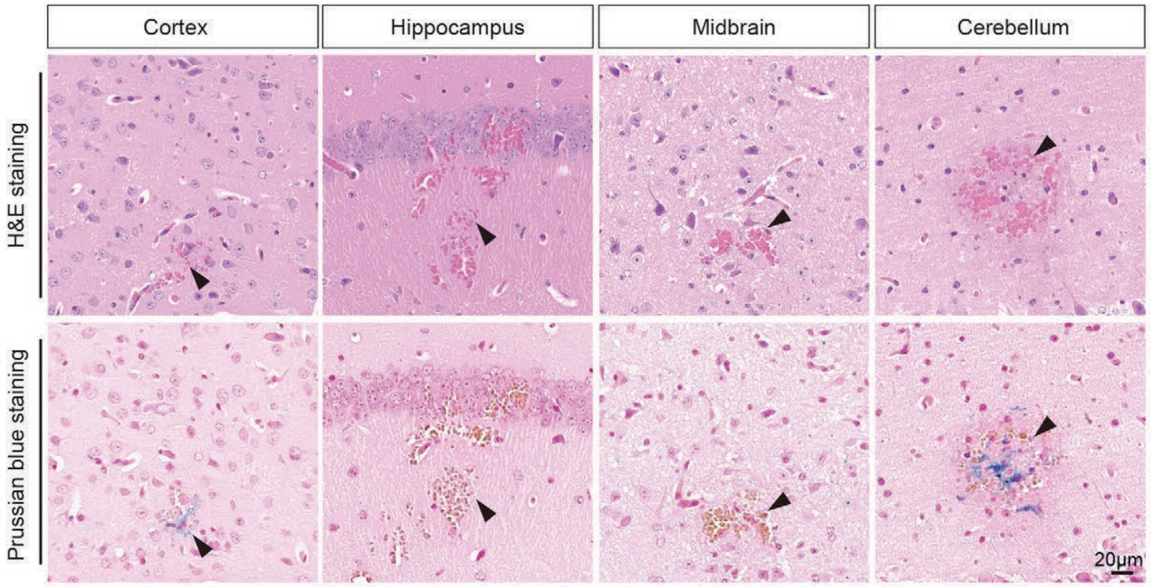

**Supplementary Figure 26**

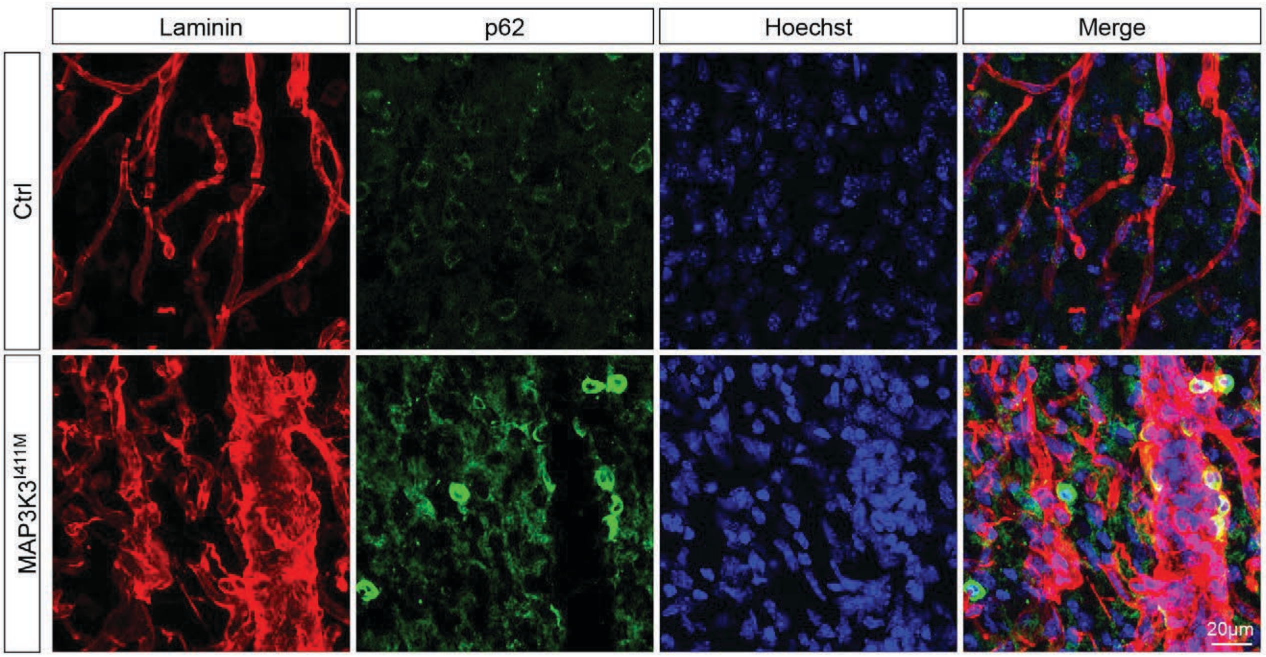

#### Supplementary Figure 27

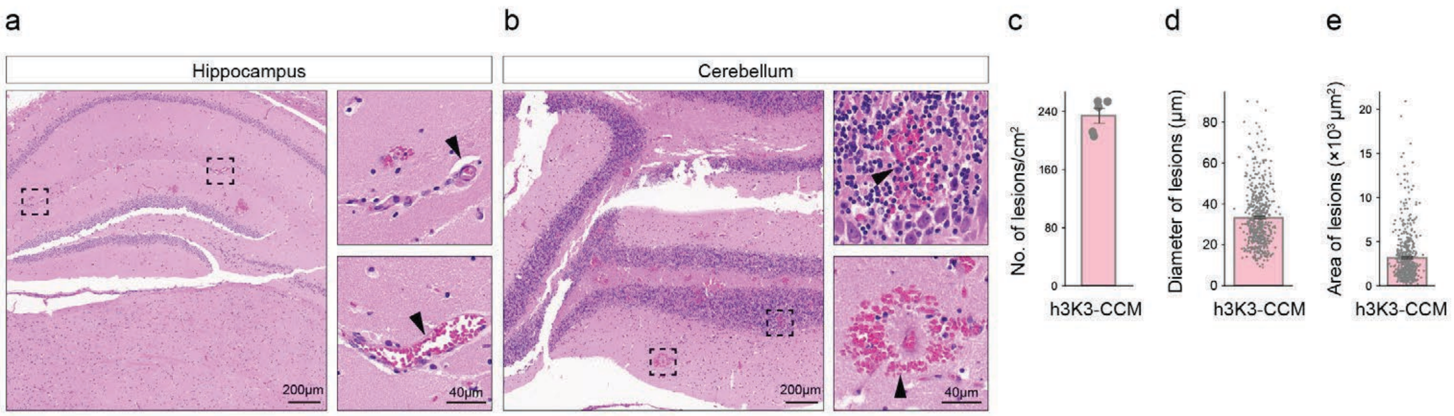

**Supplementary Figure 28**

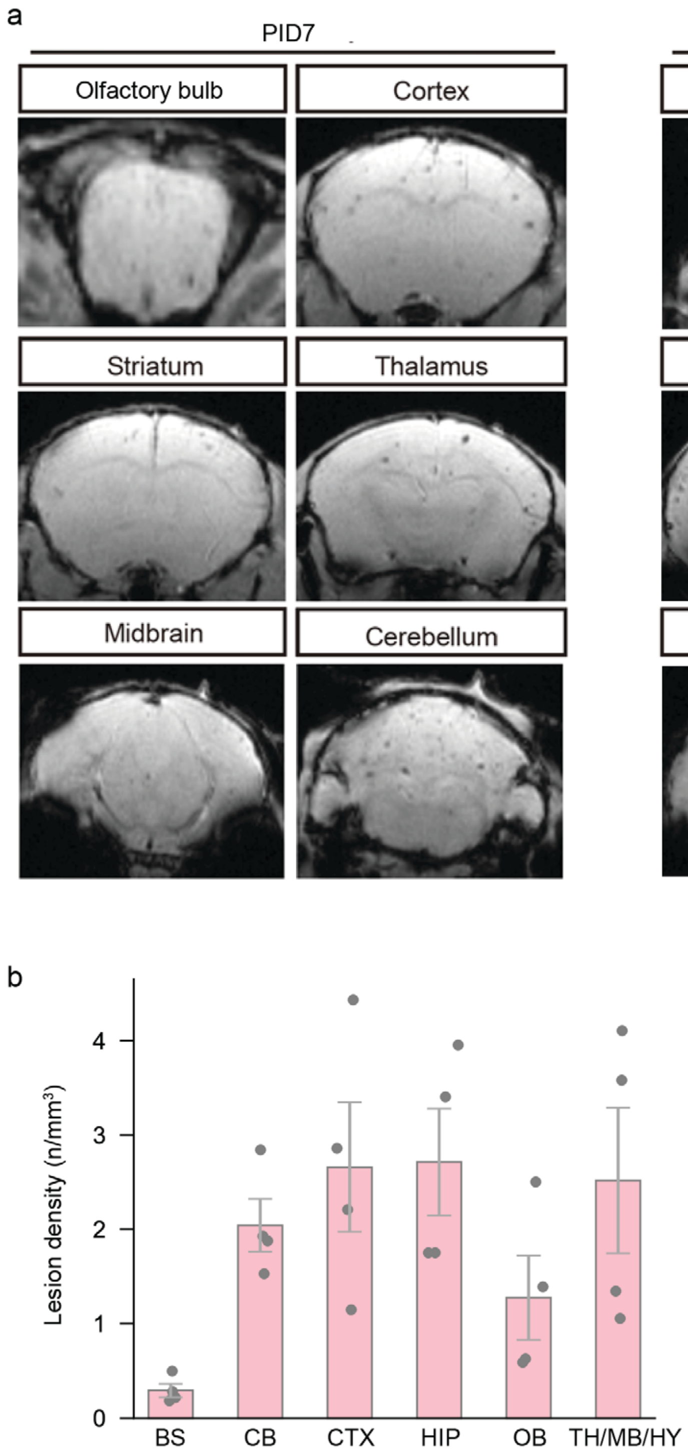

#### Supplementary Figure 29

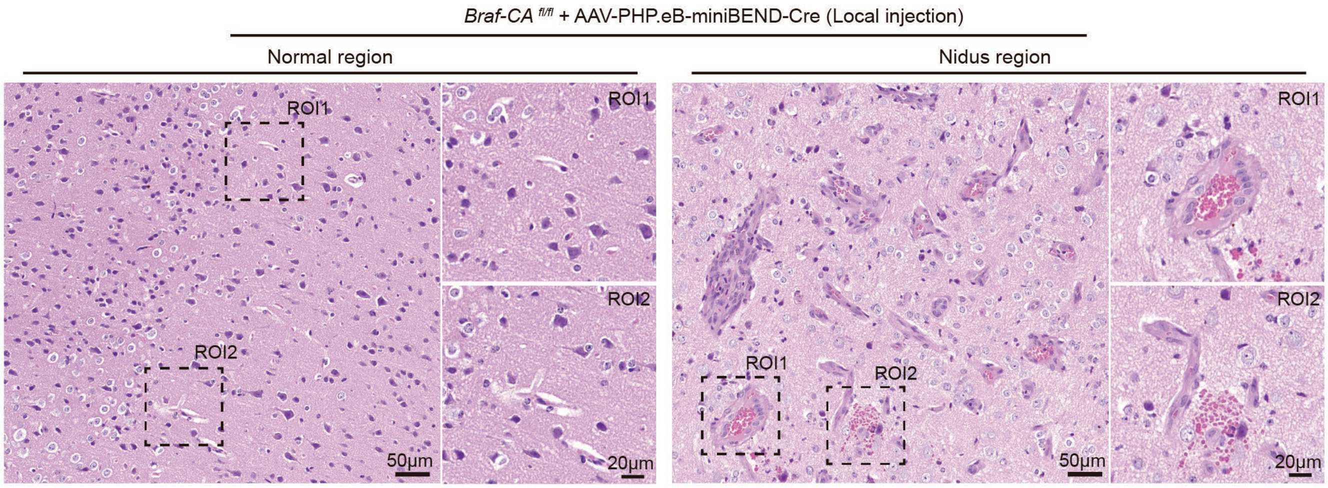

**Supplementary Figure 30**

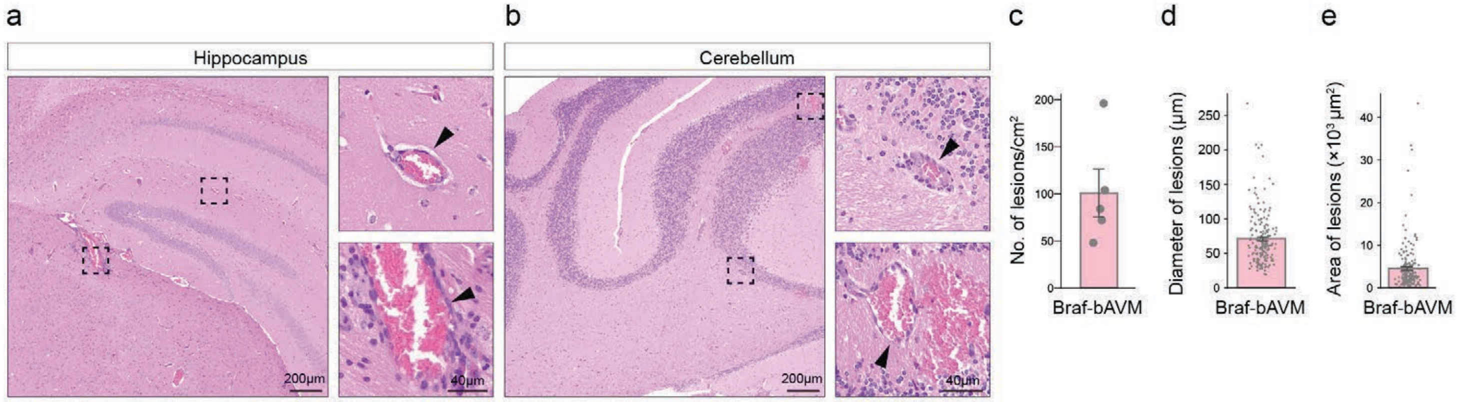

**Supplementary Figure 31**

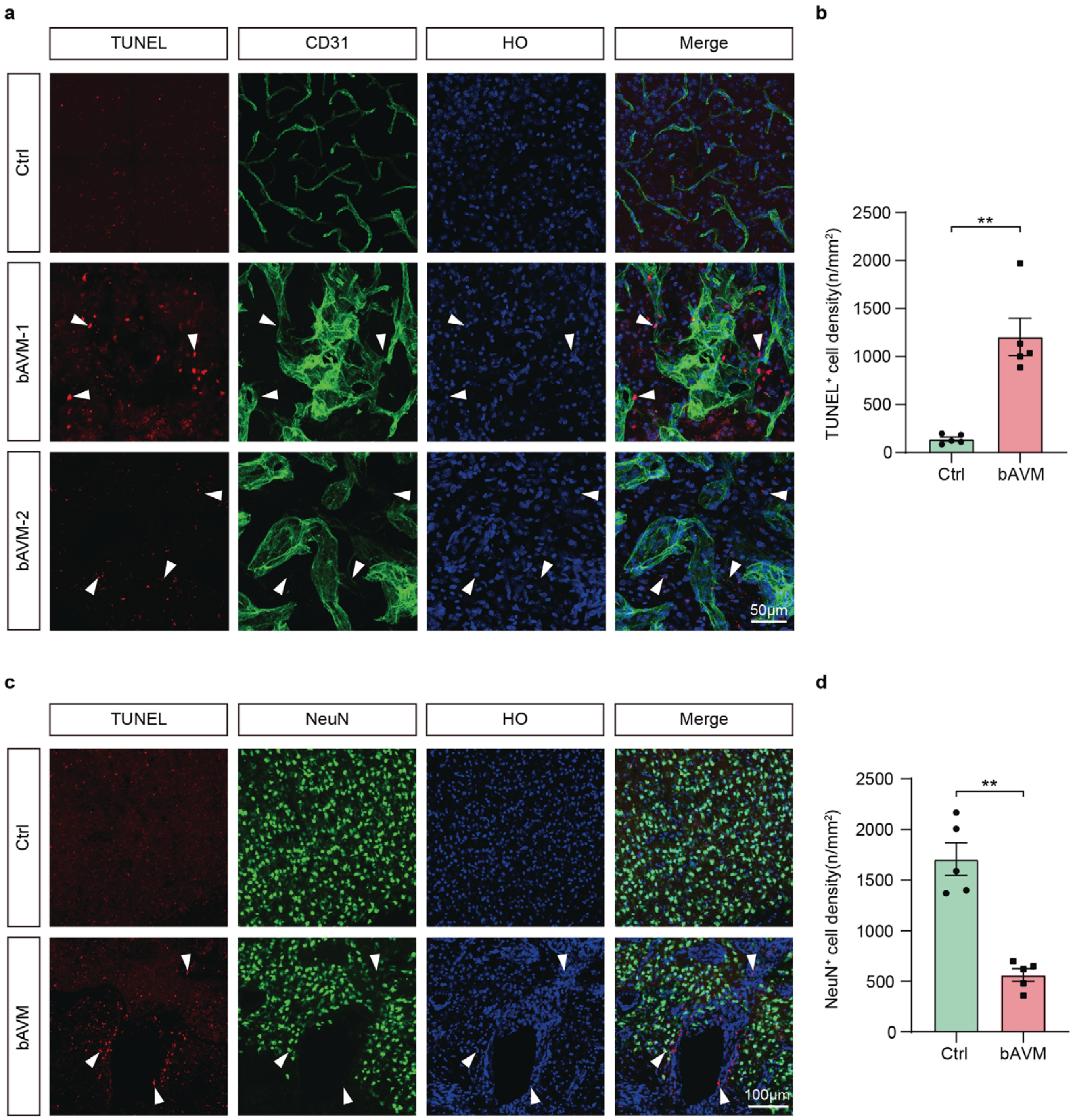

**Supplementary Figure 32**

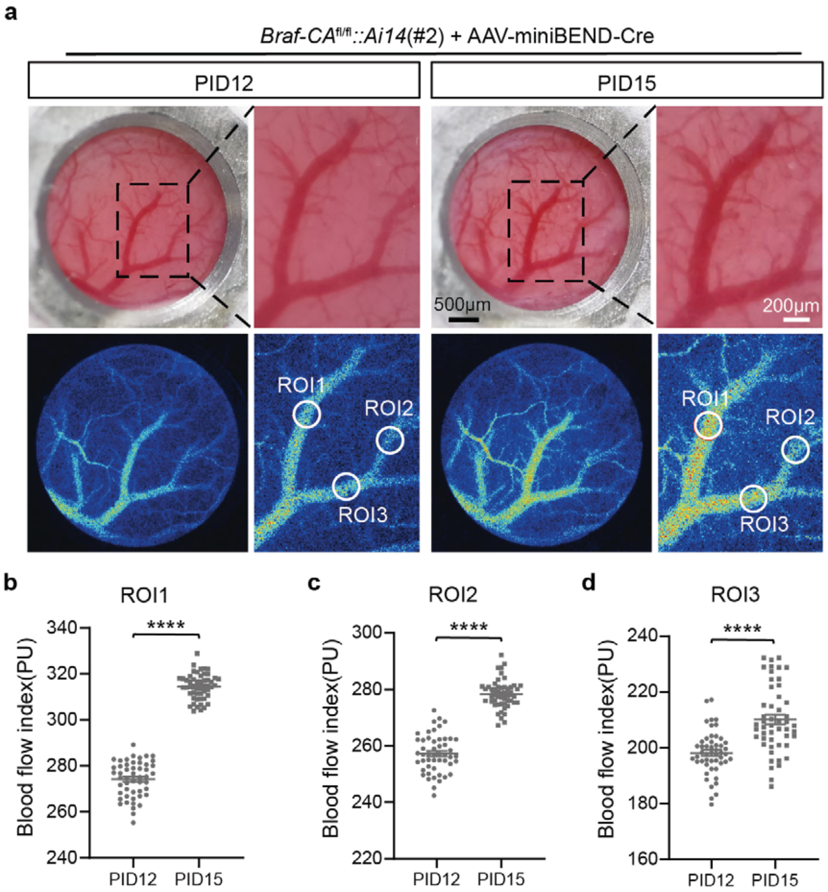

**Supplementary Figure 33**

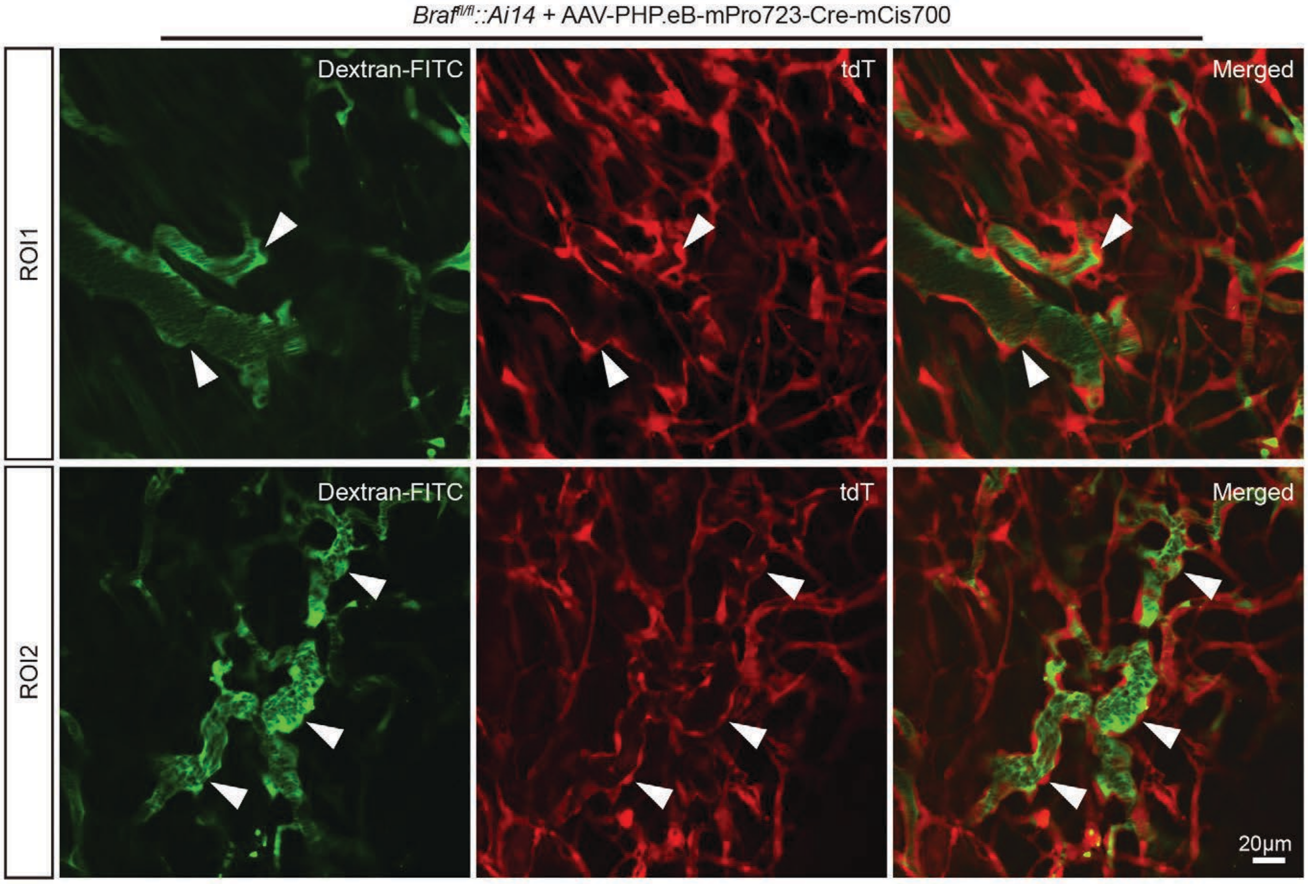

#### Supplementary Figure 34

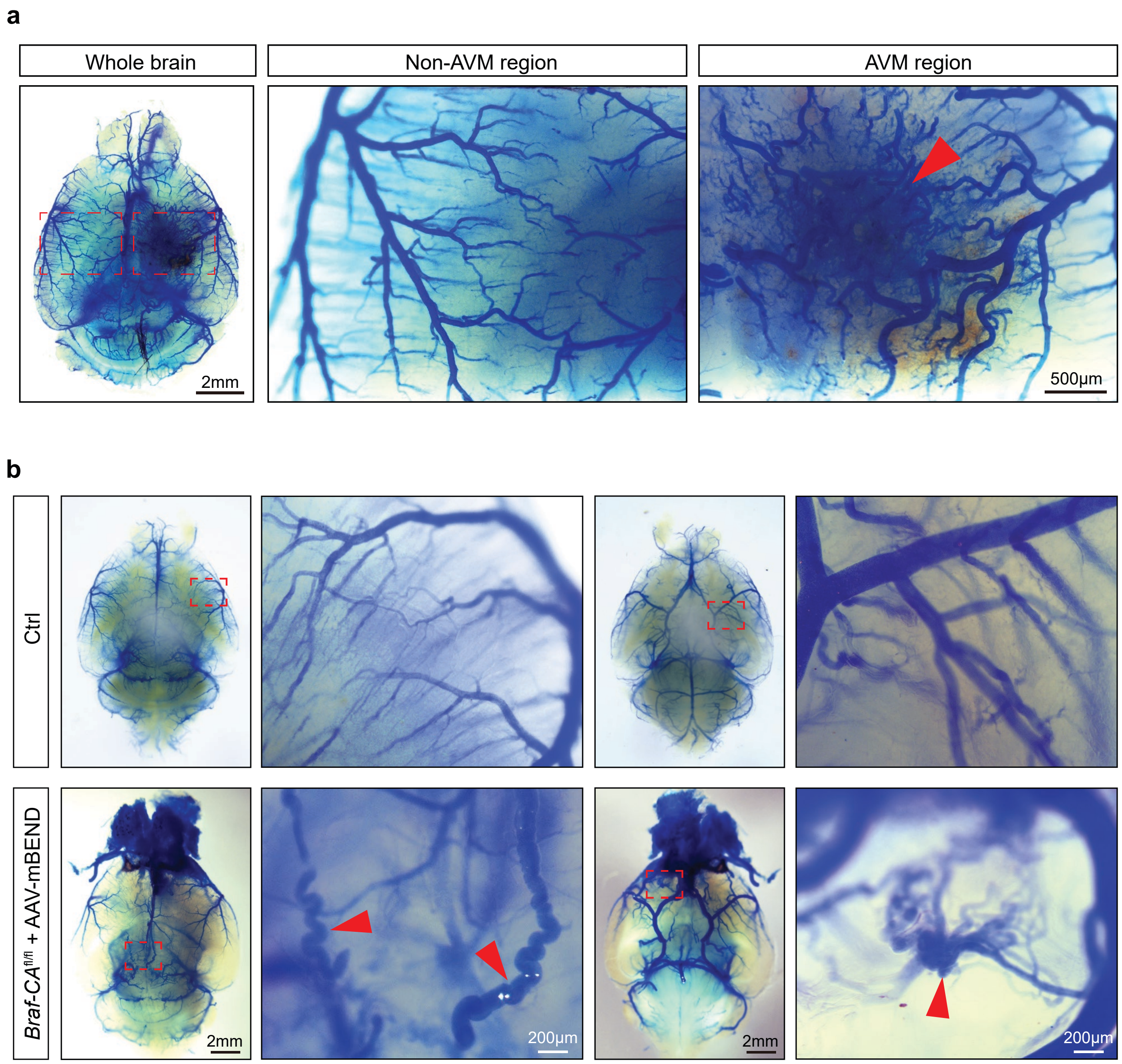

#### Supplementary Figure 35

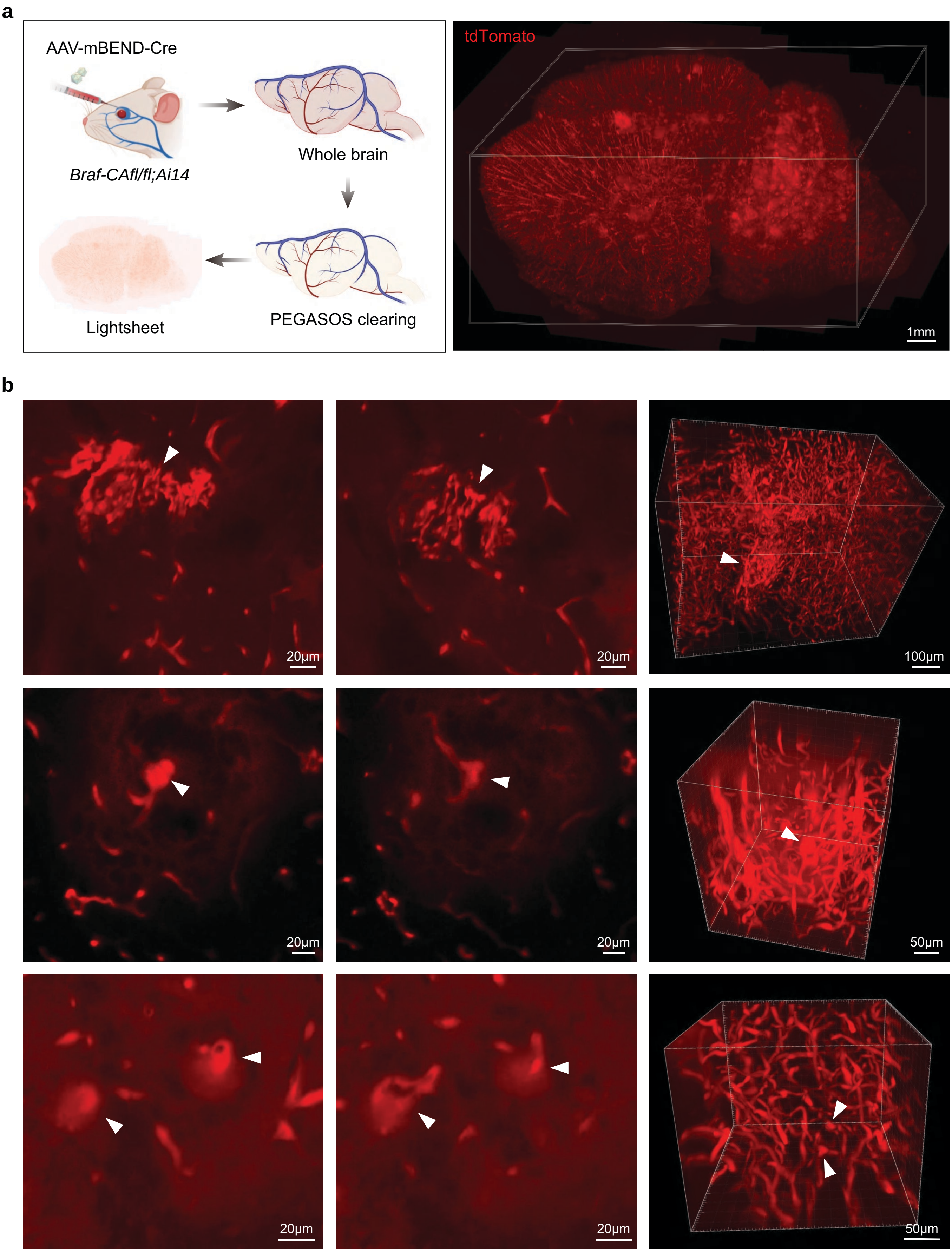

**Supplementary Figure 36**

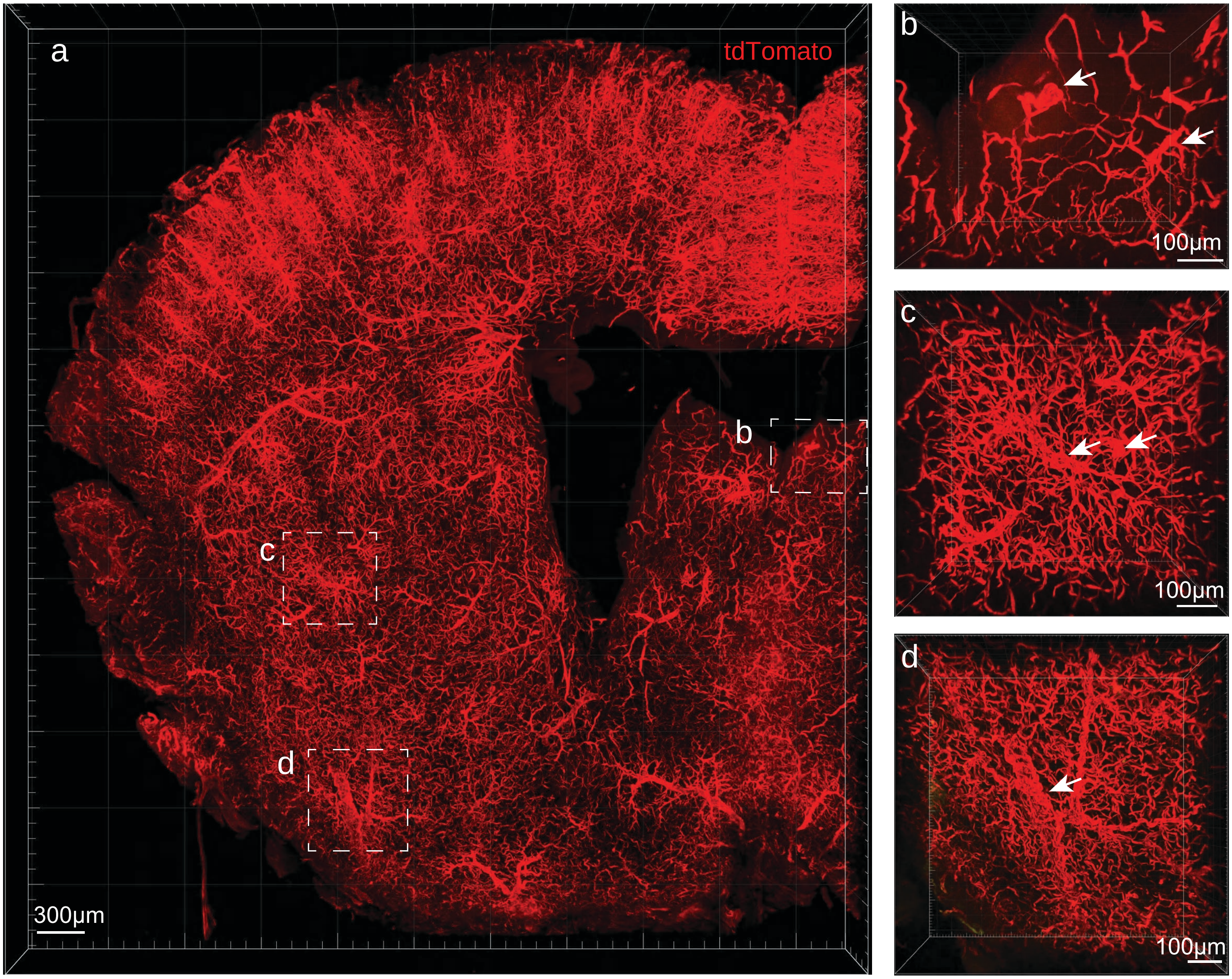

**Supplementary Figure 37**

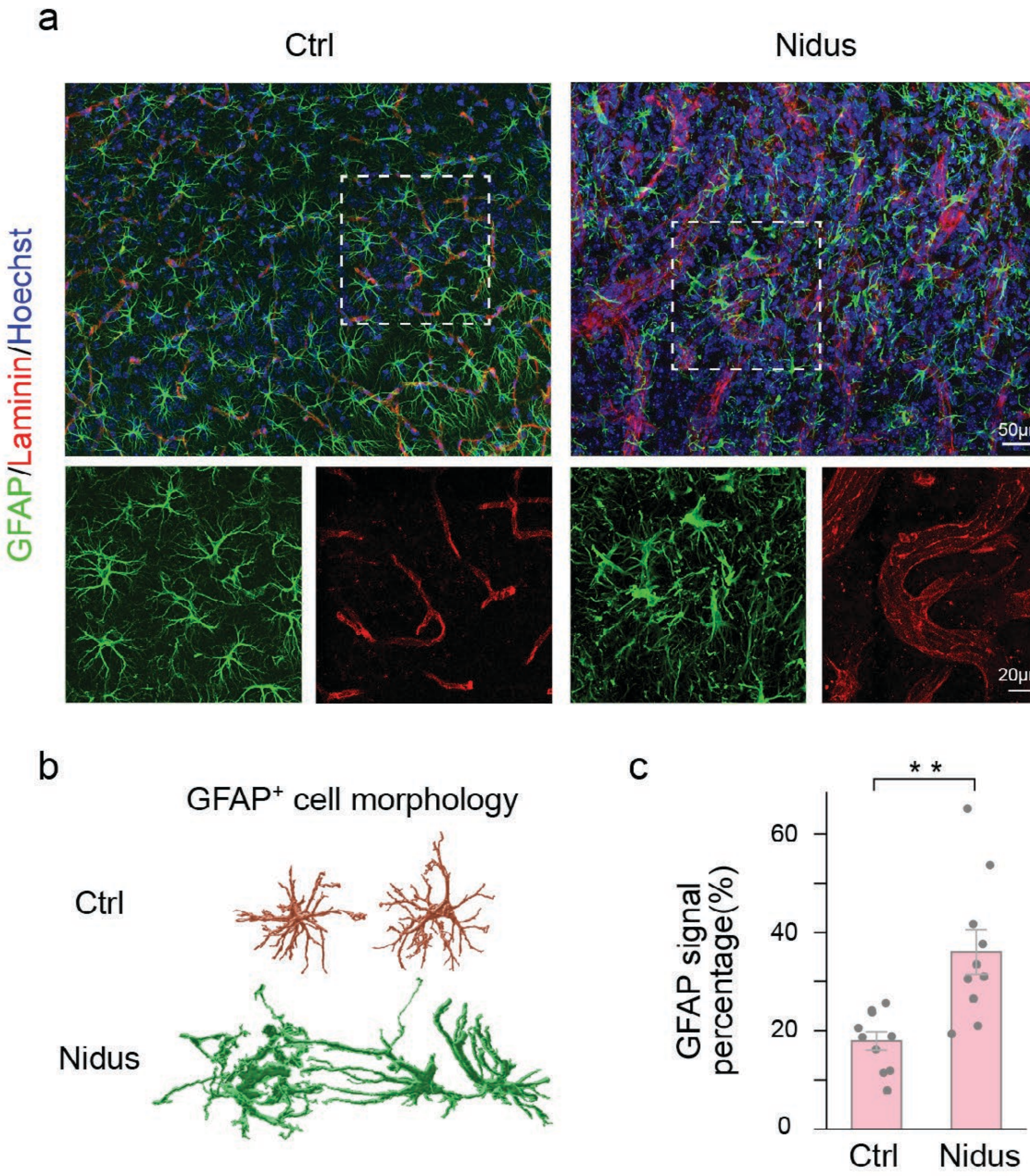

**Supplementary Figure 38**

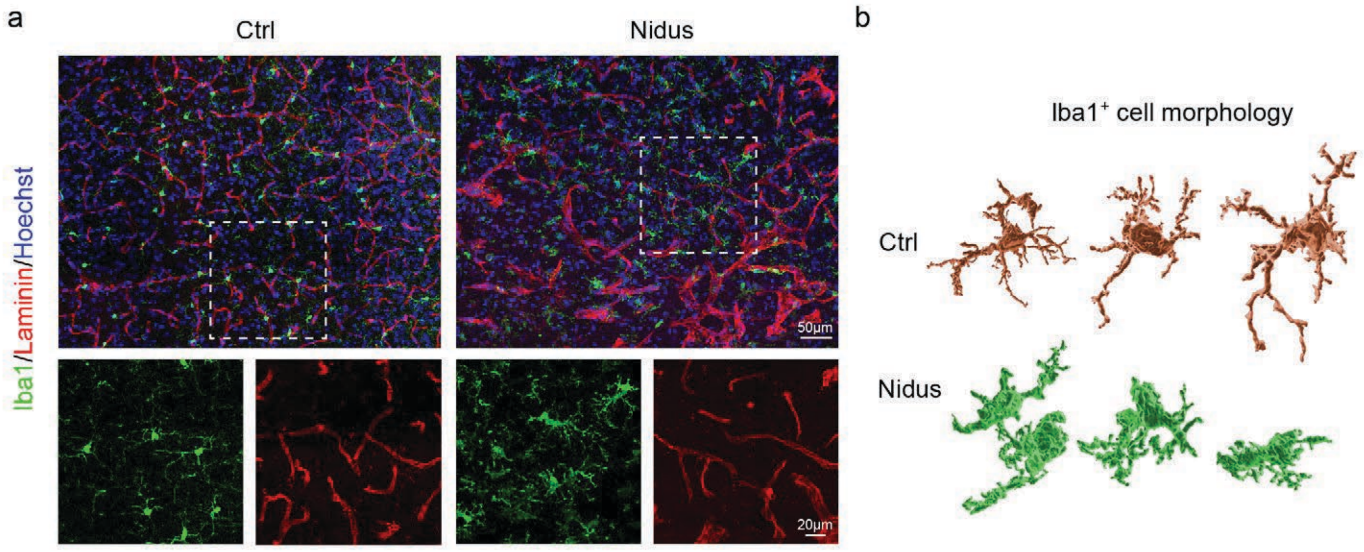

### Supplementary Figure 39

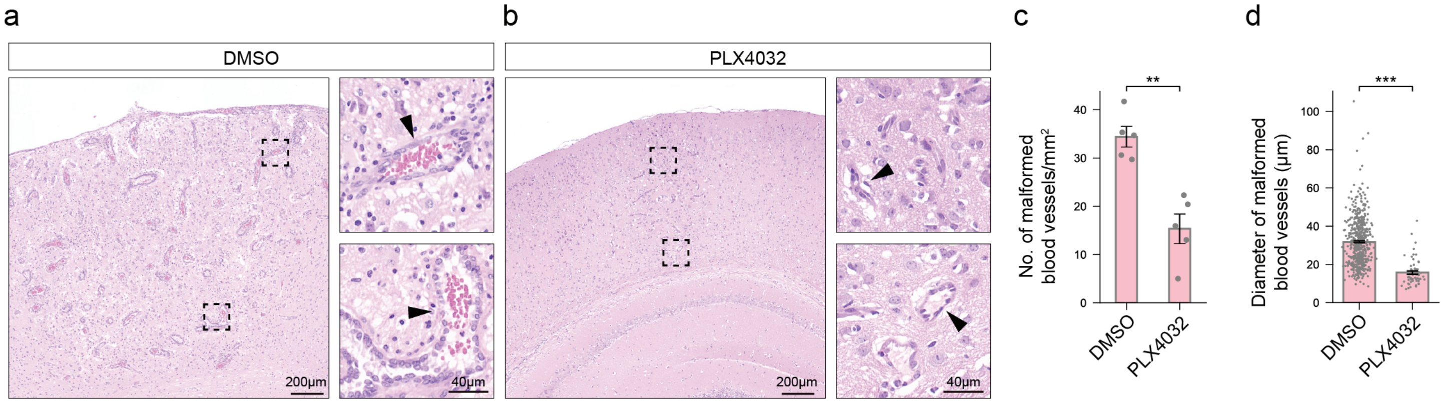

**Supplementary Figure 40**

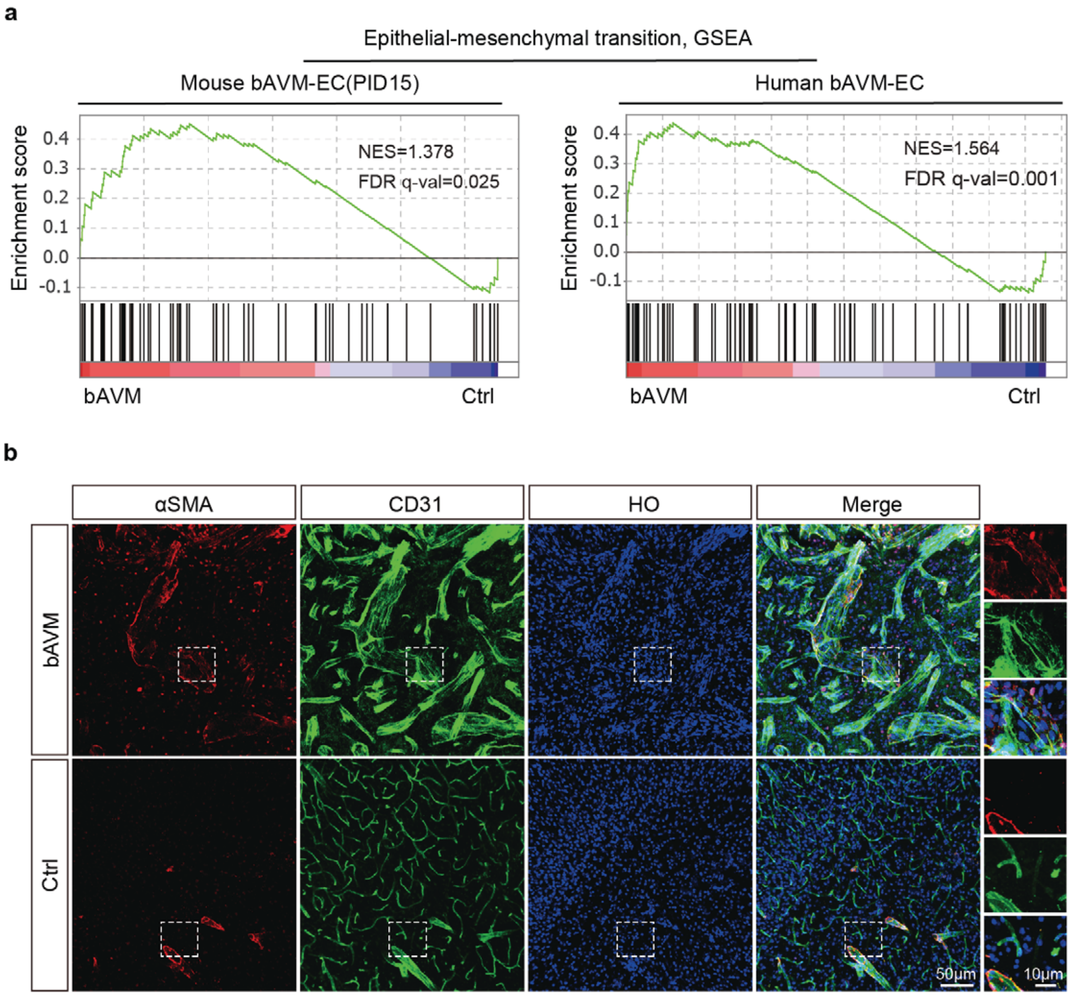
